# Pairing interactions between nucleobases and ligands in aptamer:ligand complexes of riboswitches: Crystal structure analysis, classification, optimal structures and accurate interaction energies

**DOI:** 10.1101/603159

**Authors:** Preethi S. Prabhakar, Purshotam Sharma, Abhijit Mitra

## Abstract

In the present work, sixty-seven crystal structures of the aptamer domains of RNA riboswitches, are chosen for analysis of the structure and strength of hydrogen bonding (pairing) interactions between nucleobases constituting the aptamer binding pockets and the bound ligands. A total of eighty unique base:ligand hydrogen-bonded pairs containing at least two hydrogen bonds were identified through visual inspection. Classification of these contacts in terms of the interacting edge of the aptamer nucleobase revealed that interactions involving the Watson-Crick edge are the most common, followed by the sugar edge of purines and the Hoogsteen edge of uracil. Alternatively, classification in terms of the chemical constitution of the ligand yields five unique classes of base:ligand pairs: base:base, base:amino acid, base:sugar, base:phosphate and base:other. Further, quantum mechanical (QM) geometry optimizations revealed that sixty seven out of eighty pairs exhibit stable geometries and optimal deviations from their macromolecular crystal occurrences. This indicates that these contacts are well-defined RNA aptamer:ligand interaction motifs. QM calculated interaction energies of base:ligand pairs reveal rich hydrogen bonding landscape, ranging from weak interactions (base:other, –3 kcal/mol) to strong (base:phosphate, –48 kcal/mol) contacts. The analysis was further extended to study the biological importance of base:ligand interactions in the binding pocket of the tetrahydrofolate riboswitch and thiamine pyrophosphate riboswitch. Overall, our study helps in understanding the structural and energetic features of base:ligand pairs in riboswitches, which could aid in developing meaningful hypotheses in context of RNA:ligand recognition. This can, in turn contribute towards current efforts to develop antimicrobials that target RNAs.

## INTRODUCTION

Riboswitches are important functional RNA molecules which are generally present in the 5’-untranslated mRNA regions. Riboswitches have the ability to monitor and effectively regulate the expression of fundamental genes associated with the biosynthesis and transport of small metabolites or ligands (1,2). Typically, riboswitches consist of two structural domains: the aptamer domain that recognizes and binds the ligand, and the expression platform that regulates the gene expression in response to ligand-binding (3). Due to their crucial role in regulation of a large number of metabolites, detailed mechanisms of functioning of riboswitches, including cognate ligand recognition and binding; and alteration of transcription and translation events, constitute a major area of research (4–10).

X-ray crystal structures of riboswitch aptamer domains, bound to different ligands, are available at the atomic resolution (1). These crystal structures reveal the significant roles of noncovalent interactions in the ligand-recognition process. For example, electrostatic interactions between the positively charged sulfonium group of S-adenosylmethionine (SAM) and two universally conserved base pairs within the aptamer domain of the SAM-I riboswitch help in discriminating SAM ligand from the structurally similar ligand (i.e. S-adenosylhomocysteine (11)). In addition to a number of other examples (12), recent crystal structures of the guanidinium riboswitches underscore the importance of electrostatic and base:ligand hydrogen bonding interactions in ligand discrimination and binding (13). Further, ligand binding through hydrogen bonding between the ligand and the aptamer nucleobases of purine-sensing riboswitches is responsible for triggering corresponding gene expression (14,15). Furthermore, van der Waals stacking interactions between the bound ligand and the aptamer nucleobases also play an important role in organization of ligand binding pockets in a number of riboswitches (16).

The diversity in noncovalent interactions harnessed during the ligand-binding process as exemplified by riboswitch crystal structures underscore the importance of characterization of the physicochemical forces that regulate riboswitch structure and dynamics. Specifcially, it is necessary to develop and validate hypotheses on how various RNA structural elements within different aptamers, and their respective structurally different cognate ligands, harness unique structural principles in different functional contexts. Since base:ligand hydrogen bonding is one of the most important noncovalent forces that help in ligand recognition by the aptamer domain of riboswitches, statistical analysis of this class of interactions in terms of the identity of the participating aptamer nucleobases and bound ligands constitute the first steps in this direction. Further, quantitative analysis of the structure and strength of these interactions, and careful assessment of their role in the ligand recognition, may contribute to understanding of the overall ligand binding process by riboswitches. Structural bioinformatics analysis of crystal structures of RNA aptamer:ligand complexes, coupled with quantum mechanical (QM) calculations, constitute a comprehensive implementation strategy in this direction. QM calculations are especially useful, since they can provide an accurate description of individual energy components that influence the structural stability of noncovalently bonded chemical entities on one hand, and the intrinsic strengths of noncovalent interactions on the other. Of course, given the differential effects of solvation on different energy components, the results of QM calculations in vacuo are not directly comparable to the binding free energies in solution. Nevertheless, such calculations can accurately quantify the intrinsic strength and stability of hydrogen bonding interactions between aptamer nucleobases and the bound ligands, which may, in turn, help in assessing the contribution of individual pairwise interactions towards the overall strength of binding between the ligand and riboswitches.

Although a substantial number of crystal structure analysis and QM studies have focussed on the analysis of hydrogen bonding interactions that constitute structural motifs present in RNA macromolecules (e.g. nucleobase pairs (17–27), higher order hydrogen bonding contacts including base triplets and quartets (17,28) and their post transcriptionally modified counterparts (27,29), and base-phosphate contacts (30)), very few QM studies have analysed the role of noncovalent interactions in context of specific riboswitches (31,32). Thus, existing literature does not provide a comprehensive understanding of the structural and energetic aspects of aptamer:ligand hydrogen-bonding interactions.

To address these aspects, we have carried out crystal structure database analysis of aptamer-ligand complexes, and have analysed the nature of base:ligand hydrogen-bonding interactions as a function of nucleobase and ligand types, the interacting nucleobase edges, and the nature of H-bonds present within the pair. In addition, with the help of QM calculations, we have obtained optimal geometries of base:ligand pairs and have calculated their accurate binding energies. Overall, our combined analysis of crystal structure dataset and QM geometries and energies provide unique insights into the molecular features of ligand recognition by riboswitches.

## MATERIALS AND METHODS

A dataset of 67 crystal structures (Table 1) spanning the first four of the five recently classified riboswitch categories (33) (namely coenzymes and related compounds, nucleotides and their derivatives, amino acids and sugars, Supplementary Figures S1–S4), (containing base:ligand pairs with at least two hydrogen bonds, was selected from protein data bank (PDB), on the basis of visual analysis, for preliminary analysis. This dataset excludes crystal structures in which ions act as ligands, mainly because the associated aptamer:ligand noncovalent contacts are physiochemically different from the (hydrogen bonding) interactions considered in the present work. The dataset was also curated to include only structures having at least 3.2 Å resolution, and by retaining only wild type aptamer crystal structures. The final comprehensive dataset contained structures of 38 aptamers bound to natural (biological) ligands and 29 aptamers bound to ligand derivatives.

**Table 1.**
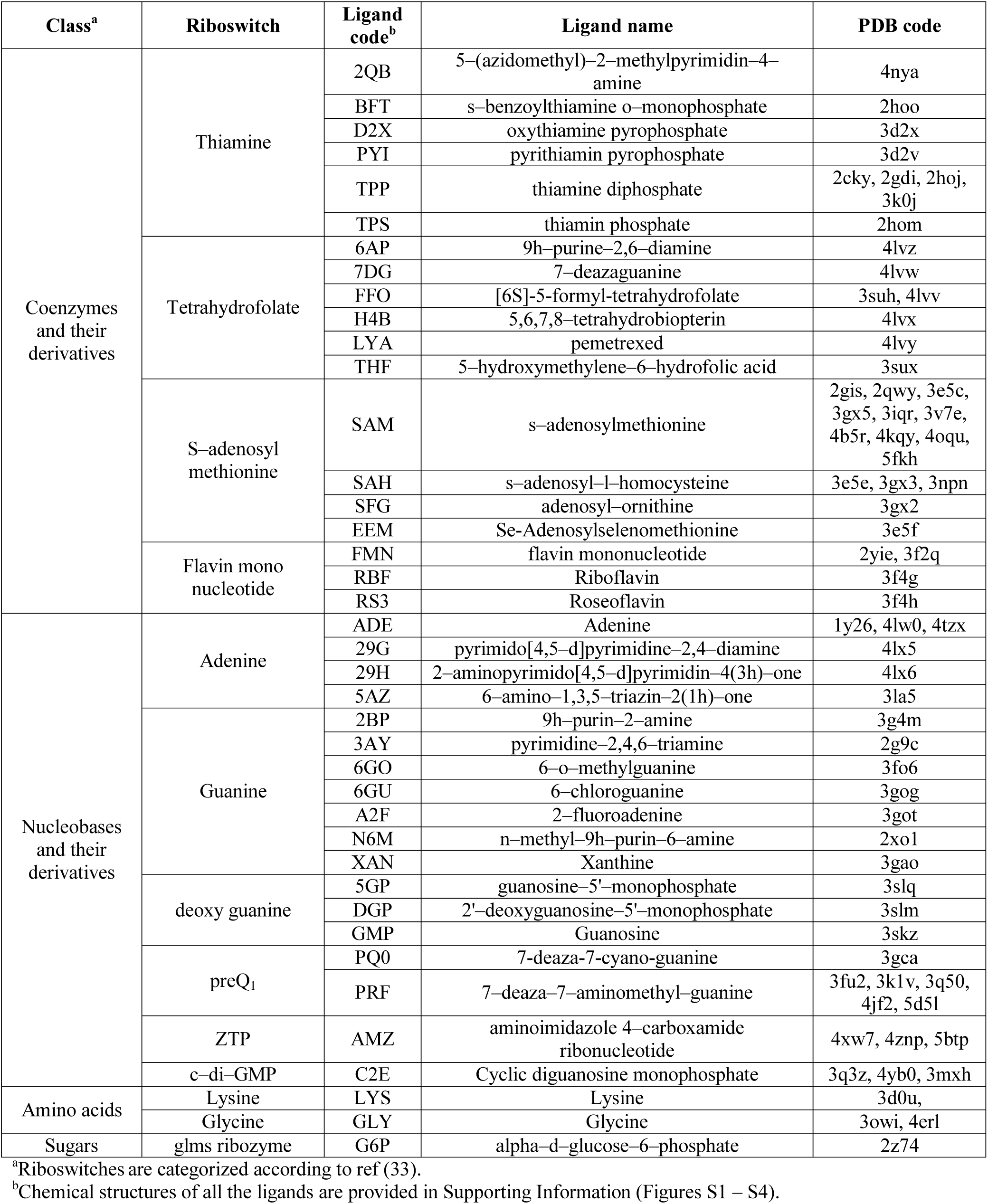
PDB accession codes of the 67 riboswitch:ligand complexes studied in the present work. The names and three-letter codes of 40 different bound ligands are provided.

A total of 80 base:ligand pairs connected through two hydrogen bonds were extracted from these crystal structures, statistically analysed and used as initial models for QM analysis. For QM geometry optimizations, the missing hydrogen atoms were added to the heavy-atom coordinates of the ligands, extracted from the respective crystal structures, to complete the necessary covalency requirements in the initial models. Subsequently, the interacting aptamer nucleotides were truncated at their glycosidic bonds, and the C1’ atoms were replaced with hydrogen atoms, wherever the ribose sugars of these nucleotides do not interact with bound ligands (20). In cases where the ribose sugar bound to the aptamer nucleobase interacts with ligands, the respective sugars were retained after replacing the 5′-phosphate group with a hydroxyl group, in order to retain a tractable, but relevant model size. Further, in case of ligands with amino acid functionality, the -NH_2_ and CO_2_H were taken as neutral and for ligands with phosphate group interactions (D2X, PYI and G6P), the phosphate group was considered as a charged moiety. Geometry optimization of the base:ligand pairs was carried out at the B3LYP/6-31G(d,p) level using Gaussian 09 (34), which was selected in synchrony with previous studies on RNA base pairs (18,19,21-24).

To understand the difference in base:ligand geometry, in its crystal occurrence and in the corresponding fully optimized form, the root mean square deviations (rmsd) was calculated using VMD v1.9 software (35). Further, the strength of each base:ligand pair was measured in terms of interaction energy, which is a measure of the stabilization acquired by the interacting system through hydrogen bonding. Basis set superposition error (BSSE) corrected interaction energies were calculated using the counterpoise method (36) at the RIMP2/aug-cc-pVDZ or MP2/aug-cc-pVDZ level (37,38) using Turbomole v6.2 (39) or Gaussian 09 (34), which was selected in analogy with previous studies on RNA base pairs (20,22-24,27,40,41). Although geometries of 67 out of total 80 base:ligand pairs could be fully optimized, remaining 13 pairs either showed highly deviated optimized structures compared to the initial structures or could not be optimized (Table 2). In these cases, as in previous studies on RNA base pairs (19), constrained optimizations, restricted to optimization of the positions of the added hydrogen atoms, were performed by freezing all the heavy atom coordinates. Interaction energies were calculated on these constrained-optimized systems at the RIMP2/aug-cc-pVDZ or MP2/aug-cc-pVDZ level.

**Table 2.**
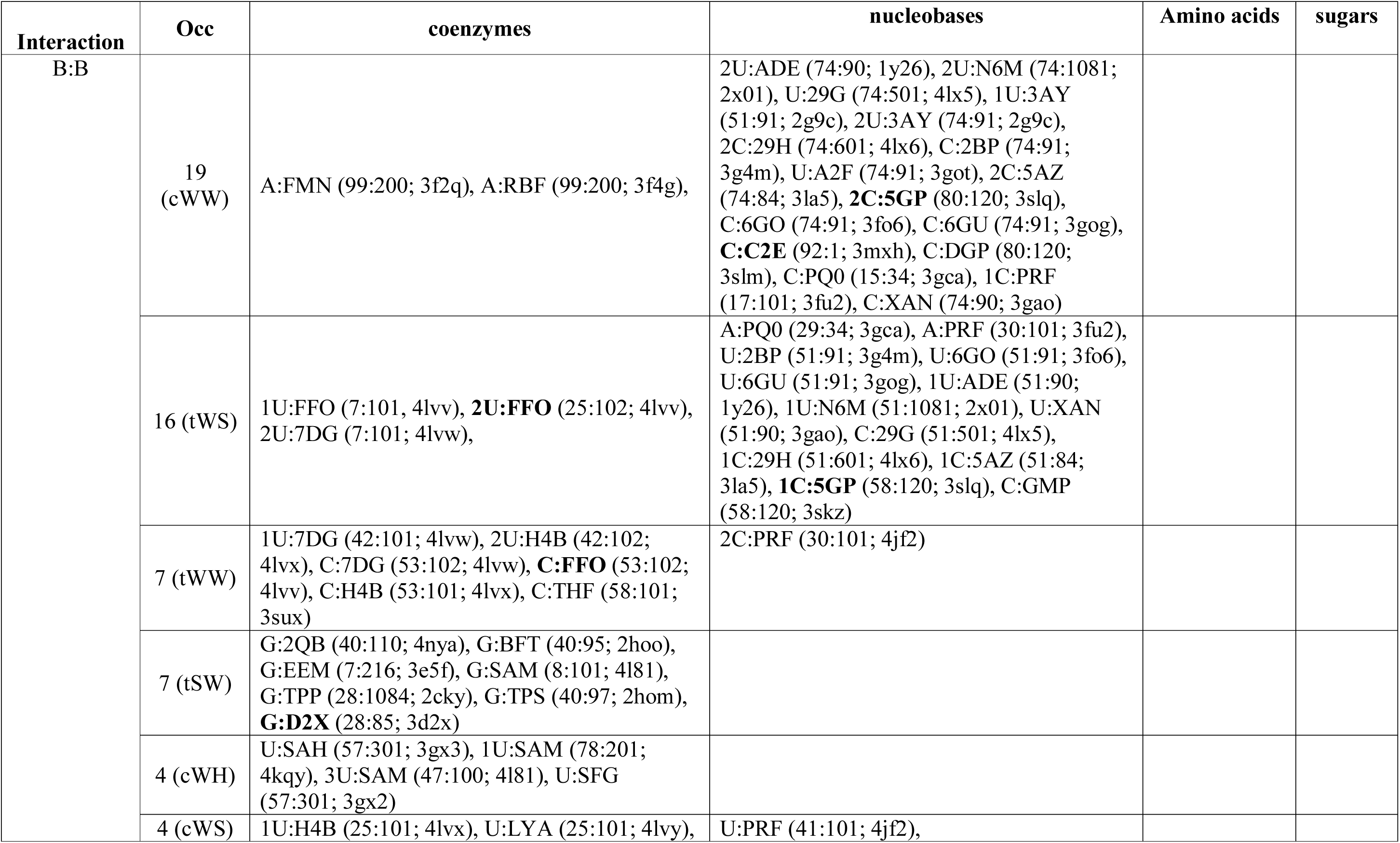
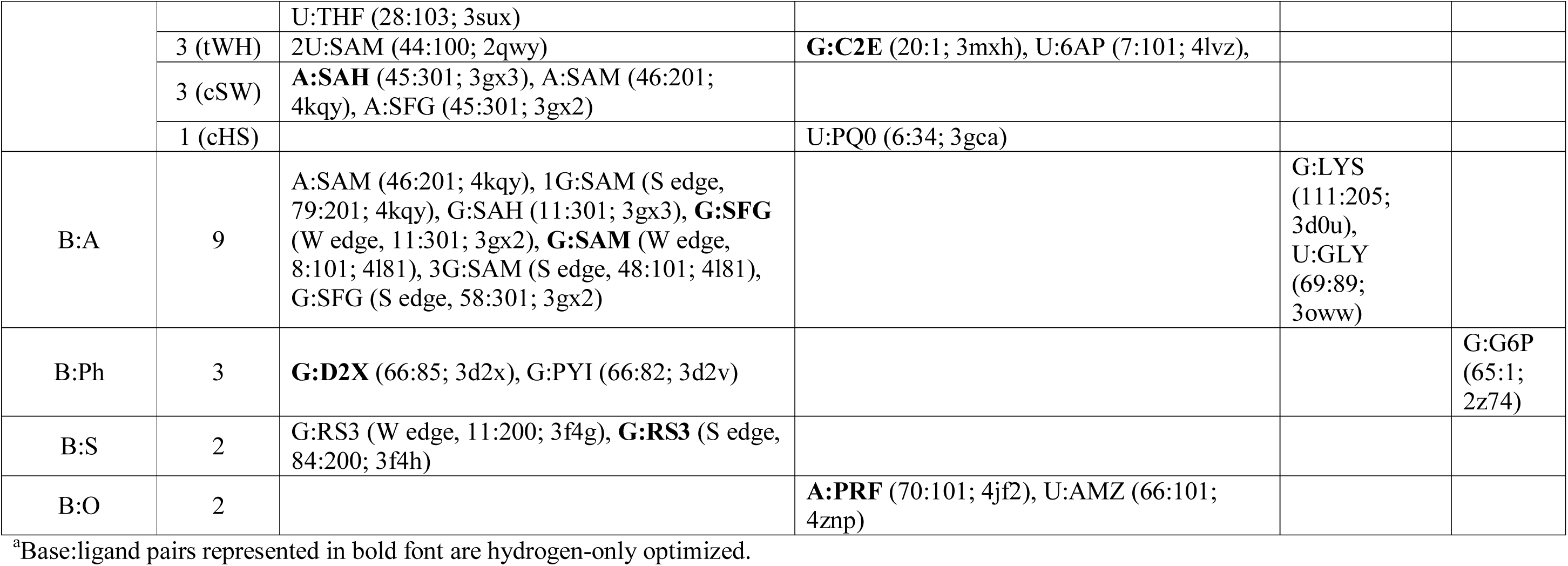
Distribution of base:ligand interactions in the riboswitch aptamers with respect to base pairing families.^a^

| Interaction | Occ | coenzymes | nucleobases | Amino acids | sugars |
| --- | --- | --- | --- | --- | --- |
| B:B | 19 (cWW) | A:FMN (99:200; 3f2q), A:RBF (99:200; 3f4g), | 2U:ADE (74:90; 1y26), 2U:N6M (74:1081; 2x01), U:29G (74:501; 4lx5), 1U:3AY (51:91; 2g9c), 2U:3AY (74:91; 2g9c), 2C:29H (74:601; 4lx6), C:2BP (74:91; 3g4m), U:A2F (74:91; 3got), 2C:5AZ (74:84; 3la5), **2C:5GP** (80:120; 3slq), C:6GO (74:91; 3fo6), C:6GU (74:91; 3gog), **C:C2E** (92:1; 3mxh), C:DGP (80:120; 3slm), C:PQ0 (15:34; 3gca), 1C:PRF (17:101; 3fu2), C:XAN (74:90; 3gao) |  |  |
|  | 16 (tWS) | 1U:FFO (7:101; 4lvv), **2U:FFO** (25:102; 4lvv), 2U:7DG (7:101; 4lvw), | A:PQ0 (29:34; 3gca), A:PRF (30:101; 3fu2), U:2BP (51:91; 3g4m), U:6GO (51:91; 3fo6), U:6GU (51:91; 3gog), 1U:ADE (51:90; 1y26), 1U:N6M (51:1081; 2x01), U:XAN (51:90; 3gao), C:29G (51:501; 4lx5), 1C:29H (51:601; 4lx6), 1C:5AZ (51:84; 3la5), **1C:5GP** (58:120; 3slq), C:GMP (58:120; 3skz) |  |  |
|  | 7 (tWW) | 1U:7DG (42:101; 4lvw), 2U:H4B (42:102; 4lvx), C:7DG (53:102; 4lvw), **C:FFO** (53:102; 4lvv), C:H4B (53:101; 4lvx), C:THF (58:101; 3sux) | 2C:PRF (30:101; 4jf2) |  |  |
|  | 7 (tSW) | G:2QB (40:110; 4nya), G:BFT (40:95; 2hoo), G:EEM (7:216; 3e5f), G:SAM (8:101; 4l81), G:TPP (28:1084; 2cky), G:TPS (40:97; 2hom), **G:D2X** (28:85; 3d2x) |  |  |  |
|  | 4 (cWH) | U:SAH (57:301; 3gx3), 1U:SAM (78:201; 4kqy), 3U:SAM (47:100; 4l81), U:SFG (57:301; 3gx2) |  |  |  |
|  | 4 (cWS) | 1U:H4B (25:101; 4lvx), U:LYA (25:101; 4lvy), | U:PRF (41:101; 4jf2), |  |  |
|  |  | U:THF (28:103; 3sux) |  |  |  |
|  | 3 (tWH) | 2U:SAM (44:100; 2qwy) | **G:C2E** (20:1; 3mxh), U:6AP (7:101; 4lvz), |  |  |
|  | 3 (cSW) | **A:SAH** (45:301; 3gx3), A:SAM (46:201; 4kqy), A:SFG (45:301; 3gx2) |  |  |  |
|  | 1 (cHS) |  | U:PQ0 (6:34; 3gca) |  |  |
| B:A | 9 | A:SAM (46:201; 4kqy), 1G:SAM (S edge, 79:201; 4kqy), G:SAH (11:301; 3gx3), **G:SFG** (W edge, 11:301; 3gx2), **G:SAM** (W edge, 8:101; 4l81), 3G:SAM (S edge, 48:101; 4l81), G:SFG (S edge, 58:301; 3gx2) |  | G:LYS (111:205; 3d0u), U:GLY (69:89; 3oww) |  |
| B:Ph | 3 | **G:D2X** (66:85; 3d2x), G:PYI (66:82; 3d2v) |  |  | G:G6P (65:1; 2z74) |
| B:S | 2 | G:RS3 (W edge, 11:200; 3f4g), **G:RS3** (S edge, 84:200; 3f4h) |  |  |  |
| B:O | 2 |  | **A:PRF** (70:101; 4jf2), U:AMZ (66:101; 4znp) |  |  |

<sup>a</sup>Base:ligand pairs represented in bold font are hydrogen-only optimized.

## RESULTS

### Statistical overview of base:ligand interactions from crystal structural occurrences

In the dataset of chosen crystal structures (Table 1) the size of the bound ligands varies considerably, ranging from small chemical entities (e.g. glycine and lysine) to complex chemical structures (e.g. folate-related ligands, S-adenosyl methionine and thiamine pyrophosphate derivatives, Supplementary Figures S1–S4). The total 80 unique base:ligand interactions, thus identified, can be further classified based on the interacting edge (i.e. Watson-Crick edge (W edge), Hoogsteen edge (H edge) or sugar edge (S edge) of the aptamer nucleobase (adenine (A), cytosine (C), guanine (G) or uracil (U)). It is observed that 78% of these interactions involve the nucleobase W edge (7% A, 9% G, 36% U and 26% C), followed by the S edge of purines (21% total (i.e. 5% A and 16% G)) and the H edge of U (1%, Figure 1C and Table 2). Thus, pyrimidines in the binding pocket of aptamers interact with the ligands, almost exclusively, through their W edge. On the other hand, the purines mainly use the W edge or S edge, where the highest binding occurrence is shown by U (37%), followed by C (26%). A close third is G (25%) using its S edge (16%) and W edge (9%) in the ratio of around 5:3, whereas the least frequent is A (12%) using its S edge (5%) and W edge (7%) in an approximately 1:1 ratio. In contrast to multiple examples of W-edge and S-edge interactions, we could find only a single occurrence of H-edge interaction, where U interacts with PQ0 in the aptamer domain of the preQ_1_ riboswitch (PDB code: 3gca).

**Figure 1.**
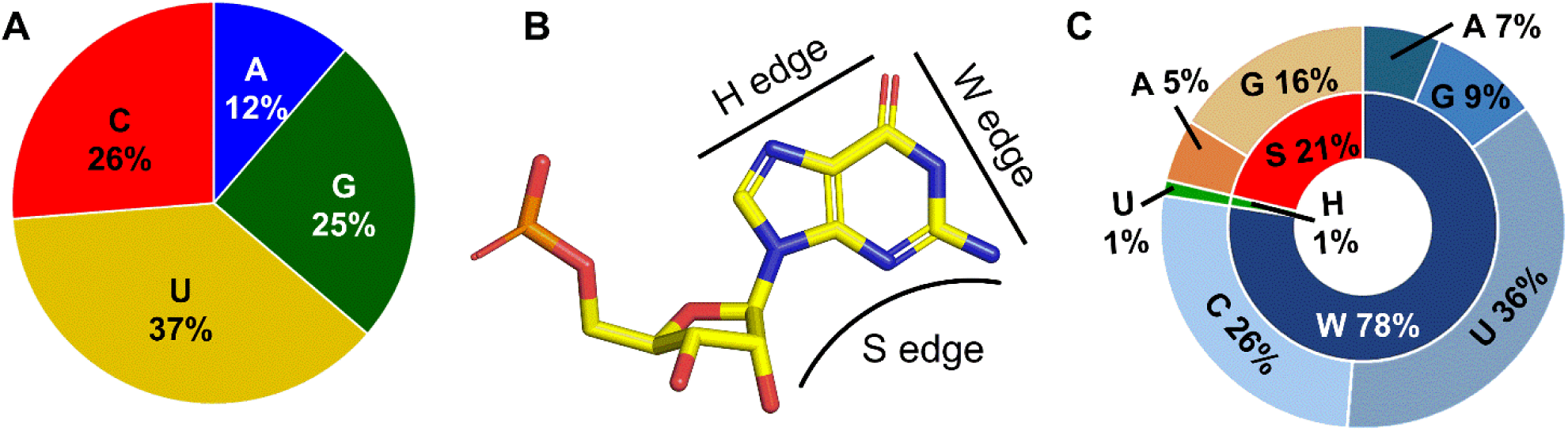
(A) Statistical distribution of the aptamer nucleobases that interact with the ligand in riboswitches in the crystal structures. (B) Interacting edges (Watson-Crick edge (W edge); Hoogsteen edge (H edge) and Sugar edge (S edge)) of the aptamer nucleobases. (C) Distribution of base:ligand pairs as a function of interacting edge (inner circle) and the aptamer base (outer circle).

### Classification of base:ligand interactions

Classification of base:ligand pairs on the basis of the chemical composition of the ligand, as well as on the nature of the ligand-aptamer interactions at the molecular level, reveals five basic interaction types: base:base (B:B), base:amino acid (B:A), base:sugar (B:S), base:phosphate (B:Ph) and base:other (B:O, Figure 2 and Table 2). Although, B:B (80%) and B:A (11%) interactions have significant occurrences, instances of the other three types of interactions (B:S (2 instances), B:Ph (3 instances) and B:O (2 instances)) constitute only the remaining 8% of occurrences. Interestingly, the aptamer nucleobase component of all three B:Ph interactions involve the W edge of the aptamer base, whereas B:S and B:O interactions involve either W edge or S edge of the base.

**Figure 2.**
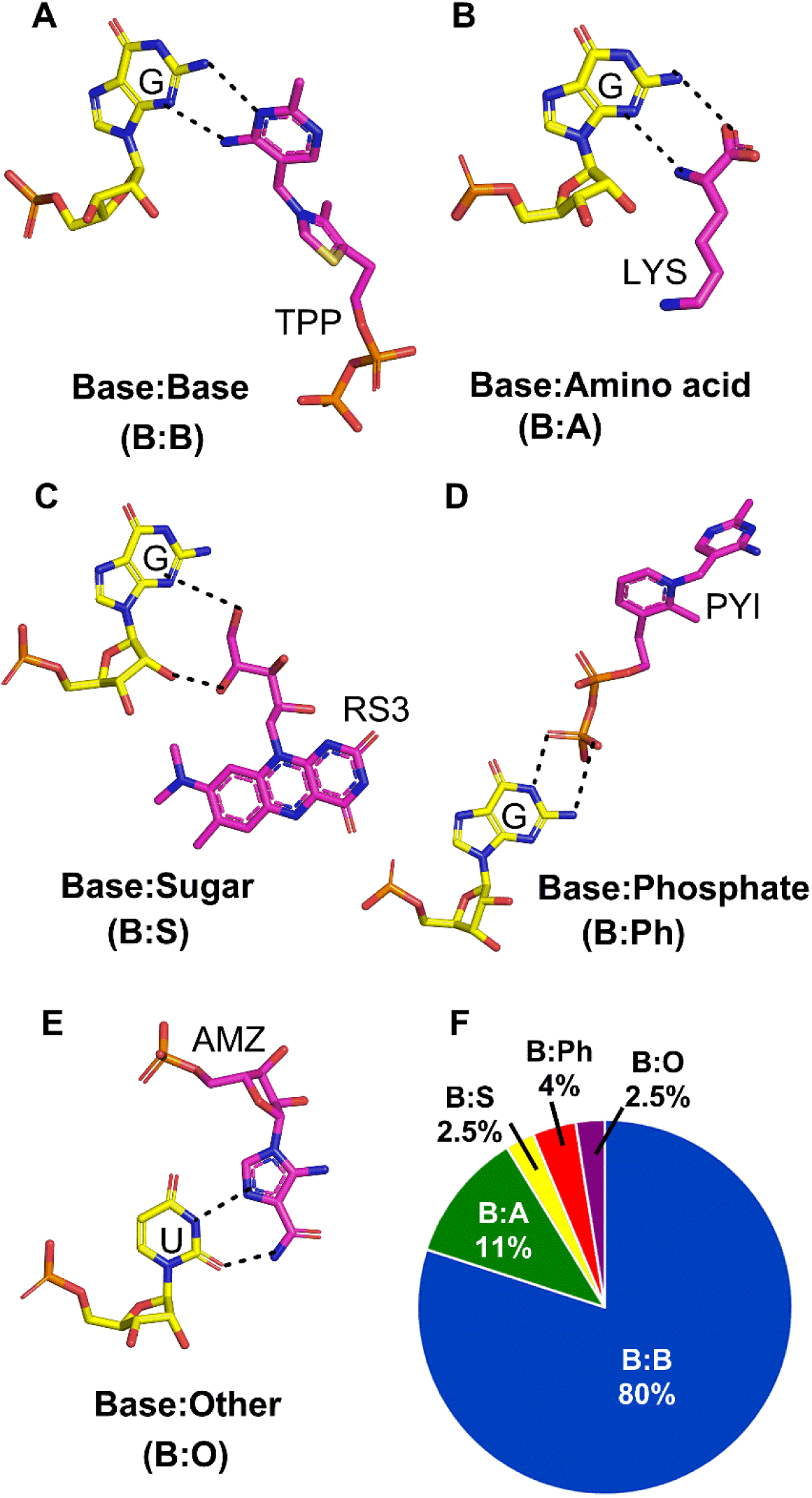
(A-E) Examples of each base:ligand interaction type (base:base (B:B), base:amino acid (B:A), base:sugar (B:S), base:phosphate (B:Ph) and base:other (B:O) observed in the crystal structures of ligand-bound aptamer domains of riboswitches. (F) Statistical distribution of different base:ligand interactions in the crystal structures.

The most abundant (B:B) contacts involve ligands possessing partial or complete structural similarity with the canonical RNA nucleobases. Consequently, the pairing geometries of aptamer-ligand B:B interactions are comparable with the base pairing geometries observed in folded RNA structures (42). Notably, about 30% of these interactions adopt cWW base pairing, followed by tWS (25%), tWW (11%), cWS (6%), cWH (6%) and tWH (6%, Table 2) pairings (42). All these interactions involve the W edge of the aptamer nucleobase and constitute 84% of the B:B interactions. In contrast, only two pairing geometries involve the S edge of the aptamer nucleobase (cSW, 5% and tSW, 11%). The single interaction involving the H edge of the aptamer nucleobase forms cHS pairing. Overall, the observed B:B interactions span 7 (c/tWW, c/tWH, c/tWS, and cHS) of the 12 possible RNA base pairing geometries.

In contrast, B:A interactions (Figure 2B) span 9 instances of base:amino acid contacts, and can be further classified into two interaction types: 2 interactions involving a free amino acid and 7 contacts where the interacting amino acid moiety is a part of the more complex chemical structure of the ligand (Supplementary Figures S1 and S4). Both the interaction types involve the W edge or S edge of aptamer nucleobases. On the other hand, B:S, B:Ph or B:O interactions with very few available examples, involve the interaction of aptamer nucleobase with the sugar, phosphate or side chain moieties of the ligands.

### QM calculations

QM geometry optimization of base:ligand pairs, isolated from their respective crystal environments, is expected to reveal their intrinsic structural features. This may in turn, provide insights into the contribution of base:ligand pairing to the overall stabilization of ligand within the aptamer binding pocket. Accordingly, geometry optimizations, followed by interaction energy calculations were attempted on all base:ligand pairs (Supplementary Table S1). Although the hydrogen bonding patterns of 8 B:B, 2 B:A, and 1 each of B:Ph, B:S and B:O pairs, either could not be optimized or deviated significantly from their crystal structures on full optimization (Table 2), all the remaining pairs retained their respective hydrogen bonding patterns on optimization.

**Figure 3.**
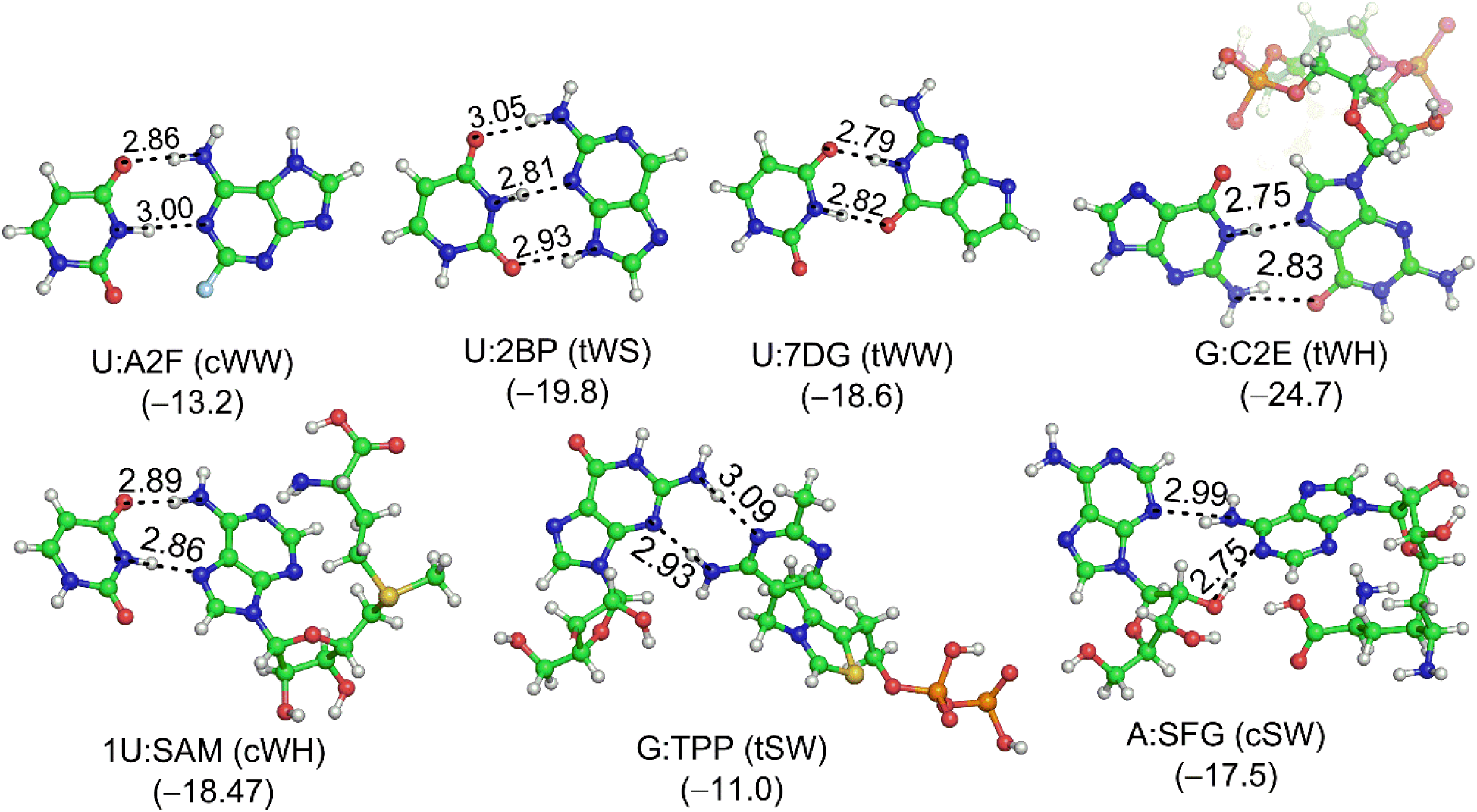
Representative B3LYP/6-31G(d,p) optimized geometries and MP2/aug-cc-pVDZ interaction energies (parentheses, kcal/mol) of base:base (B:B) type of base:ligand interactions. The hydrogen bonding donor–acceptor (D–A) distances (Å) are provided against each hydrogen bond (dotted lines).

#### B:B pairs

Among the B:B interactions, all 53 pairs that interact through W edge of the aptamer nucleobase show linear N-H⋅⋅⋅N/O or O-H⋅⋅⋅N H-bonds on optimization, indicating strong base pairing characteristics (Supplementary Figure S5-S8, Supplementary Tables S2–S7), 19 of these interactions possess a minimum of three strong (N-H⋅⋅⋅N/O or O-H⋅⋅⋅N type) hydrogen bonds. Except for complex ligands with flexible functional groups (e. g. SAM and its derivatives, TPP, FMN and THF), all the W-edge B:B pairs showed relatively small deviations from their respective crystal structure geometries on optimization (rmsd ranging from 0.1 Å ≤ 1 Å), which points towards their significance as well defined base:ligand interaction motifs. However, in case of complex ligands, flexible functional groups (e. g., thiazole and pyrophosphate groups of TPP derivatives, sugar moiety of FMN derivatives, p-amino benzoate and glutamate of THF derivatives) the structural deviation increased upon optimization, resulting in higher rmsd (1.2 Å – 2.2 Å), albeit without affecting the base:ligand hydrogen bonding. Regardless, in all cases, interaction energy calculations reveal substantial (–14 to –32 kcal/mol) strength of W-edge B:B pairing (Supplementary table S1).

With the exception of three pairs (G:TPS, G:EEM and A:SFG), the structural deviation on optimization of all the S-edge B:B interactions (rmsd of 1.0 Å – 2.2 Å) is high (Supplementary Table S1). This occurs partly due to the inherent torsional flexibility associated with ribose, which falls to the nearest local minima on optimization. Further, except for three interactions mediated by at least three H-bonds (G:2QB, G:EEM and 2G:SAM), all other S-edge pairs possess only two hydrogen bonds on optimization (Supplementary Figures S6-S8 and Supplementary Tables S8-S9). Albeit smaller than for W-edge pairs, the strengths of S-edge B:B base pairs is evidenced by the calculated interaction energies (–11 to –18 kcal/mol, Supplementary Table S1).

#### B:A pairs

For the B:A interactions, optimizations reveal three distinct hydrogen bonding patterns (Figure 4, Supplementary Tables S11). The first pattern includes interaction of both amino (–NH_2_) and carboxylic groups (–CO_2_H) of the amino acid moiety of the ligand. In this type, the amino group acts a donor, whereas the carboxylic oxygen atoms either act as a donor or acceptor, thus stabilizing each pair with a minimum of two strong H-bonds. Optimized structures of five of the total nine B:A pairs exhibit this interaction pattern, which spans a wide range (–8 to –40 kcal/mol) of interaction strength. The second pattern observed only in one (G:SFG) pair, simultaneously harnesses both the donor as well as acceptor properties of the amino nitrogen of the amino acid and possesses significant (–23 kcal/mol) interaction energy. Specifically, the - NH_2_ group of the ornithine moiety of SFG acts as a H-bond donor (N-H(SFG)⋅⋅⋅N3(G)) as well as acceptor (N2-H(G)⋅⋅⋅N(SFG)) in the optimized G:SFG pair, and. In contrast, the third interaction pattern observed in three (G:SAH, G:SAM and G:SFG) B:A pairs involves the acceptor interaction of the –CO_2_H group of the amino acid moiety with the W edge of G. However, relatively small magnitude of interaction strength (–8 to –18 kcal/mol) suggest the inherent weakness of this hydrogen-bonding pattern compared to the other two patterns (Supplementary Table S1). Regardless, the B:A pairs span a relatively smaller range of rmsd (0.9 Å – 1.5 Å) compared to B:B pairs (0.1 Å – 2.2 Å, Supplementary Table S1).

**Figure 4.**
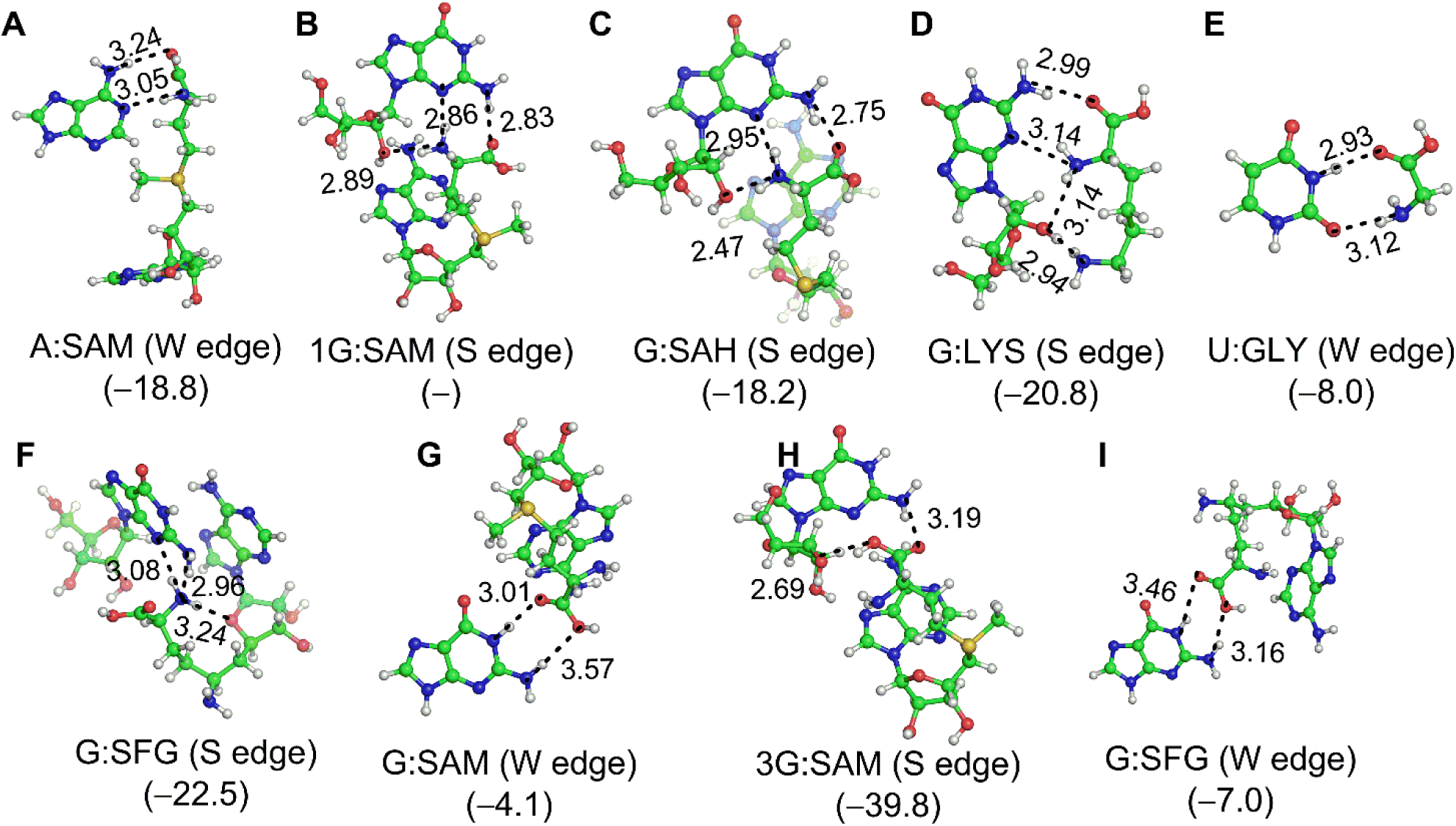
B3LYP/6-31G(d,p) optimized geometries and MP2/aug-cc-pVDZ interaction energies (parentheses, kcal/mol) of base:amino acid (B:A) type of base:ligand pairs. (A–E) B:A pairs involving interaction of aptamer nucleobase with both –NH_2_ and –CO_2_H groups of the amino acid. (F) B:A pair involving interaction of aptamer nucleobase with only –NH_2_ groups of the amino acid. (G-I) B:A pairs involving interaction of aptamer nucleobase with only –CO_2_H groups of the amino acid. The hydrogen bonding donor–acceptor (D–A) distances (Å) are provided against each hydrogen bond (dotted lines). For structures G:SAM (W edge) and G:SFG (W edge), only the positions of hydrogen atoms were optimized, whereas full optimization was carried out for all other structures.

#### B:Ph pairs

Among the B:Ph interactions, except for G:D2X pair which could not be fully-optimized, the other two pairs have more than two hydrogen bonds. For example, G:G6P is stabilized by three hydrogen bonds, whereas four acceptor-bifurcated hydrogen bonds are observed for the G:PYI pair. Regardless, interaction energies (–31 kcal/mol and –48 kcal/mol) of the fully-optimized pairs reveal that B:Ph pairs are stronger than all other categories of base:ligand pairs (Figure 5, Supplementary Table S11).

**Figure 5.**
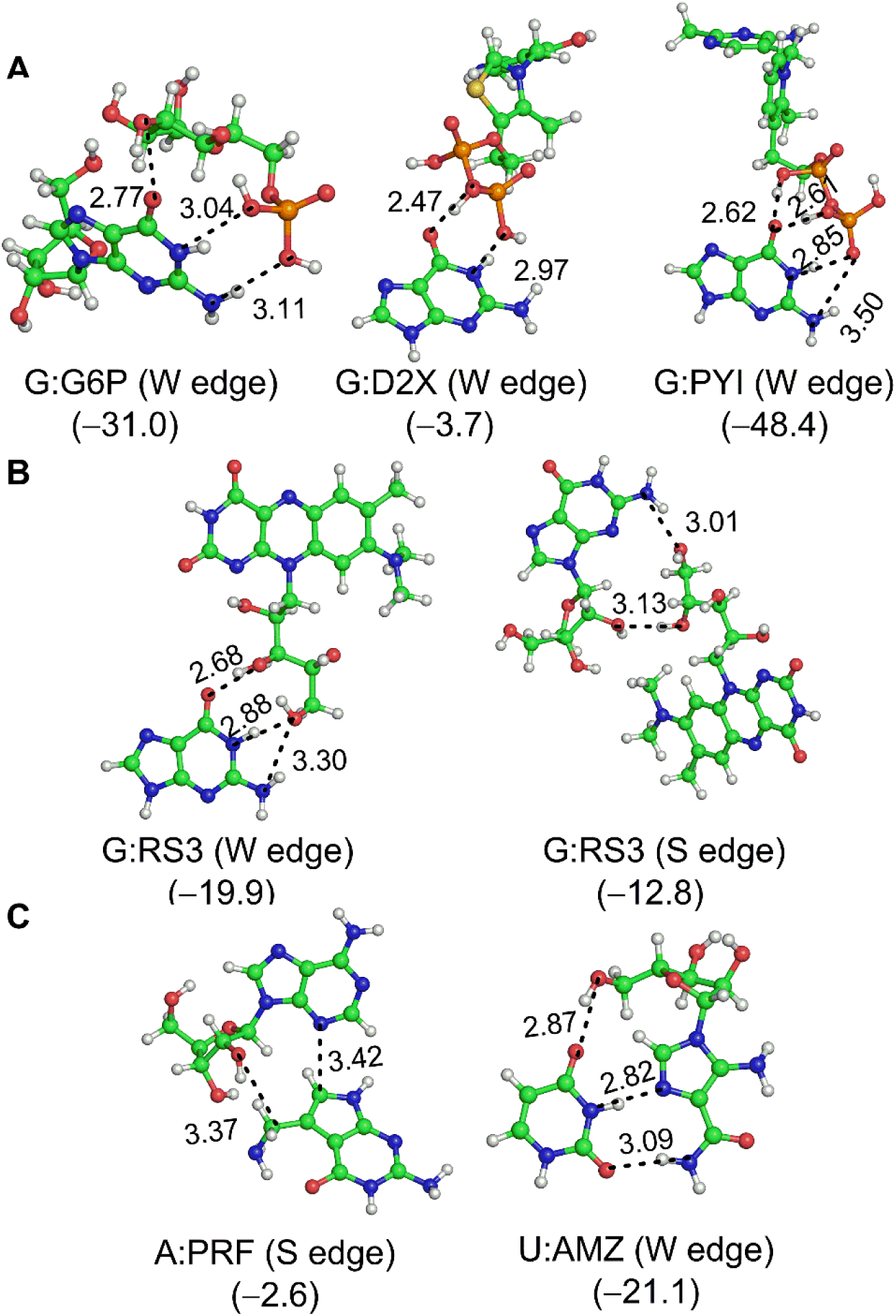
B3LYP/6-31G(d,p) optimized geometries and MP2/aug-cc-pVDZ interaction energies (parentheses, kcal/mol) of (A) base:phosphate (B:Ph), (B) base:sugar (B:S) and (C) base:other (B:O) types of base:ligand pairs. For structures G:D2X (W edge), G:RS3 (S edge) and A:PRF (S edge), only the positions of hydrogen atoms were optimized, whereas full optimization was carried out for all other structures. The hydrogen bonding donor–acceptor (D–A) distances (Å) are provided against each hydrogen bond (dotted lines).

#### B:S pairs

In context of B:S interactions, despite the presence of sugar moiety in the chemical structure of all flavin-related ligands, only RS3 shows significant (i.e. involving at least two hydrogen bonds) interaction, where the sugar group of RS3 interacts with W edge or S edge of G. While the optimized geometry of the W-edge pair is similar to the crystal geometry (rmsd of 0.7 Å) and is stabilized by a substantial (–20 kcal/mol) binding energy, the optimized S-edge pair shows relatively weaker (–13 kcal/mol) binding. Further, interaction energies of B:S pairs are similar to the S-edge interactions of B:B pairs (Figure 5, Supplementary Table S11).

#### B:Ph pairs

Among the two B:O pairs, the A:PRF pair involves a weak (–3 kcal/mol) interaction of side chain of PRF with the S-edge of adenine, which is mediated by two C-H⋅⋅⋅N/O H-bonds. Consequently, this structure shows a high rmsd (2.1 Å) on optimization. However, the second (U:AMZ) pair involves the strong interaction between imadazole group of AMZ ligand and W edge of U, and is mediated by three (two N-H⋅⋅⋅N/O and one O-H⋅⋅⋅O) hydrogen bonds. Thus, this pair exhibits small rmsd (0.7 Å) and high (–21 kcal/mol) interaction energy on optimization (Figure 5 and Supplementary Table S11).

### Biological relevance of base:ligand pairs: Case studies and insights

#### 1. Alternate conformation of Thiamine Pyrophosphate (TPP) aptamer induced by 2QB ligand

The chemical structure of the TPP ligand is composed of three distinct chemical moieties-pyrimidine, thiazole and pyrophosphate. The pyrimidine and pyrophosphate moieties are recognized by the pyrimidine domain and the pyrophosphate sensor domain of the aptamer respectively (Figure 6A (43)). However, since thiazole moiety is not specifically recognized, ligands similar to TPP, but lacking the thiazole moiety (e.g. PYI) may also bind with the aptamer, and consequently trigger riboswitch action (43). Among 11 crystal structures of TPP aptamer analyzed in the present study, only six (2QB, BFT, D2X, PYI, TPP and TPS) interact with the G in the pyrimidine sensor domain to form B:B pairs mediated by two H-bonds with the following order of strength: G:2QB (–16 kcal/mol) > G:D2X (–14 kcal/mol) > G:BFT (–13 kcal/mol) > G:TPP (–11 kcal/mol) = G:TPS (–11 kcal/mol). However, despite strongest B:B (i.e. G:2QB) pairing, 2QB induces a different conformation of the aptamer, compared to binding with other ligands (44). Analysis of B:Ph interactions between the ligands and the pyrophosphate domain reveals that due to absence of the pyrophosphate group in its chemical structure, 2QB misses out on a very significant B:Ph interaction, whereas other ligands form strong B:Ph contacts. For example, D2X forms B:B contacts with G28 and G11 of the pyrimidine sensor element and additional B:Ph contacts with G66 of the pyrophosphate sensor element (PDB code: 3d2x). Thus, the absence of crucial B:Ph interactions explains the overall weaker binding of the 2QB. Overall, this example illustrates the importance of different interacting moieties in the chemical structure of the ligand in optimal base:ligand binding.

**Figure 6.**
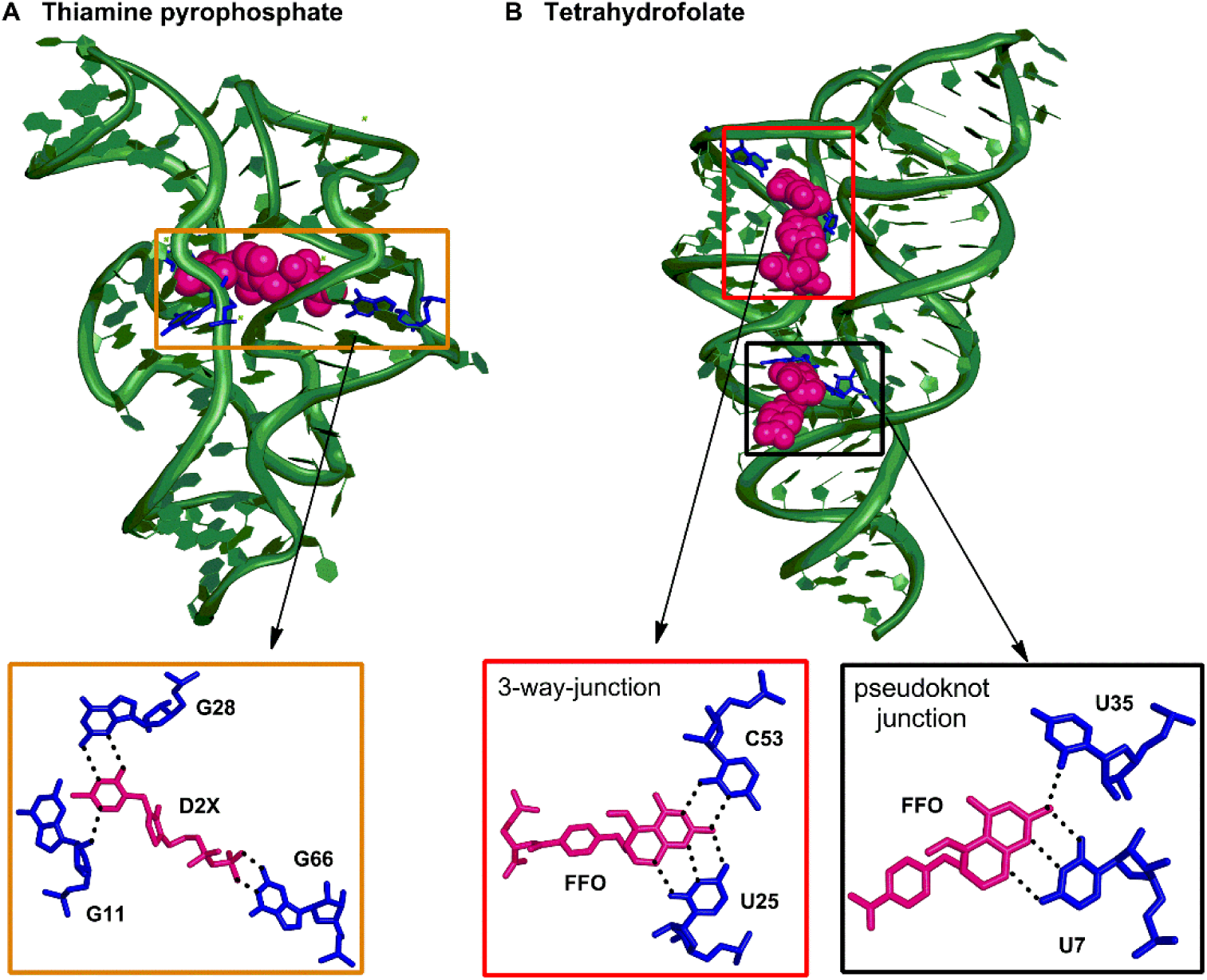
(A) D2X ligand bound to the aptamer domain of thiamine pyrophosphate riboswitch. (B) FFO ligand bound to the aptamer domain of tetrahydrofolate riboswitch.

#### 2. Importance of ligand chemical structure and allosteric binding pockets in ligand binding with the aptamer domain of THF riboswitch

The aptamer domain of THF riboswitch binds folate-related ligands that are composed of three structural elements-pterin ring, p-aminobenzoate (pABA) group and glutamate group (Figure 6B). Although pterin ring plays the most important role in molecular recognition by forming B:B interactions with the aptamer, the pABA region also interacts with the aptamer bases either through hydrogen bonding or stacking interactions. However, apart from facilitating proper ligand orientation, the glutamate group does not directly participate in noncovalent interactions with the aptamer domain.

Within the studied crystal structures, the aptamer domain of THF riboswitch is bound to six different ligands (6AP, 7DG, FFO, H4B, LYA and THF). These ligands either contain all three structural moieties of THF (FFO, LYA or THF) or contain tailored purine bases similar to pterin ring of THF (6AP, 7DG or H4B). Our QM calculations reveal that the observed order of stability of B:B pairs involving the pterin ring of the ligand is 7DG (–43 kcal/mol) > FFO (–34 kcal/mol) > THF (–31 kcal/mol) = H4B (–31 kcal/mol). However, although the B:B pair involving 7DG possesses highest interaction energy, the aptamer possesses overall low affinity towards this pterin analogue (45), mainly due to absence of pABA moiety, which stabilizes the ligand through stacking interactions with the aptamer domain. This further reiterates the synergy in interactions involving different chemical moieties and the aptamer in determining the overall ligand:aptamer affinities.

The aptamer domain of THF riboswitch can simultaneously bind two ligands through two different binding pockets – the pseudoknot (PK) site and the three-way junction (3WJ) site (PDB code: 4lvx). The aptamer bases U7 and U42 of the PK-site interact with the ligand through at least two hydrogen bonds. Similar interactions were observed between ligand present in 3WJ-site and aptamer bases U25 and C53. However, our calculations reveal that the sum of binding energies of nucleobase:ligand pairs at the PK site (–37 kcal/mol) is greater than the 3WJ site (– 31 kcal/mol). Thus, the inability of ligand at 3WJ site to reach the interaction energy threshhold required for triggering gene expression explains the experimentally observed generalization that only the PK site ligand triggers the gene expression in THF riboswitches (45).

## DISCUSSION AND CONCLUSIONS

In the present work, we combined the techniques of structural bioinformatics and quantum chemical calculations to analyse the nature of hydrogen bonded contacts between ligands and RNA in aptamer:ligand complexes. Altogether, 80 nonredundant base:ligand pairs were identified that contain at least two hydrogen bonds. The bound ligands differ in size and complexity, where the specific functional groups act as key instruments of molecular recognition. In synchrony with a previous study (1), we found that coenzymes constitute the highest proportion of the bound ligands, which is followed by nucleobases. Further, the aptamer nucleobases most commonly interact with the ligand through the W edge, and the uracil is the most commonly interacting aptamer nucleobase.

On the basis of the chemical structure of the bound ligand, we classified the base:ligand pairs into five basic types-B:B, B:A, B:Ph, B:S and B:O. Among these, B:B interactions were further classified into two types-those involving nucleobases as ligands (e.g. 2U:ADE, PDB code: 1y26), and those involving nucleobase moiety as a part of the chemical structure of the ligands (e.g. G:TPP, PDB code: 2cky). Such contacts span seven (cWW, tWW, cWH, tWH, cWS and tWS and cHS) distinct base pairing geometries, and largely possess structural characteristics similar to base pairs found in available RNA crystal structures (19,24). Our crystal structure analysis further revealed two types of B:A contacts involving either the free amino acid as ligand (e. g. G:LYS pair (PDB code: 3d0u) or amino acid moiety as a part of the chemical structure of the ligand (e.g. MET, HEY and ORN are a part of SAM (PDB code: 4l81), SAH (PDB code: 3gx3) and SFG (PDB code: 3gx2) respectively). The B:Ph interactions, on the other hand, involve a total of three contacts, and possess interaction patterns similar to one of the base-backbone phosphate interaction types (4BPh) observed previously in RNA structures (30). However, B:S and B:O contacts involve only a few specific examples.

QM calculations reveal that more than one-third of the pairs belonging to the highly populated interaction type, the W-edge B:B pairing, possess three strong H-bonds, and significant (–14 to –32 kcal/mol) strengths, which are comparable to the strength of canonical (cWW) A:U and G:C RNA base pairs (–15 and –29 kcal/mol respectively (46)). However, the strength of S-edge B:B base pairs (–11 to –18 kcal/mol) is relatively weaker than the W-edge pairs. In contrast, the B:A interactions exhibit three distinct geometries involving (i) both –NH_2_ (donor) and –CO_2_H (donor or acceptor) of the amino acid with the aptamer nucleobase and a relatively wider (–8 to –40 kcal/mol) range of stabilization energies, (ii) –NH_2_ as both donor and acceptor, without the involvement of –CO_2_H, and a significant (–23 kcal/mol) interaction strength and (iii) Both oxygen atoms of –CO_2_H as acceptors, without the involvement of –NH_2_ and a relatively weaker (by up to –21 kcal/mol) interaction strength. On the other hand, B:Ph interactions, involve at least three hydrogen bonds. Further, significant binding strengths (–31 to –48 kcal/mol) indicates that B:Ph interactions are the strongest among all base:ligand pairs, and are even stronger than canonical base pairing interactions in RNA. It is noteworthy, that B:Ph interactions are stronger than the usual base-phosphate interactions (–1 to –10 kcal/mol) in functional RNA (30), mainly because of positive charge on the ligand, which possibly result in enhanced electrostatic interactions. Although B:S interactions have moderate (–13 kcal/mol and –20 kcal/mol) binding strength, the strength of B:O pairs (–3 kcal/mol and –21 kcal/mol), however, depends on the interacting (W or S) edge of the aptamer base and the type of hydrogen bonds.

In summary, our calculations yield important insights into the abundance and strength of base:ligand interactions. Due to close similarities with strong and structurally-important canonical and non-canonical base pairs observed in RNA macromolecular structures, our results reiterate that base:ligand interactions provide significant stability to the aptamer:ligand complexes and thereby play an important role in ligand recognition. Indeed, examples of aptamer base:ligand pairs in higher order motifs in the ligand binding pocket are crucial for transfer of chemical information from binding pocket to expression platform. Thus, the present analysis of the base:ligand interactions may help trigger advances in RNA based drug design and development.

