## Supplementary for "Pairing interactions between nucleobases and ligands in aptamer:ligand complexes of riboswitches: Crystal structure analysis, classification, optimal structures and accurate interaction energies"

**Table S1**: RMSD (Å) and Interaction energies (kcal/mol) of base:ligand pairs.[a]

| Base:Ligand interactions | Geometry | RMSD | Interaction energy |
|---|---|---|---|
| A:FMN | cWW | – | –15.1 |
| A:RBF | | 1.799 | –14.8 |
| U:29G | | 0.273 | –18.5 |
| 1U:3AY | | 0.203 | –18.2 |
| 2U:3AY | | 0.197 | –18.2 |
| U:A2F | | – | –13.2 |
| 2U:ADE | | – | –15.3 |
| 2U:N6M | | 0.260 | –15.3 |
| 2C:29H | | 0.289 | –29.2 |
| C:2BP | | 0.378 | –15.2 |
| 2C:5AZ | | 0.208 | –31.6 |
| 2C:5GP | | – | –30.0 |
| C:6GO | | 0.359 | –11.1 |
| C:6GU | | 0.347 | –16.5 |
| C:C2E | | – | –32.3 |
| C:DGP | | 1.430 | –29.9 |
| C:PQ0 | | 0.247 | –30.2 |
| 1C:PRF | | 0.279 | –28.6 |
| C:XAN | | 0.336 | –17.9 |
| A:PQ0 | tWS | 0.172 | –12.8 |
| A:PRF | | 0.290 | –12.6 |
| U:2BP | | 0.204 | –19.8 |
| U:6GO | | 0.370 | –12.3 |
| U:6GU | | 0.195 | –19.9 |
| 2U:7DG | | 0.327 | –13.7 |
| 1U:ADE | | – | –17.3 |
| 1U:FFO | | 0.468 | –19.6 |
| 2U:FFO | | – | –16.2 |
| 1U:N6M | | 0.217 | –17.6 |
| U:XAN | | 0.825 | –17.9 |
| C:29G | | 0.762 | –19.9 |
| 1C:29H | | 0.751 | –22.7 |

| | | | |
|---|---|---|---|
| 1C:5AZ | | 0.118 | –19.8 |
| 1C:5GP | | – | –19.6 |
| C:GMP | | – | –19.3 |
| 1U:7DG | tWW | 0.339 | –18.6 |
| 2U:H4B | | – | –18.4 |
| C:7DG | | 0.984 | –29.4 |
| C:FFO | | – | –17.8 |
| C:H4B | | 0.772 | –12.7 |
| 2C:PRF | | 0.714 | –13.5 |
| C:THF | | – | –12.6 |
| 1U:SAM | cWH | – | –18.5 |
| 3U:SAM | | 0.630 | –20.7 |
| U:SFG | | 1.744 | –22.9 |
| U:SAH | | 0.634 | –28.6 |
| 1U:H4B | cWS | 0.769 | –18.3 |
| U:LYA | | 0.479 | –13.7 |
| U:PRF | | 0.277 | –20.1 |
| U:THF | | 0.246 | –18.5 |
| G:C2E | tWH | – | –24.7 |
| U:6AP | | 0.196 | –16.5 |
| 2U:SAM | | 2.039 | –14.1 |
| U:PQ0 | cHS | 1.491 | –10.6 |
| G:2QB | tSW | 1.070 | –15.9 |
| G:BFT | | 1.443 | –12.6 |
| G:D2X | | – | –13.5 |
| G:EEM | | 0.715 | –12.2 |
| 2G:SAM | | 1.014 | –12.4 |
| G:TPP | | 2.175 | –10.9 |
| G:TPS | | 0.462 | –10.8 |
| A:SFG | cSW | 0.830 | –17.5 |
| A:SAH | | – | –15.5 |
| A:SAM | | 1.013 | – |
| A:SAM | B:A | 0.953 | –18.8 |
| G:SAH | | – | –18.2 |
| G:SAM | | – | –4.1 |
| G:LYS | | 0.932 | –20.8 |
| 1G:SAM | | – | – |
| 3G:SAM | | 1.526 | –39.8 |

| | | | |
|---|---|---|---|
| G:SFG (S edge) | | 1.168 | –22.5 |
| G:SFG (W edge) | | – | –7.0 |
| U:GLY | | 1.420 | –8.0 |
| G:D2X | B:Ph | – | –3.7 |
| G:G6P | | – | –31.1 |
| G:PYI | | 2.716 | –48.4 |
| G:RS3 (W edge) | B:S | 0.698 | –19.9 |
| G:RS3 (S edge) | | – | –12.8 |
| A:PRF | B:O | – | –2.6 |
| U:AMZ | | 0.646 | –21.1 |

[a]grey filled cells correspond to hydrogen-only optimized base:pairs.

**Table S2.** Hydrogen bonding properties of B:B nucleobase:ligand pairs involving cWW pairing geometries.

| Pair | H–bond interactions | D–H…A (Å) | A–H (Å) | D–H…A (°) |
|---|---|---|---|---|
| A:FMN | N6(A)–H…O(FMN) | 2.92 | 1.98 | 155.25 |
| | N(FMN)–H…N1(A) | 3.00 | 2.02 | 162.79 |
| A:RBF | N6(A)–H…O(RBF) | 2.97 | 1.96 | 173.54 |
| | N(RBF)–H…N1(A) | 2.84 | 1.79 | 179.15 |
| U:29G | N(29G)–H…O4(U) | 2.89 | 1.87 | 179.79 |
| | N3(U)–H…N1(29G) | 2.93 | 1.88 | 179.65 |
| | O2(U)–H…N(29G) | 2.95 | 1.94 | 179.53 |
| 1U:3AY | N(3AY)–H…O2(U) | 2.97 | 1.96 | 175.46 |
| | N3(U)–H…N(3AY) | 2.88 | 1.82 | 179.56 |
| | N(3AY)–H…O4(U) | 2.94 | 1.92 | 177.51 |
| 2U:3AY | N(3AY)–H…O4(U) | 2.93 | 1.91 | 179.20 |
| | N3(U)–H…N(3AY) | 2.90 | 1.84 | 179.77 |
| | N(3AY)–H…O2(U) | 2.95 | 1.94 | 176.26 |
| U:A2F | N6(A2F)–H…O4(U) | 2.86 | 1.83 | 179.33 |
| | N3(U)–H…N1(A2F) | 3.00 | 1.97 | 172.03 |
| 2U:ADE | N6(ADE)–H…O4(U) | 2.93 | 1.92 | 174.36 |
| | N3(U)–H…N1(ADE) | 2.84 | 1.79 | 179.74 |
| 2U:N6M | N6(N6M)–H…O4(U) | 2.96 | 1.94 | 177.40 |
| | N3(U)–H…N1(N6M) | 2.84 | 1.79 | 179.71 |
| 2C:29H | N2(29H)–H…O2(C) | 2.91 | 1.89 | 178.55 |
| | N1(29H)–H…N3(C) | 2.90 | 1.86 | 177.42 |
| | N4(C)–H…O6(29H) | 2.84 | 1.81 | 178.86 |
| C:2BP | N2(2BP)–H…N3(C) | 2.96 | 1.94 | 177.08 |
| | N4(C)–H…N1(2BP) | 2.98 | 1.94 | 179.36 |
| 2C:5AZ | N2(5AZ)–H…O2(C) | 2.93 | 1.91 | 175.82 |
| | N1(5AZ)–H…N3(C) | 2.86 | 1.82 | 178.81 |
| | N4(C)–H…O6(5AZ) | 2.80 | 1.77 | 179.44 |
| 2C:5GP | N2(5GP)–H…O2(C) | 2.89 | 1.89 | 176.95 |
| | N1(5GP)–H…N3(C) | 2.97 | 1.90 | 175.87 |
| | N4(C)–H…O6(5GP) | 2.82 | 1.80 | 175.92 |
| C:6GO | N2(6GO)–H…N3(C) | 3.01 | 2.02 | 163.25 |
| | N4(C)–H…N1(6GO) | 3.09 | 2.06 | 179.26 |
| C:6GU | N2(6GU)–H…N3(C) | 2.94 | 1.92 | 177.85 |
| | N4(C)–H…N1(6GU) | 3.03 | 2.01 | 176.95 |
| C:C2E | N2(C2E)–H…O2(C) | 2.84 | 1.81 | 179.91 |
| | N1(C2E)–H…N3(C) | 2.88 | 1.85 | 176.57 |
| | N4(C)–H…O6(C2E) | 2.84 | 1.81 | 174.55 |
| C:DGP | N2(DGP)–H…O2(C) | 2.91 | 1.89 | 179.10 |
| | N1(DGP)–H…N3(C) | 2.92 | 1.88 | 177.16 |
| | N4(C)–H…O6(DGP) | 2.80 | 1.77 | 179.38 |

| | | | | |
|---|---|---|---|---|
| C:PQ0 | N2(PQ0)–H…O2(C) | 2.91 | 1.89 | 179.87 |
| | N1(PQ0)–H…N3(C) | 2.91 | 1.88 | 177.99 |
| | N4(C)–H…O6(PQ0) | 2.80 | 1.77 | 179.50 |
| 1C:PRF | N2(PRF)–H…O2(C) | 2.93 | 1.91 | 178.30 |
| | N1(PRF)–H…N3(C) | 2.93 | 1.90 | 177.62 |
| | N4(C)–H…O6(PRF) | 2.80 | 1.76 | 179.26 |
| C:XAN | N4(C)–H…O6(XAN) | 2.85 | 1.82 | 176.40 |
| | N1(XAN)–H…O2(C) | 2.96 | 1.93 | 171.49 |

**Table S3.** Hydrogen bonding properties of B:B nucleobase:ligand pairs involving tWS pairing geometries.

| **RNA:Ligand interactions** | **H–bond interactions** | **D–H…A (Å)** | **A–H (Å)** | **D–H…A (°)** |
|---|---|---|---|---|
| A:PQ0 | N6(A)–H…N3(PQ0) | 3.03 | 2.01 | 177.22 |
| | N2(PQ0)–H…N1(A) | 2.95 | 1.92 | 176.65 |
| A:PRF | N6(A)–H…N3(PQ0) | 3.01 | 1.98 | 177.03 |
| | N2(PQ0)–H…N1(A) | 2.98 | 1.95 | 178.14 |
| U:2BP | N2(2BP)–H…O2(U) | 3.07 | 2.06 | 170.50 |
| | N3(U)–H…N3(2BP) | 2.82 | 1.77 | 179.77 |
| | N9(2BP)–H…O4(U) | 2.92 | 1.94 | 159.17 |
| U:6GO | N2(6GO)–H…O2(U) | 2.93 | 1.92 | 174.02 |
| | N3(U)–H…N3(6GO) | 2.96 | 1.92 | 171.45 |
| U:6GU | N2(6GU)–H…O2(U) | 3.08 | 2.08 | 170.47 |
| | N3(U)–H…N3(6GU) | 2.82 | 1.77 | 179.34 |
| | N9(6GU)–H…O4(U) | 2.90 | 1.92 | 160.33 |
| 2U:7DG | N2(7DG)–H…O2(U) | 2.86 | 1.84 | 176.71 |
| | N3(U)–H…N3(7DG) | 2.99 | 1.96 | 168.85 |
| 1U:ADE | N3(U)–H…N3(ADE) | 2.86 | 1.82 | 176.58 |
| | N9(ADE)–H…O4(U) | 2.84 | 1.84 | 161.88 |
| 1U:FFO | N2(FFO)–H…O2(U) | 2.91 | 1.89 | 179.25 |
| | N3(U)–H…N3(FFO) | 2.92 | 1.87 | 178.92 |
| | N9(FFO)–H…O4(U) | 2.94 | 1.92 | 175.95 |
| 2U:FFO | N2(FFO)–H…O4(U) | 2.88 | 1.86 | 179.36 |
| | N3(U)–H…N3(FFO) | 2.91 | 1.87 | 179.76 |
| | N9(FFO)–H…O2(U) | 2.96 | 1.94 | 176.71 |
| 1U:N6M | N3(U)–H…N3(N6M) | 2.85 | 1.81 | 176.65 |
| | N9(N6M)–H…O4(U) | 2.83 | 1.84 | 163.05 |
| U:XAN | N3(U)–H…O(XAN) | 2.86 | 1.83 | 174.44 |
| | N(XAN)–H…O4(U) | 2.76 | 1.73 | 178.59 |
| C:29G | N2(29G)–H…N3(C) | 2.93 | 1.90 | 174.60 |
| | N4(C)–H…N3(29G) | 2.94 | 1.91 | 175.24 |
| 1C:29H | N2(29H)–H…N3(C) | 2.85 | 1.81 | 176.50 |
| | N4(C)–H…N3(29H) | 2.96 | 1.93 | 172.68 |

| | | | | |
|---|---|---|---|---|
| 1C:5AZ | N(5AZ)–H…N3(C) | 2.85 | 1.81 | 178.33 |
| | N4(C)–H…N(5AZ) | 2.99 | 1.97 | 175.81 |
| 1C:5GP | N2(5GP)–H…N3(C) | 2.92 | 1.88 | 169.41 |
| | N4(C)–H…N3(5GP) | 3.03 | 2.00 | 177.20 |
| C:GMP | N2(GMP)–H…N3(C) | 2.91 | 1.88 | 172.09 |
| | N4(C)–H…N3(GMP) | 2.98 | 1.96 | 179.20 |

**Table S4.** Hydrogen bonding properties of B:B nucleobase:ligand pairs involving tWW pairing geometries.

| RNA:Ligand interactions | H–bond interactions | D–H…A (Å) | A–H (Å) | D–H…A (°) |
|---|---|---|---|---|
| 1U:7DG | N3(U)–H…O6(7DG) | 2.82 | 1.77 | 176.92 |
| | N1(7DG)–H…O4(U) | 2.79 | 1.76 | 176.37 |
| 2U:H4B | N3(U)–H…O6(H4B) | 2.75 | 1.70 | 178.62 |
| | N1(H4B)–H…O4(U) | 2.81 | 1.79 | 174.34 |
| C:7DG | N2(7DG)–H…O2(C) | 2.87 | 1.98 | 144.83 |
| | N1(7DG)–H…O2(C) | 2.82 | 1.88 | 150.50 |
| | O2'(C)–H…O6(7DG) | 2.65 | 1.66 | 173.89 |
| C:FFO | N2(FFO)–H…N3(C) | 3.12 | 2.10 | 179.85 |
| | N1(FFO)–H…O2(C) | 2.89 | 1.88 | 169.55 |
| C:H4B | N2(H4B)–H…N3(C) | 3.19 | 2.17 | 172.52 |
| | N1(H4B)–H…O2(C) | 2.96 | 1.96 | 167.05 |
| 2C:PRF | N2(PRF)–H…N3(C) | 3.17 | 2.16 | 175.83 |
| | N1(PRF)–H…O2(C) | 2.95 | 1.94 | 168.62 |
| C:THF | N2(THF)–H…N3(C) | 3.19 | 2.18 | 175.37 |
| | N1(THF)–H…O2(C) | 2.96 | 1.96 | 167.82 |

**Table S5.** Hydrogen bonding properties of B:B nucleobase:ligand pairs involving cWH pairing geometries.

| RNA:Ligand interactions | H–bond interactions | D–H…A (Å) | A–H (Å) | D–H…A (°) |
|---|---|---|---|---|
| 1U:SAM | N6(SAH)–H…O4(U) | 2.89 | 1.87 | 168.05 |
| | N3(U)–H…N7(SAH) | 2.86 | 1.83 | 175.14 |
| 3U:SAM | N6(SAM)–H…O4(U) | 2.91 | 1.90 | 168.32 |
| | N3(U)–H…N7(SAM) | 2.85 | 1.82 | 174.88 |
| U:SFG | N6(SFG)–H…O4(U) | 2.99 | 2.01 | 162.32 |
| | N3(U)–H…N7(SFG) | 2.83 | 1.81 | 168.69 |
| U:SAH | N6(SAH)–H…O4(U) | 2.97 | 1.96 | 168.74 |
| | N3(U)–H…N7(SAH) | 2.83 | 1.79 | 174.45 |

| **Table S6.** Hydrogen bonding properties of B:B nucleobase:ligand pairs involving cWS pairing geometries. | | | | |
|---|---|---|---|---|
| **RNA:Ligand interactions** | **H–bond interactions** | **D–H…A (Å)** | **A–H (Å)** | **D–H…A (°)** |
| 1U:H4B | N2(FFO)–H…O4(U) | 2.91 | 1.89 | 179.76 |
| | N3(U)–H…N3(FFO) | 2.91 | 1.86 | 178.93 |
| | N9(FFO)–H…O2(U) | 3.00 | 1.99 | 176.85 |
| U:LYA | N2(FFO)–H…O4(U) | 3.11 | 2.10 | 170.78 |
| | N3(U)–H…N3(FFO) | 3.00 | 1.99 | 165.75 |
| | N9(FFO)–H…O2(U) | 3.03 | 2.07 | 155.38 |
| U:PRF | N2(FFO)–H…O4(U) | 2.96 | 1.94 | 175.96 |
| | N3(U)–H…N3(FFO) | 2.82 | 1.77 | 179.65 |
| | N9(FFO)–H…O2(U) | 2.97 | 1.98 | 164.35 |
| U:THF | N2(THF)–H…O4(U) | 2.92 | 1.89 | 178.76 |
| | N3(U)–H…N3(THF) | 2.91 | 1.86 | 178.66 |
| | N9(THF)–H…O2(U) | 2.99 | 1.97 | 178.66 |

| **Table S7.** Hydrogen bonding properties of B:B nucleobase:ligand pairs involving tWH pairing geometries. | | | | |
|---|---|---|---|---|
| **RNA:Ligand interactions** | **H–bond interactions** | **D–H…A (Å)** | **A–H (Å)** | **D–H…A (°)** |
| G:C2E | N1(G)–H…N7(C2E) | 2.75 | 1.72 | 174.67 |
| | N2(G)–H…O6(C2E) | 2.83 | 1.85 | 160.86 |
| U:6AP | N6(6AP)–H…O4(U) | 2.97 | 1.96 | 169.48 |
| | N3(U)–H…N7(6AP) | 2.81 | 1.76 | 174.72 |
| 2U:SAM | N6(SAM)–H…O2(U) | 3.04 | 2.04 | 167.65 |
| | N3(U)–H…N7(SAM) | 2.82 | 1.78 | 172.48 |

| **Table S8.** Hydrogen bonding properties of B:B nucleobase:ligand pairs involving tSW pairing geometries. | | | | |
|---|---|---|---|---|
| **RNA:Ligand interactions** | **H–bond interactions** | **D–H…A (Å)** | **A–H (Å)** | **D–H…A (°)** |
| G:2QB | N2(G)–H…N(2QB) | 3.06 | 2.04 | 174.19 |
| | N(2QB)–H…N3(G) | 3.02 | 2.00 | 168.72 |
| | O2'(G)–H…N(2QB) | 3.09 | 2.15 | 161.48 |
| G:BFT | N2(G)–H…N(BFT) | 3.01 | 1.99 | 170.35 |
| | N(BFT)–H…N3(G) | 3.04 | 2.02 | 177.13 |
| G:D2X | N2(G)–H…N(D2X) | 3.30 | 2.31 | 166.64 |
| | O(D2X)–H…N3(G) | 3.31 | 2.33 | 170.30 |
| G:EEM | N2(G)–H…N1(EEM) | 3.11 | 2.11 | 168.66 |
| | N6(EEM)–H…N3(G) | 2.97 | 1.95 | 175.31 |
| | N6(EEM)–H…O2'(G) | 3.01 | 2.10 | 147.33 |

| | | | | |
|---|---|---|---|---|
| 2G:SAM | N2(G)–H…N1(SAM) | 3.10 | 2.08 | 174.83 |
| | N6(SAM)–H…N3(G) | 2.98 | 1.95 | 175.83 |
| | N6(SAM)–H… O2'(G) | 3.00 | 2.10 | 147.45 |
| G:TPP | N2(G)–H…N(TPP) | 2.93 | 1.90 | 174.01 |
| | N(TPP)–H…N3(G) | 3.09 | 2.08 | 166.60 |
| G:TPS | N2(G)–H…N(TPS) | 3.09 | 2.09 | 166.28 |
| | N(TPS)–H…N3(G) | 2.93 | 1.90 | 172.76 |

**Table S9.** Hydrogen bonding properties of B:B nucleobase:ligand pairs involving cSW pairing geometries.

| RNA:Ligand interactions | H–bond interactions | D–H…A (Å) | A–H (Å) | D–H…A (°) |
|---|---|---|---|---|
| A:SAH | N6(SAH)–H…N3(A) | 3.31 | 2.34 | 156.60 |
| | O2'(A)–H…N1(SAH) | 3.02 | 2.06 | 165.21 |
| A:SAM | N6(SAH)–H…N3(A) | 2.88 | 1.88 | 159.57 |
| | O2'(A)–H…N1(SAM) | 2.83 | 1.87 | 165.70 |
| A:SFG | N6(SFG)–H…N3(A) | 2.99 | 2.01 | 159.35 |
| | O2'(A)–H…N1(SFG) | 2.75 | 1.77 | 170.49 |

**Table S10.** Hydrogen bonding properties of B:B nucleobase:ligand pairs involving cHS

| RNA:Ligand interactions | H–bond interactions | D–H…A (Å) | A–H (Å) | D–H…A (°) |
|---|---|---|---|---|
| U:PQ0 | N9(PQ0)–H…O4(U) | 2.86 | 1.83 | 175.18 |
| | C5(U)–H…N3(PQ0) | 3.53 | 2.48 | 163.70 |

**Table S11.** Hydrogen bonding properties of B:A, B:S, B:Ph and B:O base:ligand interactions.

| RNA:Ligand interactions | Base pair | Edge | H–bond interactions | D–H…A (Å) | A–H (Å) | D–H…A (°) |
|---|---|---|---|---|---|---|
| A:SAM | B:A | W | N6(A)–H…O(SAM) | 2.98 | 1.97 | 174.02 |
| | | | N(SAM)–H…N1(A) | 3.13 | 2.13 | 164.73 |
| G:SAH | B:A | W | N1(G)–H…O(SAH) | 2.88 | 1.89 | 169.43 |
| | | | N2(G)–H…O(SAH) | 2.77 | 1.79 | 164.84 |
| G:LYS | B:A | S | N2(G)–H…O(LYS) | 2.99 | 1.98 | 170.01 |
| | | | N(LYS)–H…N3(G) | 3.14 | 2.29 | 141.50 |
| | | | N(LYS)–H…O2'(G) | 3.14 | 2.38 | 130.77 |
| | | | O2'(G)–H…N(LYS) | 2.94 | 1.95 | 175.19 |
| 1G:SAM | B:A | S | N2(G)–H…O(SAM) | 2.83 | 1.81 | 174.04 |
| | | | N(SAM)–H…N3(G) | 2.86 | 1.83 | 160.90 |
| | | | N(SAM)–H… O2'(G) | 2.89 | 1.88 | 165.15 |
| | | | N6(SAM)–H… O4'(G) | 3.03 | 2.10 | 150.93 |
| G:SFG | B:A | S | N2(G)–H…N(SFG) | 2.96 | 1.98 | 157.05 |

| | | | | | | |
|---|---|---|---|---|---|---|
| | | | N(SFG)–H…N3(G) | 3.08 | 2.29 | 133.84 |
| | | | C(SFG)–H…O2'(G) | 3.57 | 2.71 | 134.86 |
| U:GLY | B:A | W | N(GLY)–H…O2(U) | 3.17 | 2.17 | 168.80 |
| | | | N3(U)–H…O(GLY) | 2.90 | 1.88 | 171.09 |
| G:SAM | BA | W | N1(G)–H…O(SAM) | 3.01 | 1.99 | 179.72 |
| | | | N2(G)–H…O(SAM) | 3.57 | 2.69 | 145.57 |
| 3G:SAM | BA | S | N2(G)–H…O(SAM) | 3.19 | 2.18 | 169.90 |
| | | | O2'(G)–H…N7(SAM) | 2.83 | 1.88 | 161.96 |
| | | | O(SAM)–H… O3'(G) | 2.69 | 1.77 | 152.19 |
| G:SFG | BA | W | N1(G)–H…O(SFG) | 3.46 | 2.51 | 156.08 |
| | | | N2(G)–H…O(SFG) | 3.16 | 2.24 | 151.63 |
| G:D2X | BPh | W | OP(D2X)–H…O6(G) | 2.47 | 1.42 | 176.90 |
| | | | N1(G)–H…OP(D2X) | 2.97 | 1.99 | 162.84 |
| G:G6P | B:Ph | W | N2(G)–H…OP(G6P) | 3.07 | 2.29 | 138.83 |
| | | | N1(G)–H…OP(G6P) | 2.71 | 1.91 | 135.41 |
| | | | OP(G6P)–H…O6(G) | 2.7 | 1.8 | 148.00 |
| G:PYI | B:Ph | W | OP(PYI)–H…O6(G) | 2.62 | 1.63 | 166.66 |
| | | | OP(PYI)–H…O6(G) | 2.61 | 1.65 | 158.42 |
| | | | N1(G)–H…OP(PYI) | 2.85 | 1.84 | 167.70 |
| G:RS3 | B:S | W | O2'(RS3)–H…O6(G) | 2.88 | 1.87 | 164.95 |
| | | | N1(G)–H…O5'(RS3) | 2.68 | 1.71 | 168.37 |
| G:RS3 | BS | S | N2(G)–H…O5'(RS3) | 3.01 | 2.03 | 161.02 |
| | | | O2'(G)–H…O4'(RS3) | 3.13 | 2.32 | 140.74 |
| A:PRF | B:O | S | C(PRF)–H…N3(A) | 3.42 | 2.53 | 139.37 |
| | | | C(PRF)–H…O2'(A) | 3.37 | 2.32 | 160.57 |
| | | | N(PRF)–H…O3'(A) | 3.47 | 2.96 | 111.75 |
| U:AMZ | B:O | W | N(AMZ)–H…O2(U) | 3.09 | 2.11 | 161.35 |
| | | | N3(U)–H…N(AMZ) | 2.82 | 1.77 | 174.60 |
| | | | O5'(AMZ)–H…O4(U) | 2.87 | 1.94 | 159.29 |

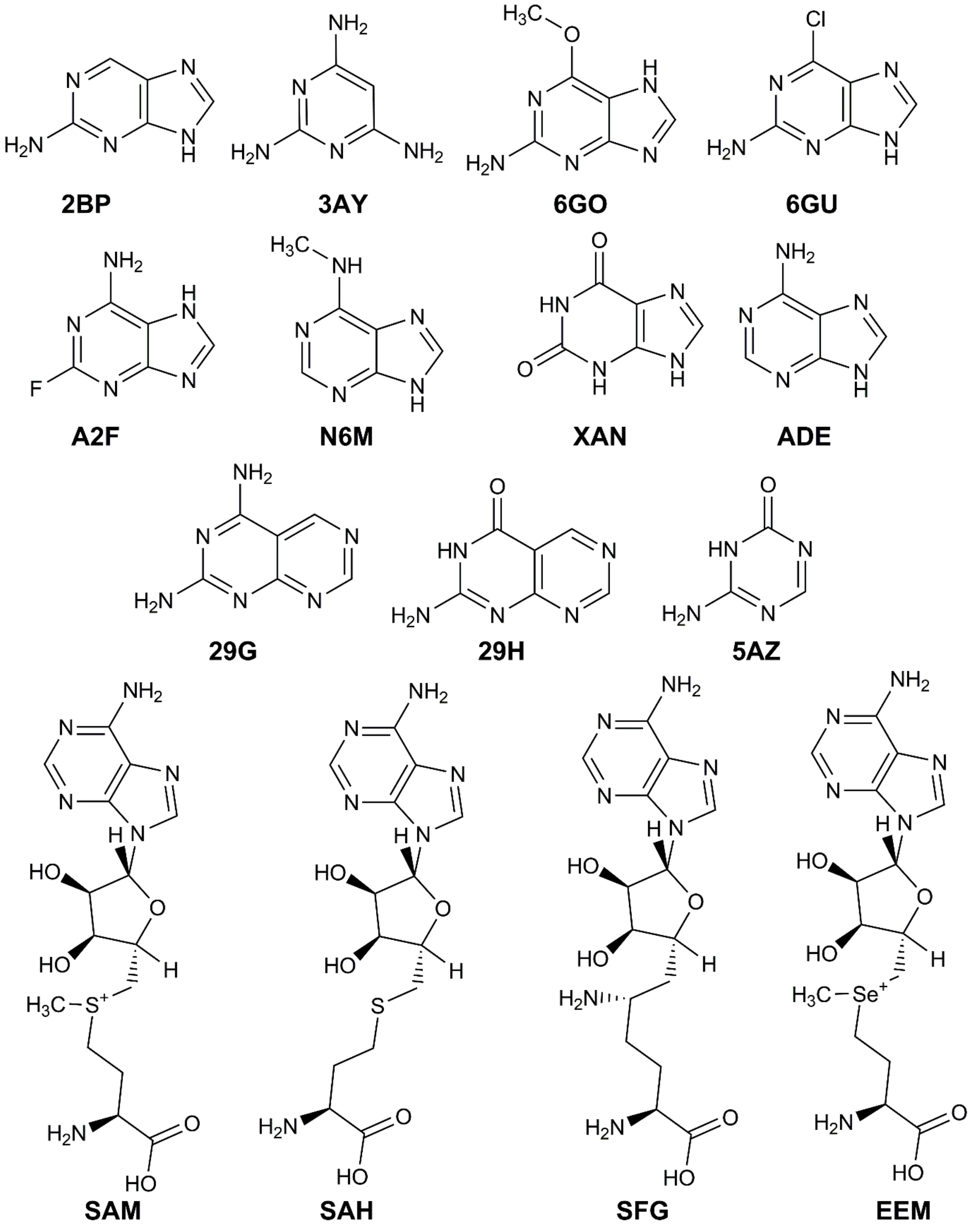


**Figure S1.** Chemical structures of the ligands (2BP, 3AY, 6GO, 6GU, A2F, N6M, XAN, ADE, 29G, 29H, 5AZ, SAM, SAH, SFG and EEM).

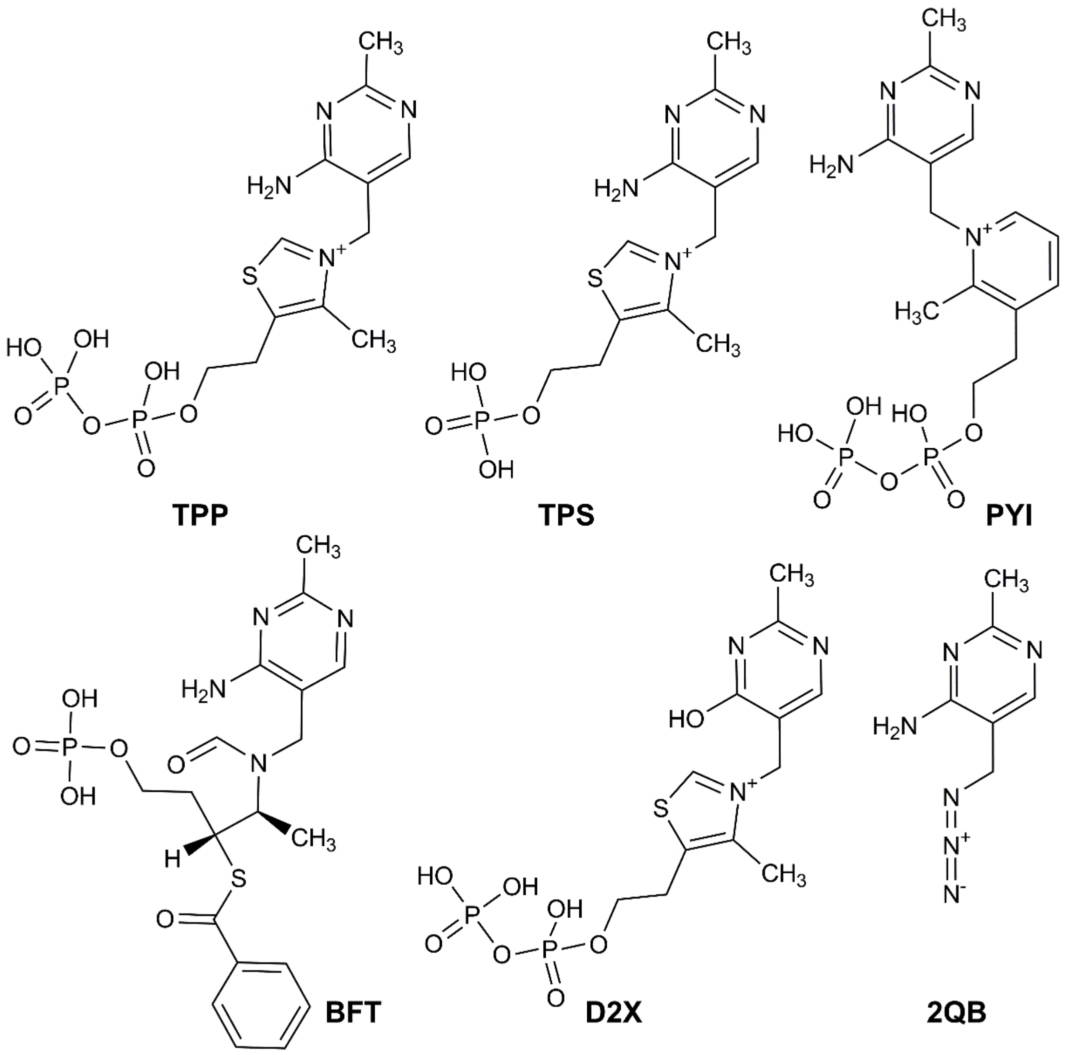


**Figure S2.** Chemical structures of the ligands (TPP, TPS, PYI, BFT, D2X and 2QB).

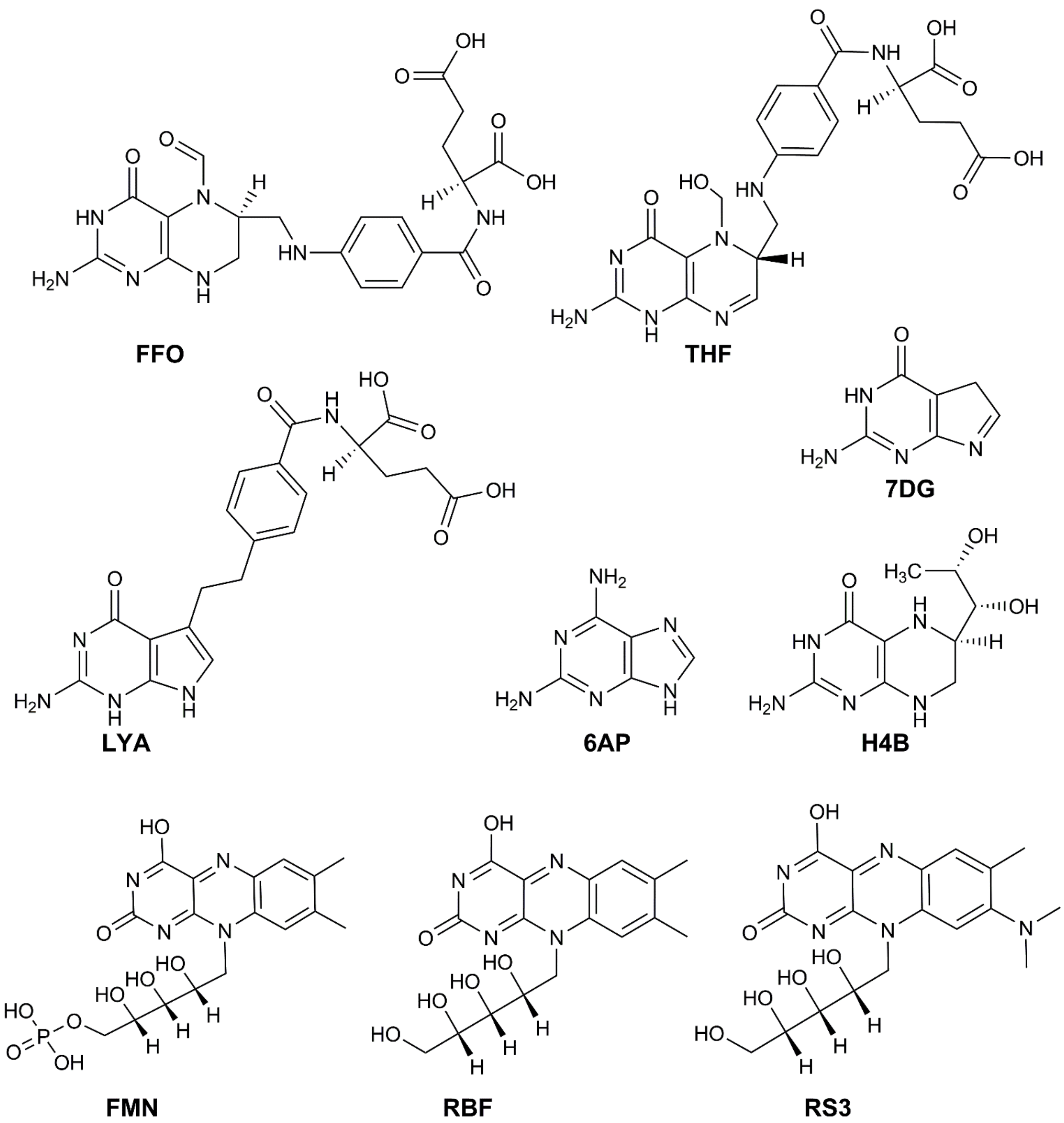


**Figure S3.** Chemical structures of the ligands (FFO, THF, LYA, 6AP, H4B, FMN, RBF and RS3).

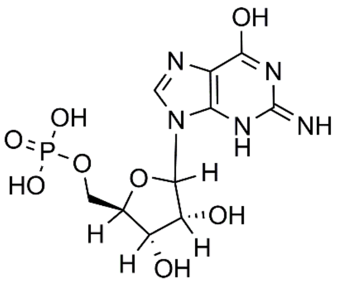

**5GP**

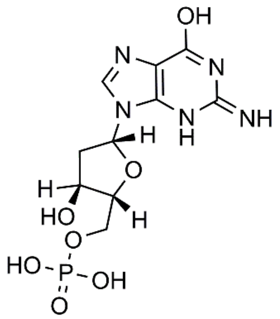

**DGP**

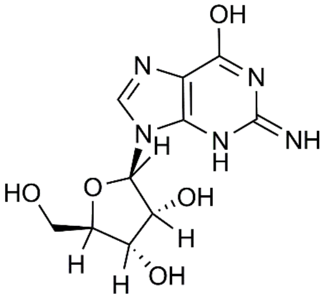

**GMP**

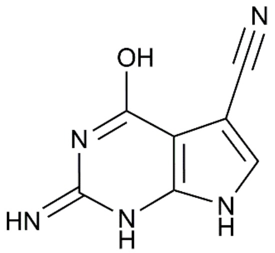

**PQ0**

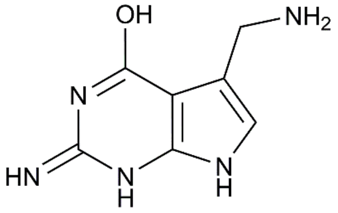

**PRF**

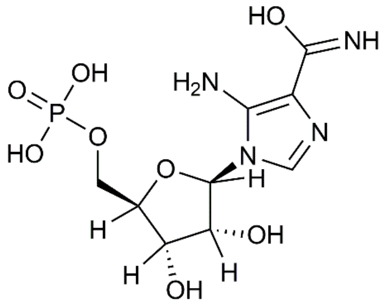

**AMZ**

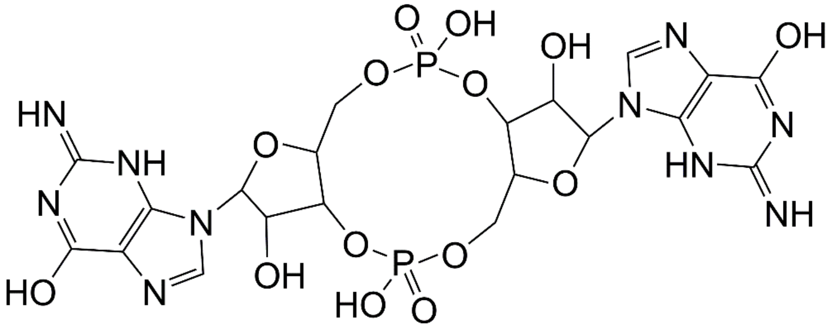

**C2E**

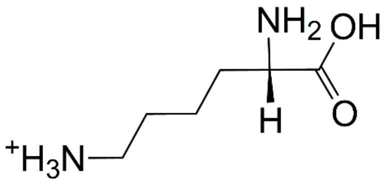

**LYS**

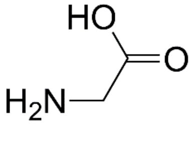

**GLY**

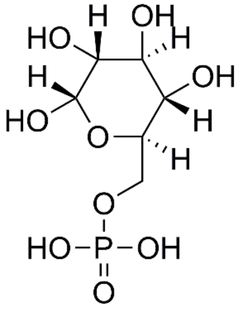

**G6P**

**Figure S4.** Chemical structures of the ligands (5GP, DGP, GMP, PQ0, PRF, AMZ, C2E, LYS, GLY and G6P).

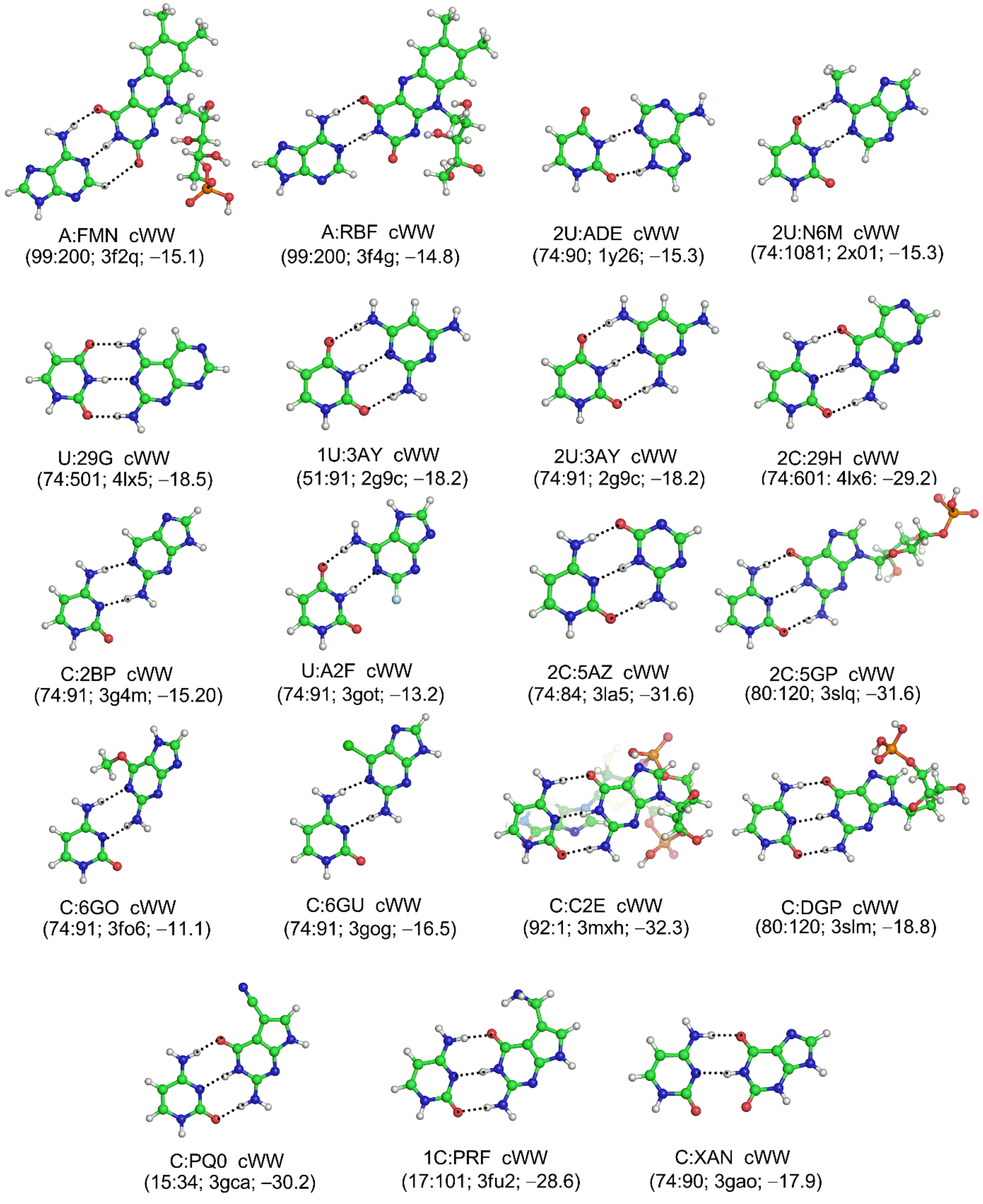


**Figure S5.** Optimized base:ligand structures that resemble cWW base pair geometry (interaction energy in kcal/mol is given in parentheses). For structures 2C:5GP and C:C2E, only the positions of hydrogen atoms were optimized, whereas full optimization was carried out for all other structures.

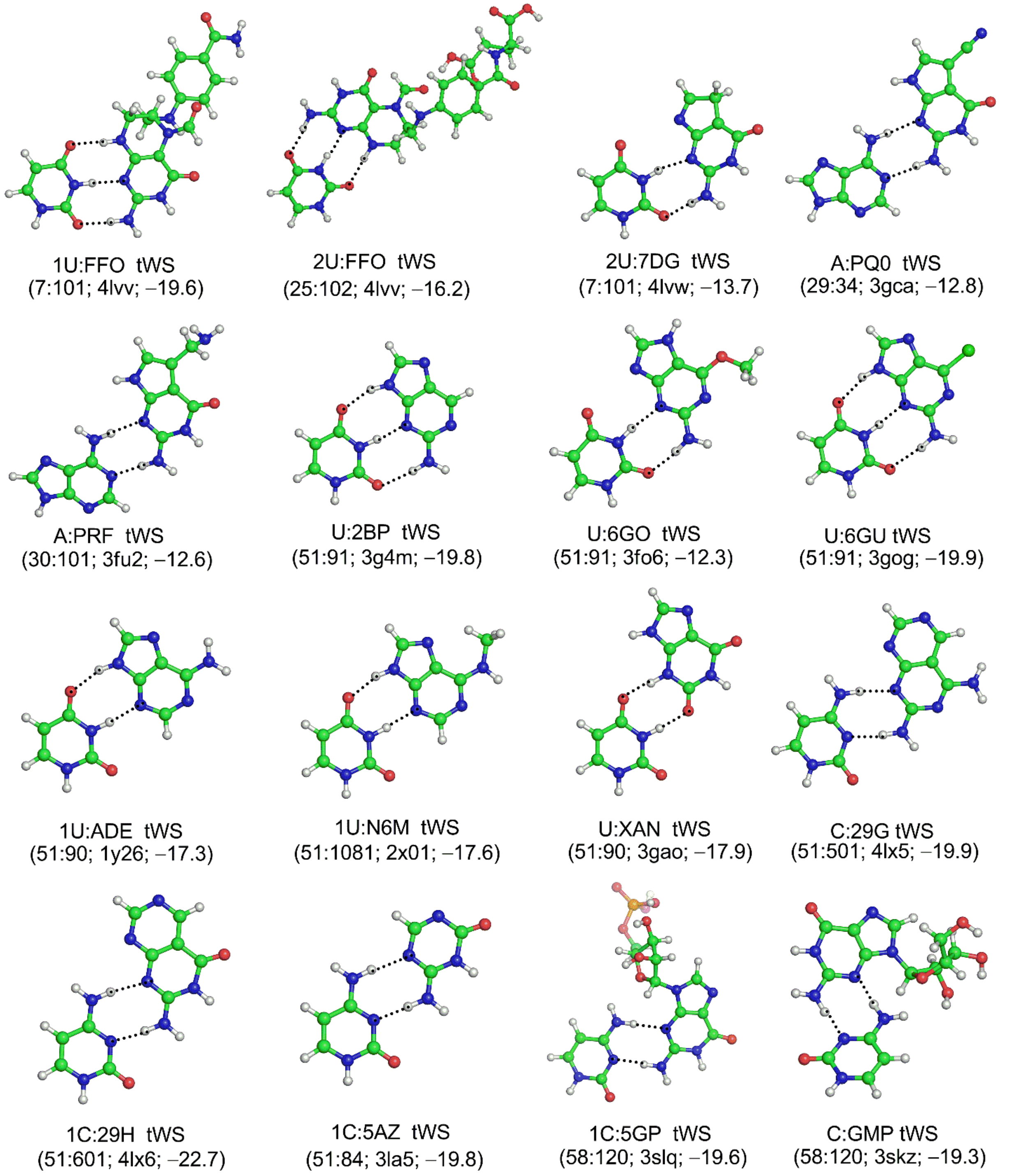


**Figure S6.** Optimized base:ligand structures that resemble tWS base pair geometry (interaction energy in kcal/mol is given in parentheses). For structures 2U:FFO and 1C:5GP, only the positions of hydrogen atoms were optimized, whereas full optimization was carried out for all other structures.

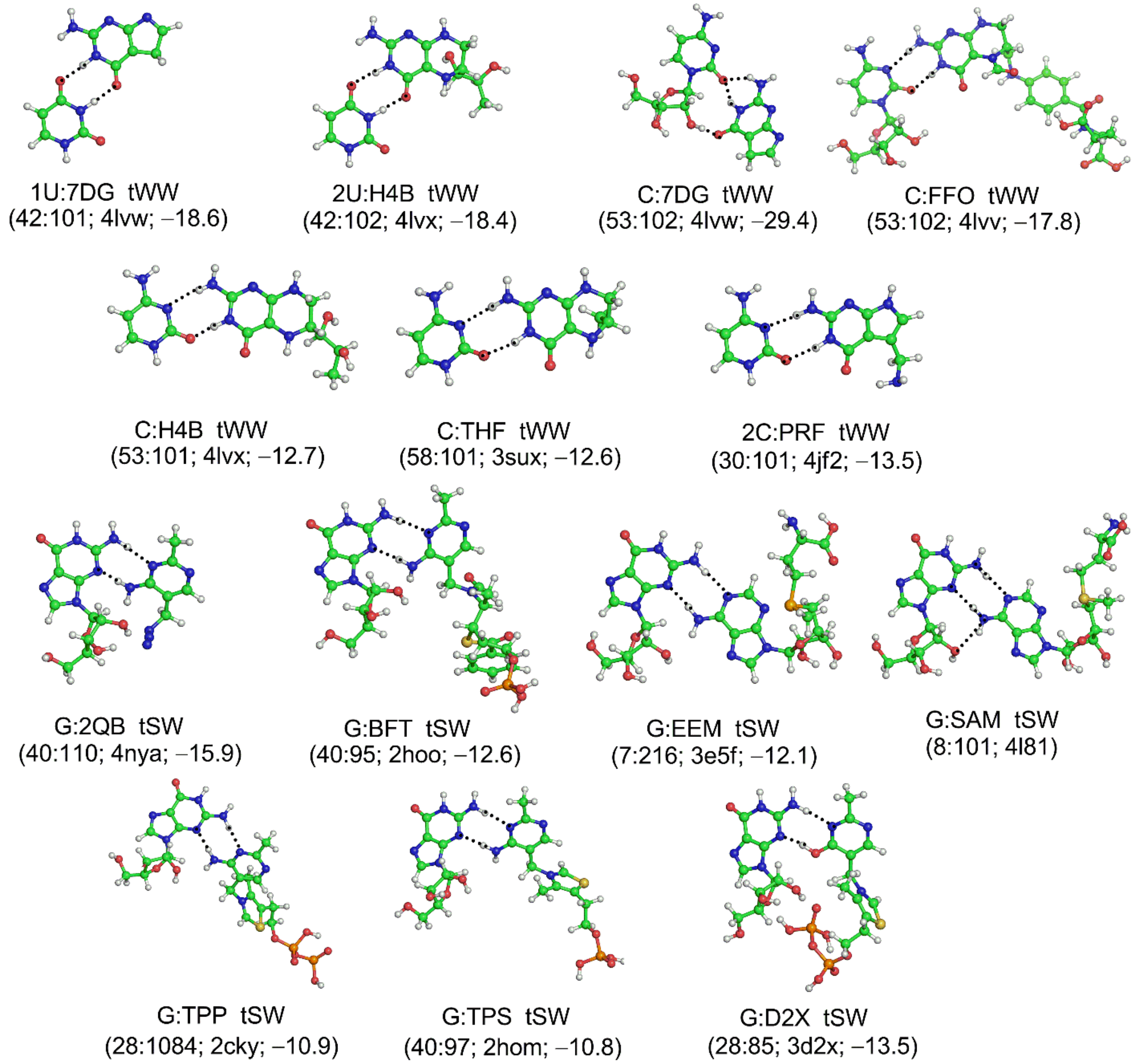


**Figure S7.** Optimized base:ligand structures that resemble either tWW or tSW base pair geometry (interaction energy in kcal/mol is given in parentheses). For structures C:FFO (tWW) and G:D2X (tSW), only the positions of hydrogen atoms were optimized, whereas full optimization was carried out for all other structures.

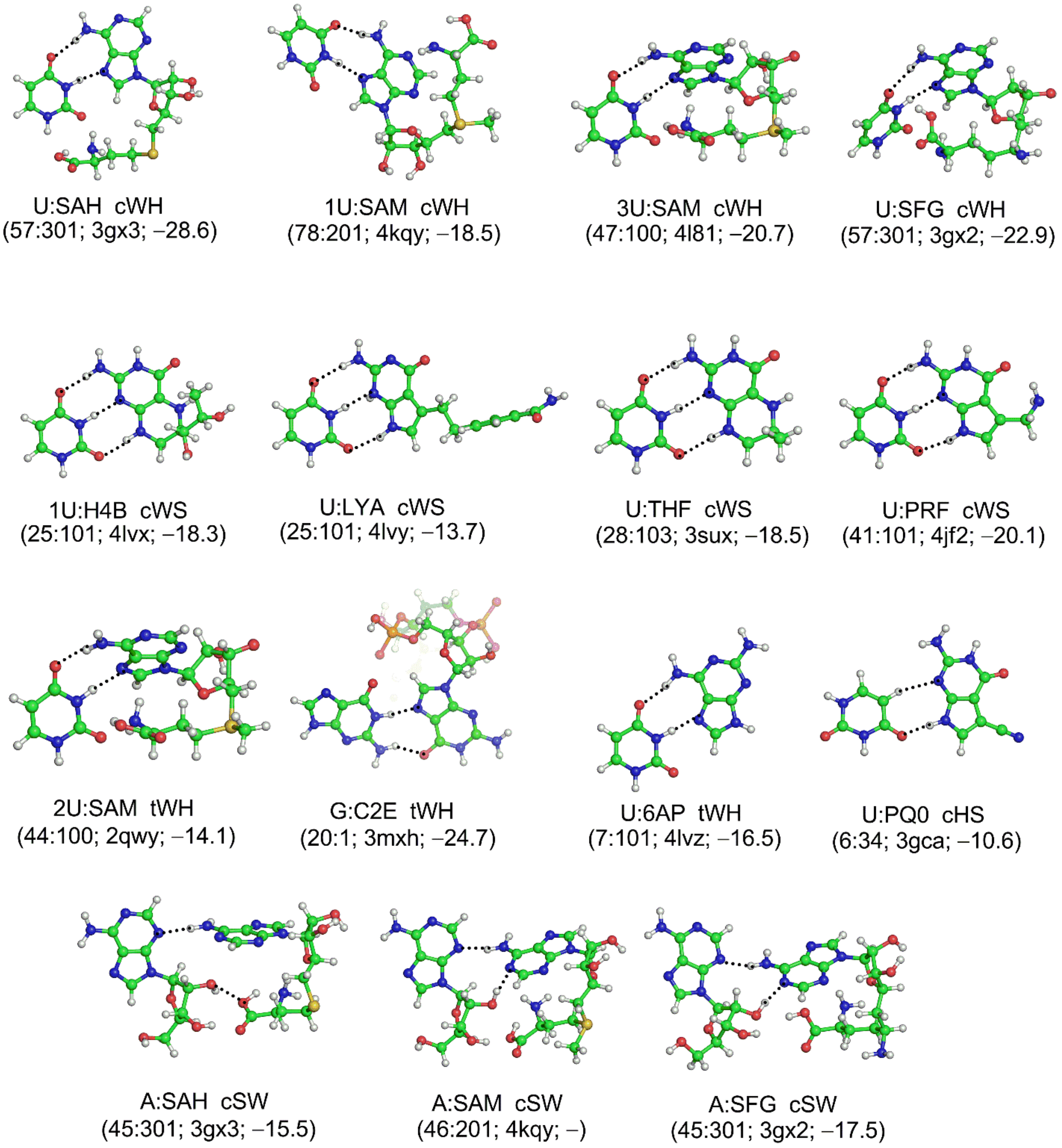


**Figure S8.** Optimized base:ligand structures that resemble either cWH, cWS, tWH, cHS or cSW base pair geometry (interaction energy in kcal/mol is given in parentheses). For structures G:C2E (tWH) and A:SAH (cSW), only the positions of hydrogen atoms were optimized, whereas full optimization was carried out for all other structures.

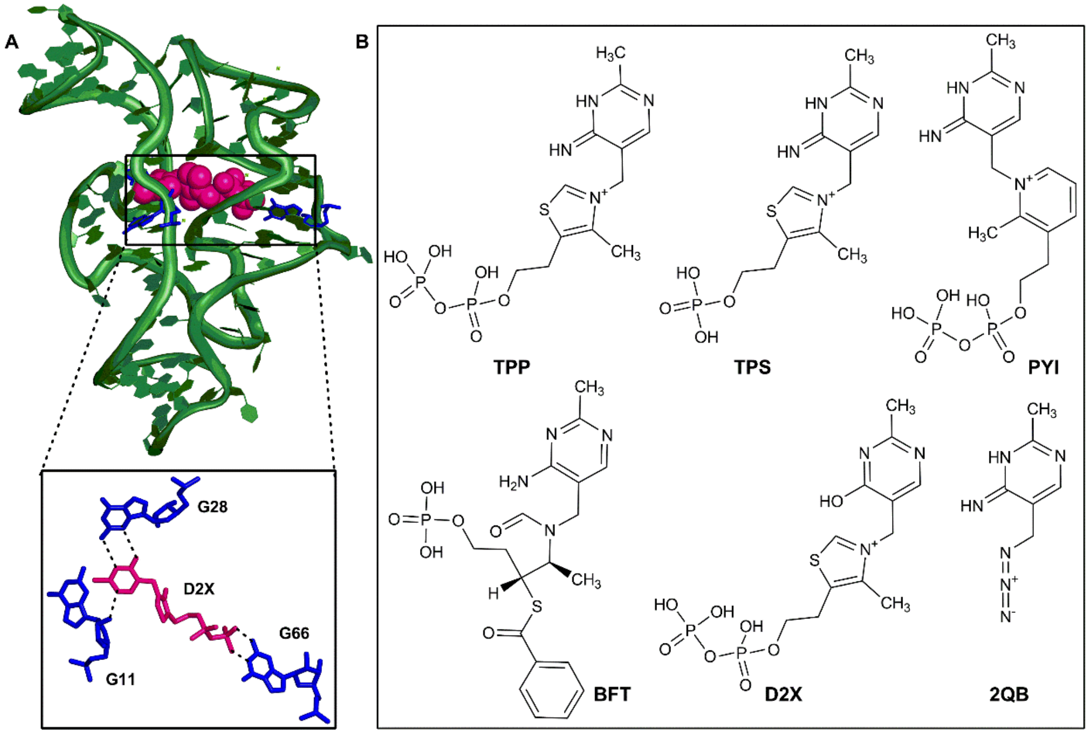


**Figure S9.** (A) D2X ligand bound to the aptamer domain of thiamine pyrophosphate riboswitch. (B) Structures of different TPP–related ligands that bind to the aptamer domain of TPP riboswitch.

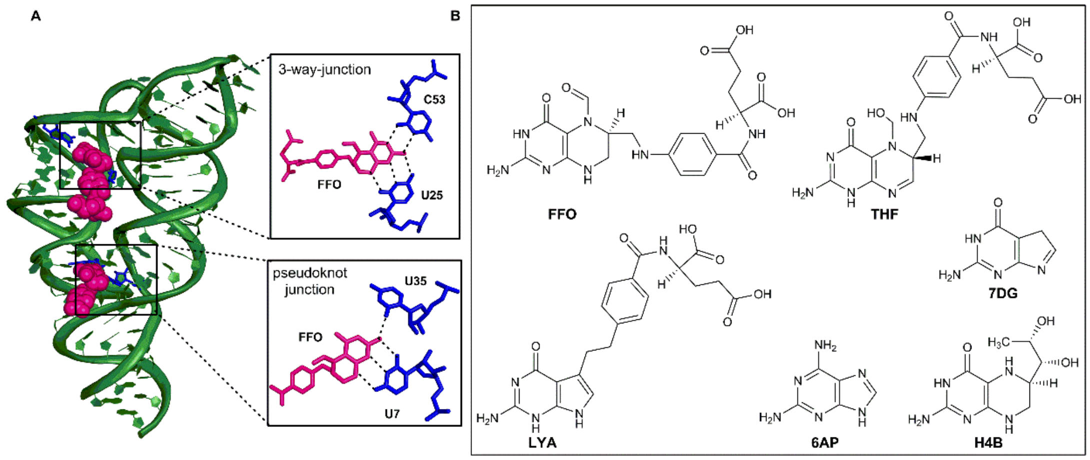


**Figure S10.** (A) FFO ligand bound to the aptamer domain of thiamine pyrophosphate riboswitch. (B) Representation of different THF–related ligands that bind to the riboswitch aptamer.

SAM riboswitch

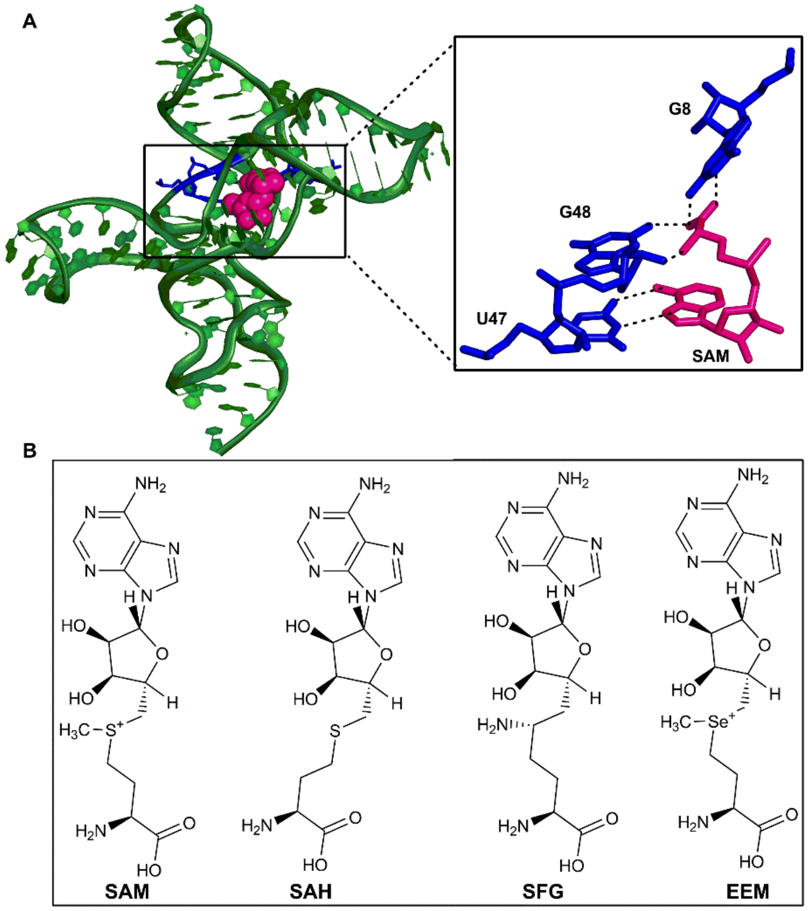


**Figure S11.** (A) SAM ligand bound in the aptamer domain of SAM riboswitch. (B) Structures of different SAM–related ligands that bind to the riboswitch aptamer.

### Purine riboswitch

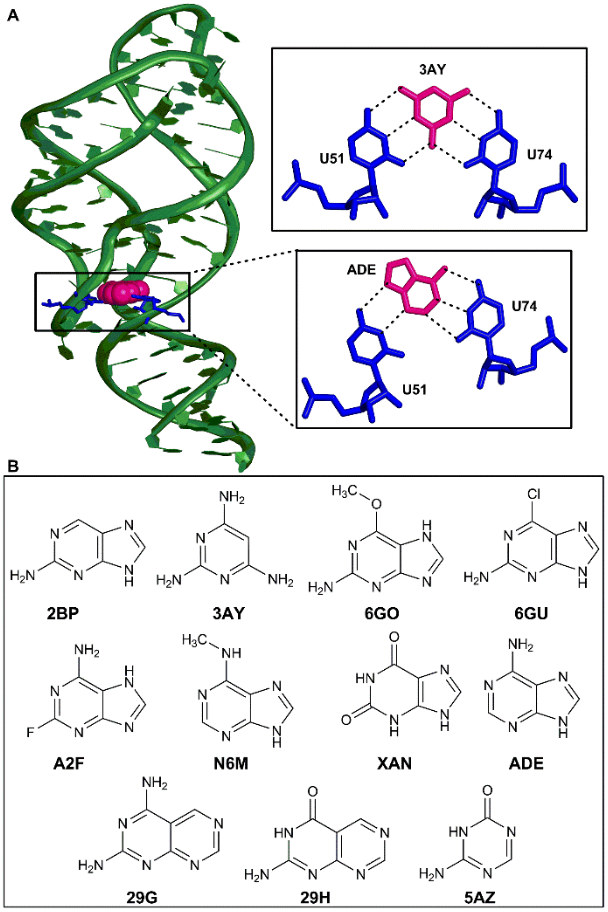


**Figure S12.** (A) ADE ligand bound in the aptamer domain of ADE riboswitch. (B) Representation of different purine ligands that bind to the riboswitch aptamer.

**Table S12.** Optimized coordinates of A:PQ0 (W-edge) complex.
Energy: –681550.0068039 (au)

| | | | |
|---|---|---|---|
| N | –6.24584 | –0.50090 | –0.08807 |
| C | –5.87647 | –1.81396 | –0.30257 |
| N | –4.57766 | –1.99109 | –0.32682 |
| C | –4.06475 | –0.72318 | –0.11754 |
| C | –2.73780 | –0.24220 | –0.03059 |
| N | –1.66895 | –1.04600 | –0.15897 |
| N | –2.56064 | 1.08242 | 0.18597 |
| C | –3.64541 | 1.87153 | 0.31552 |
| N | –4.93236 | 1.53767 | 0.25567 |
| C | –5.08148 | 0.22242 | 0.03477 |
| N | 2.46715 | 2.02137 | –0.15839 |
| C | 1.24326 | 1.42690 | 0.03978 |
| N | 1.11533 | 0.11845 | 0.20933 |
| C | 2.29546 | –0.55459 | 0.19520 |
| C | 3.58129 | –0.04779 | –0.00118 |
| C | 3.73802 | 1.36667 | –0.20499 |
| O | 4.74771 | 2.02477 | –0.38980 |
| C | 4.49554 | –1.15769 | 0.06929 |
| C | 5.90892 | –1.14349 | –0.07126 |
| N | 7.06715 | –1.16651 | –0.18195 |
| C | 3.73407 | –2.28395 | 0.30531 |
| N | 2.40796 | –1.90935 | 0.37646 |
| N | 0.15756 | 2.22750 | 0.08667 |
| H | –6.61494 | –2.59332 | –0.43333 |
| H | –1.83233 | –2.02721 | –0.31790 |
| H | –3.43265 | 2.92371 | 0.49296 |
| H | 4.03914 | –3.31156 | 0.42456 |
| H | 1.63397 | –2.52491 | 0.57134 |
| H | –0.78302 | 1.80724 | 0.14877 |
| H | 2.53129 | 3.02751 | –0.25210 |
| H | 0.22636 | 3.17603 | –0.24454 |
| H | –0.71953 | –0.68614 | –0.03250 |
| H | –7.18180 | –0.12861 | –0.02879 |

**Table S13.** Optimized geometry coordinates of A:PRF (W-edge)
Energy: –683069.1560738 (au)

| | | | |
|---|---|---|---|
| N | –6.36432 | –0.52413 | 0.25505 |
| C | –6.01227 | –1.82218 | –0.05759 |
| N | –4.72438 | –1.98038 | –0.24466 |
| C | –4.20036 | –0.71507 | –0.04460 |
| C | –2.87564 | –0.22002 | –0.09689 |

| | | | |
|---|---|---|---|
| N | –1.82370 | –1.00216 | –0.38551 |
| N | –2.68581 | 1.09727 | 0.15406 |
| C | –3.75401 | 1.86453 | 0.44265 |
| N | –5.03705 | 1.51682 | 0.52402 |
| C | –5.19935 | 0.20943 | 0.26809 |
| N | 2.34324 | 2.05473 | –0.22843 |
| C | 1.10771 | 1.45354 | –0.25780 |
| N | 0.96276 | 0.14020 | –0.26029 |
| C | 2.14118 | –0.54505 | –0.21799 |
| C | 3.44180 | –0.03079 | –0.16459 |
| C | 3.61226 | 1.39294 | –0.20922 |
| O | 4.63317 | 2.06627 | –0.24808 |
| C | 4.35787 | –1.13688 | –0.13574 |
| C | 5.85303 | –1.06456 | –0.01293 |
| N | 6.24917 | –0.86654 | 1.39164 |
| C | 3.58509 | –2.27013 | –0.17512 |
| N | 2.24083 | –1.90640 | –0.22361 |
| N | 0.02014 | 2.26519 | –0.33110 |
| H | –6.75450 | –2.60543 | –0.13186 |
| H | –1.99870 | –1.97711 | –0.56887 |
| H | –3.53165 | 2.91232 | 0.63455 |
| H | 6.29190 | –1.96351 | –0.48102 |
| H | 6.20964 | –0.19612 | –0.57446 |
| H | 3.87207 | –3.31144 | –0.17295 |
| H | 1.45753 | –2.53846 | –0.25428 |
| H | –0.90404 | 1.84733 | –0.15456 |
| H | –7.28925 | –0.16685 | 0.44136 |
| H | –0.86617 | –0.63422 | –0.36788 |
| H | 0.12996 | 3.22689 | –0.04889 |
| H | 2.41560 | 3.06234 | –0.29292 |
| H | 7.25704 | –0.73564 | 1.44476 |
| H | 6.03516 | –1.70714 | 1.92393 |

**Table S14.** Optimized geometry coordinates of A:RBF (W-edge)
Energy: –1128023.6605379 (au)

| | | | |
|---|---|---|---|
| N | 8.37721 | –0.11907 | 0.08206 |
| C | 8.67241 | 1.22556 | 0.18896 |
| N | 7.60941 | 1.99248 | 0.18895 |
| C | 6.55566 | 1.10196 | 0.07569 |
| C | 5.15512 | 1.29113 | 0.02104 |
| N | 4.57654 | 2.50215 | 0.07335 |
| N | 4.38220 | 0.18649 | –0.09188 |
| C | 4.95996 | –1.03039 | –0.14556 |
| N | 6.25747 | –1.32756 | –0.10320 |
| C | 7.00604 | –0.21953 | 0.00764 |
| N | –0.51350 | –0.94562 | –0.40156 |

| | | | |
|---|---|---|---|
| C | 0.87555 | –0.94858 | –0.32091 |
| O | 1.51345 | –1.99115 | –0.32663 |
| N | 1.54155 | 0.28010 | –0.22435 |
| C | 0.96014 | 1.51368 | –0.09978 |
| O | 1.60514 | 2.55111 | 0.02102 |
| C | –0.53637 | 1.48926 | –0.12551 |
| N | –1.19124 | 2.60609 | 0.00971 |
| C | –2.55611 | 2.55649 | –0.01840 |
| C | –3.27377 | 3.75701 | 0.17342 |
| C | –4.65429 | 3.79566 | 0.15261 |
| C | –5.39223 | 5.09448 | 0.36059 |
| C | –5.36276 | 2.57527 | –0.05607 |
| C | –6.87002 | 2.56303 | –0.04996 |
| C | –4.67281 | 1.38380 | –0.25257 |
| C | –3.26864 | 1.34709 | –0.24763 |
| N | –2.53497 | 0.18266 | –0.47249 |
| C | –1.16548 | 0.19389 | –0.32551 |
| C | –3.18466 | –1.02372 | –1.03887 |
| C | –3.76488 | –2.06356 | –0.03370 |
| O | –4.36984 | –1.40889 | 1.07011 |
| C | –2.80396 | –3.14365 | 0.49409 |
| O | –1.86260 | –2.57530 | 1.40176 |
| C | –2.17072 | –4.04319 | –0.59169 |
| O | –3.19146 | –4.88930 | –1.12154 |
| C | –1.00387 | –4.86024 | –0.03116 |
| O | –1.40601 | –5.58242 | 1.13285 |
| H | 9.69347 | 1.57469 | 0.26329 |
| H | 5.16251 | 3.31452 | 0.17581 |
| H | 4.26195 | –1.85968 | –0.23325 |
| H | –2.68597 | 4.65406 | 0.33936 |
| H | –4.69366 | 5.92225 | 0.50097 |
| H | –6.03250 | 5.33659 | –0.49566 |
| H | –6.04383 | 5.05216 | 1.24119 |
| H | –7.26715 | 2.93184 | 0.90287 |
| H | –7.27626 | 3.21410 | –0.83298 |
| H | –7.26069 | 1.55616 | –0.21037 |
| H | –5.23366 | 0.46379 | –0.33742 |
| H | –3.99900 | –0.65165 | –1.66145 |
| H | –2.44809 | –1.49077 | –1.69269 |
| H | –4.55549 | –2.58856 | –0.58190 |
| H | –3.42945 | –3.80090 | 1.10924 |
| H | –1.79405 | –3.45217 | –1.43328 |
| H | –3.30382 | –5.59479 | –0.46513 |
| H | –0.71571 | –5.60420 | –0.78039 |
| H | –0.13814 | –4.21584 | 0.15939 |
| H | –1.41171 | –4.93333 | 1.85177 |

| | | | |
|---|---|---|---|
| H | 2.58958 | 0.23610 | –0.18276 |
| H | 3.56121 | 2.59387 | 0.05729 |
| H | –3.66330 | –1.34225 | 1.73487 |
| H | 9.02617 | –0.89123 | 0.06214 |
| H | –1.21049 | –2.09393 | 0.84410 |

**Table S15.** Optimized geometry coordinates of A:SAM (W-edge)
Energy: –1352535.9857483 (au)

| | | | |
|---|---|---|---|
| N | –4.51391 | 4.23758 | –2.20071 |
| C | –4.64625 | 4.93682 | –1.01649 |
| N | –4.56331 | 4.17623 | 0.04814 |
| C | –4.36711 | 2.90822 | –0.46348 |
| C | –4.22100 | 1.65262 | 0.16508 |
| N | –4.22013 | 1.51484 | 1.50674 |
| N | –4.04463 | 0.56375 | –0.62501 |
| C | –4.04338 | 0.73552 | –1.96734 |
| N | –4.17354 | 1.85600 | –2.66739 |
| C | –4.33452 | 2.91899 | –1.86157 |
| N | –4.43154 | –2.51079 | –0.18666 |
| C | –4.04899 | –2.92422 | 1.16230 |
| C | –4.97458 | –2.31530 | 2.23581 |
| O | –5.98639 | –3.07683 | 2.68660 |
| O | –4.85809 | –1.17413 | 2.62636 |
| C | –2.60668 | –2.45417 | 1.44120 |
| C | –1.64633 | –2.99826 | 0.37827 |
| S | –0.17652 | –1.89834 | 0.21442 |
| C | –0.78138 | –0.63209 | –0.94815 |
| C | 0.95091 | –2.93609 | –0.82146 |
| C | 1.98685 | –2.06585 | –1.53786 |
| O | 2.38545 | –1.03267 | –0.63994 |
| C | 3.27232 | –2.83641 | –1.95385 |
| O | 3.63440 | –2.63550 | –3.30919 |
| C | 4.34428 | –2.25603 | –0.99569 |
| O | 5.64534 | –2.34031 | –1.48538 |
| C | 3.81898 | –0.82162 | –0.78091 |
| N | 4.31242 | –0.10047 | 0.33837 |
| C | 4.61417 | –0.52783 | 1.63199 |
| N | 4.98811 | 0.43865 | 2.42446 |
| C | 4.93219 | 1.57836 | 1.63459 |
| C | 5.21989 | 2.94019 | 1.88312 |
| N | 5.62892 | 3.39145 | 3.08574 |
| N | 5.06329 | 3.82252 | 0.87608 |
| C | 4.64525 | 3.37561 | –0.31449 |
| N | 4.34133 | 2.12027 | –0.67774 |
| C | 4.51137 | 1.27630 | 0.34207 |
| H | –4.80271 | 6.00690 | –1.00793 |

| | | | |
|---|---|---|---|
| H | –4.37488 | 0.59820 | 1.91475 |
| H | –3.91937 | –0.17884 | –2.54575 |
| H | –4.09483 | –4.02123 | 1.21436 |
| H | –2.63059 | –1.35965 | 1.43613 |
| H | –2.28853 | –2.75714 | 2.44263 |
| H | –2.14603 | –3.04743 | –0.59387 |
| H | –1.24483 | –3.98069 | 0.64269 |
| H | –1.70003 | –0.21016 | –0.52977 |
| H | 0.00057 | 0.12420 | –1.01861 |
| H | –0.99287 | –1.08566 | –1.91781 |
| H | 0.34504 | –3.51403 | –1.52475 |
| H | 1.42223 | –3.61097 | –0.10129 |
| H | 1.53128 | –1.63528 | –2.44179 |
| H | 3.17174 | –3.91927 | –1.84674 |
| H | 3.58281 | –1.69457 | –3.53409 |
| H | 4.29584 | –2.78796 | –0.03747 |
| H | 5.58577 | –2.43973 | –2.44992 |
| H | 4.01106 | –0.18928 | –1.65752 |
| H | 4.54892 | –1.56857 | 1.91629 |
| H | 5.89820 | 4.35889 | 3.17157 |
| H | 4.53940 | 4.13342 | –1.08723 |
| H | –4.47902 | 2.32439 | 2.05179 |
| H | –4.55948 | 4.60924 | –3.13850 |
| H | 5.84737 | 2.74161 | 3.82337 |
| H | –5.33776 | –2.88620 | –0.45212 |
| H | –4.46953 | –1.48559 | –0.24527 |
| H | –5.94598 | –3.97459 | 2.32552 |

**Table S16.** Optimized geometry coordinates of A:SAM (S-edge)
Energy: –1663932.7559537 (au)

| | | | |
|---|---|---|---|
| O | –6.15349 | –3.78949 | 0.13748 |
| C | –4.84305 | –4.32488 | –0.00899 |
| C | –3.73937 | –3.31581 | 0.28397 |
| O | –3.79939 | –2.25096 | –0.68577 |
| C | –3.80638 | –2.61937 | 1.65652 |
| O | –3.23729 | –3.36090 | 2.70624 |
| C | –3.07843 | –1.29396 | 1.37005 |
| O | –1.69557 | –1.57394 | 1.47894 |
| C | –3.48938 | –1.01052 | –0.09258 |
| N | –4.67718 | –0.14258 | –0.15942 |
| C | –5.98991 | –0.51425 | –0.41687 |
| N | –6.83413 | 0.49079 | –0.37136 |
| C | –6.04575 | 1.58093 | –0.06525 |
| C | –6.35081 | 2.94273 | 0.14203 |
| N | –7.60330 | 3.42459 | 0.02837 |
| N | –5.35192 | 3.79232 | 0.46461 |

| | | | |
|---|---|---|---|
| C | –4.11473 | 3.30433 | 0.56698 |
| N | –3.68823 | 2.04350 | 0.38891 |
| C | –4.70373 | 1.21342 | 0.07258 |
| N | 2.43192 | 1.35706 | –3.23340 |
| C | 1.66699 | 0.12624 | –3.10229 |
| C | 0.18019 | 0.29208 | –2.68787 |
| O | –0.43748 | –0.57885 | –2.12397 |
| O | –0.38842 | 1.46612 | –3.01280 |
| C | 2.32696 | –0.86409 | –2.11156 |
| C | 3.76395 | –1.14708 | –2.52828 |
| S | 4.70424 | –2.38999 | –1.53109 |
| C | 3.61428 | –3.85137 | –1.50510 |
| C | 4.64589 | –1.84744 | 0.23442 |
| C | 5.64628 | –0.73691 | 0.53278 |
| O | 5.33887 | 0.42063 | –0.26148 |
| C | 5.58248 | –0.31122 | 2.03116 |
| O | 6.57399 | –0.91814 | 2.80987 |
| C | 5.71288 | 1.22982 | 1.94109 |
| O | 7.07655 | 1.61786 | 2.06276 |
| C | 5.15494 | 1.58889 | 0.55031 |
| N | 3.77636 | 2.03973 | 0.48612 |
| C | 3.34543 | 3.09051 | –0.31348 |
| N | 2.04340 | 3.18910 | –0.40350 |
| C | 1.57004 | 2.14643 | 0.37846 |
| C | 0.24606 | 1.70953 | 0.66722 |
| N | –0.83169 | 2.33128 | 0.20011 |
| N | 0.12084 | 0.60273 | 1.44658 |
| C | 1.21933 | –0.01165 | 1.89705 |
| N | 2.50357 | 0.30587 | 1.68495 |
| C | 2.61765 | 1.40947 | 0.92513 |
| H | –4.68730 | –5.20165 | 0.63384 |
| H | –4.75446 | –4.65172 | –1.04984 |
| H | –2.76920 | –3.82766 | 0.19837 |
| H | –4.85165 | –2.41556 | 1.91229 |
| H | –2.29077 | –3.14294 | 2.67982 |
| H | –3.38026 | –0.48088 | 2.04014 |
| H | –1.19927 | –0.72673 | 1.47095 |
| H | –2.68426 | –0.52187 | –0.65167 |
| H | –6.23929 | –1.54839 | –0.60924 |
| H | –7.77709 | 4.39614 | 0.22890 |
| H | –3.34728 | 4.02852 | 0.83532 |
| H | 2.40518 | 1.78224 | –4.15149 |
| H | 1.64323 | –0.34996 | –4.09210 |
| H | 2.30283 | –0.42723 | –1.10956 |
| H | 1.70876 | –1.76367 | –2.09059 |
| H | 4.38066 | –0.24822 | –2.47652 |

| | | | |
|---|---|---|---|
| H | 3.83574 | –1.55812 | –3.54157 |
| H | 4.14355 | –4.63646 | –0.96270 |
| H | 2.66258 | –3.62182 | –1.02563 |
| H | 3.62796 | –1.54141 | 0.48943 |
| H | 4.91668 | –2.74696 | 0.79694 |
| H | 6.66464 | –1.07049 | 0.29659 |
| H | 4.59951 | –0.55777 | 2.43791 |
| H | 7.28570 | –0.26424 | 2.91911 |
| H | 5.18210 | 1.73954 | 2.74577 |
| H | 7.55887 | 1.38702 | 1.25469 |
| H | 5.74762 | 2.40235 | 0.12220 |
| H | 4.05752 | 3.74780 | –0.79573 |
| H | –0.69493 | 3.11996 | –0.41418 |
| H | 1.03889 | –0.89411 | 2.50663 |
| H | –8.36664 | 2.79976 | –0.17443 |
| H | –1.80499 | 2.01237 | 0.36195 |
| H | 2.27333 | 2.04172 | –2.49914 |
| H | –6.43871 | –3.91114 | 1.05136 |
| H | 3.46245 | –4.16443 | –2.53957 |
| H | 0.28001 | 2.04157 | –3.41977 |

**Table S17.** Optimized geometry coordinates of A:SFG (S-edge)
Energy: –1448574.2043582 (au)

| | | | |
|---|---|---|---|
| O | –6.64241 | 3.39337 | –1.36984 |
| C | –5.55209 | 4.17757 | –0.89445 |
| C | –4.37253 | 3.27001 | –0.60569 |
| O | –4.70334 | 2.37832 | 0.47879 |
| C | –3.93316 | 2.36138 | –1.77291 |
| O | –3.05612 | 2.98647 | –2.67593 |
| C | –3.28197 | 1.18574 | –1.01659 |
| O | –1.96325 | 1.58798 | –0.71762 |
| C | –4.13871 | 1.10452 | 0.26745 |
| N | –5.22451 | 0.11837 | 0.16748 |
| C | –6.55999 | 0.34537 | –0.12630 |
| N | –7.28582 | –0.75046 | –0.14018 |
| C | –6.38484 | –1.75374 | 0.15861 |
| C | –6.52554 | –3.14964 | 0.29572 |
| N | –7.71109 | –3.77345 | 0.12400 |
| N | –5.43908 | –3.88960 | 0.59734 |
| C | –4.26867 | –3.26182 | 0.75235 |
| N | –3.99723 | –1.95277 | 0.65321 |
| C | –5.09920 | –1.23883 | 0.35385 |
| N | 4.11525 | 0.45017 | 3.39378 |
| C | 3.77243 | 1.79391 | 2.92099 |
| C | 2.28034 | 2.02813 | 3.11514 |
| O | 1.77518 | 3.02227 | 3.58414 |

| | | | |
|---|---|---|---|
| O | 1.54454 | 0.99309 | 2.63208 |
| C | 4.12721 | 2.11790 | 1.44392 |
| C | 5.62742 | 1.99713 | 1.14741 |
| C | 6.03066 | 2.26570 | –0.31748 |
| N | 5.80392 | 3.68594 | –0.62968 |
| C | 5.37502 | 1.31827 | –1.35066 |
| C | 5.72379 | –0.16249 | –1.27805 |
| O | 5.19965 | –0.77039 | –0.07004 |
| C | 5.11621 | –0.99217 | –2.43170 |
| O | 5.87626 | –0.97132 | –3.61606 |
| C | 4.99862 | –2.39253 | –1.80205 |
| O | 6.20802 | –3.13234 | –1.99643 |
| C | 4.75855 | –2.09536 | –0.30497 |
| N | 3.38708 | –2.25107 | 0.16352 |
| C | 2.99113 | –3.07674 | 1.20752 |
| N | 1.72429 | –2.97743 | 1.51659 |
| C | 1.24614 | –2.01961 | 0.64318 |
| C | –0.04263 | –1.45755 | 0.48801 |
| N | –1.07753 | –1.81154 | 1.26950 |
| N | –0.19998 | –0.51589 | –0.47165 |
| C | 0.85921 | –0.16575 | –1.22590 |
| N | 2.10677 | –0.62423 | –1.16788 |
| C | 2.25043 | –1.55079 | –0.20522 |
| H | –5.22563 | 4.91018 | –1.64738 |
| H | –5.82250 | 4.72078 | 0.02243 |
| H | –3.52010 | 3.89739 | –0.30975 |
| H | –4.81504 | 2.02763 | –2.32644 |
| H | –2.17832 | 2.87646 | –2.26960 |
| H | –3.30525 | 0.24275 | –1.57500 |
| H | –1.40202 | 0.78294 | –0.57163 |
| H | –3.51956 | 0.81351 | 1.12285 |
| H | –6.91160 | 1.34832 | –0.32524 |
| H | –7.77039 | –4.76479 | 0.28882 |
| H | –3.42212 | –3.90416 | 0.98746 |
| H | 5.12143 | 0.31787 | 3.34024 |
| H | 3.69183 | –0.24268 | 2.78078 |
| H | 4.27039 | 2.51851 | 3.57308 |
| H | 3.55673 | 1.43677 | 0.80323 |
| H | 3.78095 | 3.13766 | 1.23123 |
| H | 5.96336 | 0.99116 | 1.41606 |
| H | 6.17937 | 2.71198 | 1.76950 |
| H | 7.11827 | 2.11413 | –0.37500 |
| H | 6.17086 | 3.89677 | –1.55567 |
| H | 4.80397 | 3.87245 | –0.68084 |
| H | 5.69111 | 1.64372 | –2.35119 |
| H | 4.28331 | 1.41337 | –1.32073 |

| | | | |
|---|---|---|---|
| H | 6.82100 | –0.28500 | –1.28690 |
| H | 4.11485 | –0.61072 | –2.64720 |
| H | 6.46058 | –1.74583 | –3.57355 |
| H | 4.20800 | –3.00334 | –2.23940 |
| H | 6.91285 | –2.72195 | –1.47402 |
| H | 5.34284 | –2.80777 | 0.29284 |
| H | –0.91055 | –2.55760 | 1.92864 |
| H | –2.04982 | –1.62460 | 1.00078 |
| H | 0.65690 | 0.59683 | –1.97413 |
| H | 3.70258 | –3.73400 | 1.69071 |
| H | –7.37767 | 3.98423 | –1.57207 |
| H | –8.54248 | –3.23083 | –0.04207 |
| H | 0.61225 | 1.23058 | 2.77090 |

**Table S18.** Optimized geometry coordinates of G:PYI (W-edge)
Energy: –1579476.8320853 (au)

| | | | |
|---|---|---|---|
| N | 5.14839 | 4.59323 | –0.09720 |
| C | 4.23912 | 4.60455 | –1.14610 |
| N | 3.85529 | 3.41067 | –1.50863 |
| C | 4.54626 | 2.56055 | –0.66141 |
| C | 4.58071 | 1.15213 | –0.57763 |
| O | 3.96877 | 0.30915 | –1.30097 |
| N | 5.41578 | 0.69437 | 0.43766 |
| C | 6.15153 | 1.49686 | 1.28107 |
| N | 6.89359 | 0.85615 | 2.20966 |
| N | 6.16112 | 2.81704 | 1.20372 |
| C | 5.35858 | 3.28364 | 0.23243 |
| C | –9.04506 | 3.22370 | 0.10959 |
| C | –8.15197 | 2.01810 | 0.09838 |
| N | –7.22809 | 1.94389 | –0.87862 |
| C | –6.44395 | 0.86607 | –0.86155 |
| C | –6.53818 | –0.15345 | 0.08295 |
| C | –5.61668 | –1.34955 | 0.12876 |
| N | –4.26926 | –1.07624 | –0.45666 |
| C | –4.05162 | –1.50238 | –1.72700 |
| C | –2.84914 | –1.26772 | –2.35580 |
| C | –1.85828 | –0.58108 | –1.65919 |
| C | –2.07516 | –0.13747 | –0.34795 |
| C | –0.93607 | 0.55725 | 0.36902 |
| C | –0.10395 | –0.40132 | 1.24379 |
| O | 0.17341 | –1.63397 | 0.55891 |
| P | 1.43838 | –1.85930 | –0.40691 |
| O | 1.36547 | –3.17319 | –1.08188 |
| O | 1.48192 | –0.52475 | –1.27496 |
| O | 2.72441 | –1.72788 | 0.58292 |
| P | 4.15121 | –2.53353 | 0.28146 |

| | | | |
|---|---|---|---|
| O | 3.78264 | –4.06834 | 0.36945 |
| O | 5.16132 | –2.03480 | 1.24900 |
| O | 4.43946 | –2.25933 | –1.25119 |
| C | –3.31402 | –0.40133 | 0.26045 |
| C | –3.64359 | –0.00491 | 1.67077 |
| C | –7.57726 | 0.00024 | 1.03449 |
| N | –7.80417 | –0.91673 | 2.03142 |
| N | –8.35505 | 1.09034 | 1.04130 |
| H | 3.90797 | 5.53265 | –1.59153 |
| H | 6.79807 | –0.13723 | 2.36047 |
| H | –9.47381 | 3.37690 | 1.10048 |
| H | –9.86836 | 3.07638 | –0.59950 |
| H | –8.49333 | 4.10868 | –0.21288 |
| H | –5.68285 | 0.82703 | –1.64086 |
| H | –6.01174 | –2.20890 | –0.42246 |
| H | –5.46956 | –1.67817 | 1.16000 |
| H | –4.87015 | –2.02800 | –2.20174 |
| H | –2.69279 | –1.62217 | –3.36744 |
| H | –0.89641 | –0.39109 | –2.12540 |
| H | –1.29184 | 1.37844 | 0.99867 |
| H | –0.28056 | 0.99545 | –0.38487 |
| H | –0.65225 | –0.69590 | 2.14207 |
| H | 0.82149 | 0.09037 | 1.55747 |
| H | –4.60410 | 0.51280 | 1.72892 |
| H | –3.69514 | –0.88540 | 2.32220 |
| H | –2.87984 | 0.65519 | 2.07465 |
| H | –8.62829 | –0.71559 | 2.58424 |
| H | –7.66193 | –1.89791 | 1.84342 |
| H | 2.39124 | –0.10490 | –1.37804 |
| H | 3.03083 | –4.27488 | –0.22495 |
| H | 4.42552 | –1.27430 | –1.43502 |
| H | 7.38711 | 1.41618 | 2.88573 |
| H | 5.42292 | –0.31829 | 0.62972 |
| H | 5.59397 | 5.38765 | 0.33913 |

**Table S19.** Optimized geometry coordinates of G:RS3 (W-edge)
Energy: –1234629.1230812 (au)

| | | | |
|---|---|---|---|
| N | –8.40125 | 0.87517 | –1.06339 |
| C | –7.82052 | 1.91917 | –1.76725 |
| N | –6.51851 | 1.94065 | –1.66543 |
| C | –6.21780 | 0.85957 | –0.85738 |
| C | –4.96075 | 0.37459 | –0.37459 |
| O | –3.82311 | 0.82442 | –0.56421 |
| N | –5.14370 | –0.78035 | 0.41809 |
| C | –6.34678 | –1.35855 | 0.73964 |
| N | –6.27386 | –2.48353 | 1.50954 |

| | | | |
|---|---|---|---|
| N | –7.51096 | –0.90970 | 0.31981 |
| C | –7.38018 | 0.18166 | –0.47071 |
| O | 5.08870 | 4.76745 | 0.11485 |
| C | 4.04430 | 4.15600 | 0.24627 |
| N | 2.83101 | 4.78088 | 0.47246 |
| C | 1.55823 | 4.18464 | 0.61906 |
| O | 0.58281 | 4.89489 | 0.77885 |
| C | 3.91783 | 2.66699 | 0.18735 |
| C | 2.58989 | 2.10704 | 0.39497 |
| N | 1.49122 | 2.80118 | 0.57308 |
| N | 4.98358 | 1.95532 | –0.05354 |
| C | 4.85456 | 0.60040 | –0.11595 |
| C | 5.99606 | –0.18067 | –0.39879 |
| C | 5.96266 | –1.55805 | –0.47701 |
| C | 7.24749 | –2.32664 | –0.67261 |
| C | 4.69974 | –2.21114 | –0.27778 |
| N | 4.62357 | –3.61305 | –0.32562 |
| C | 3.51536 | –4.26463 | 0.36396 |
| C | 4.94093 | –4.27166 | –1.59430 |
| C | 3.55995 | –1.45488 | 0.02907 |
| C | 3.61704 | –0.05748 | 0.12119 |
| N | 2.51023 | 0.72677 | 0.42202 |
| C | 1.22014 | 0.10942 | 0.77317 |
| C | 0.34590 | –0.18138 | –0.45824 |
| O | 0.10908 | 0.98397 | –1.21672 |
| C | –0.98938 | –0.84137 | –0.05555 |
| O | –1.70880 | –0.01439 | 0.86430 |
| C | –0.84324 | –2.23821 | 0.58049 |
| O | –0.02384 | –3.10087 | –0.20877 |
| C | –2.20251 | –2.90236 | 0.85045 |
| O | –3.00026 | –2.14998 | 1.77459 |
| H | –8.41933 | 2.62172 | –2.33042 |
| H | –5.42565 | –2.65206 | 2.03349 |
| H | 6.93023 | 0.35833 | –0.52222 |
| H | 8.09566 | –1.73751 | –0.31492 |
| H | 7.22416 | –3.27154 | –0.12269 |
| H | 7.44516 | –2.56241 | –1.72540 |
| H | 3.74790 | –5.32995 | 0.45537 |
| H | 3.40892 | –3.85143 | 1.36956 |
| H | 2.54828 | –4.17205 | –0.15526 |
| H | 5.75898 | –3.76881 | –2.10555 |
| H | 5.23700 | –5.30862 | –1.40657 |
| H | 4.07226 | –4.27815 | –2.27252 |
| H | 2.61498 | –1.96643 | 0.13745 |
| H | 1.43249 | –0.80002 | 1.33679 |
| H | 0.69457 | 0.80709 | 1.42509 |

| | | | |
|---|---|---|---|
| H | 0.87096 | –0.88121 | –1.11923 |
| H | 0.05446 | 1.74064 | –0.60612 |
| H | –1.57092 | –0.93741 | –0.98574 |
| H | –2.42352 | 0.44913 | 0.35427 |
| H | –0.32448 | –2.14256 | 1.54136 |
| H | –0.40695 | –3.14882 | –1.09657 |
| H | –2.03368 | –3.90147 | 1.26513 |
| H | –2.74354 | –3.02166 | –0.10268 |
| H | –2.59059 | –1.25402 | 1.79455 |
| H | 2.84320 | 5.79309 | 0.51438 |
| H | –7.14468 | –2.79105 | 1.91356 |
| H | –4.30294 | –1.21767 | 0.80602 |
| H | –9.38234 | 0.65284 | –0.98623 |

**Table S20.** Optimized geometry coordinates of G:SAH (S-edge)
Energy: –1686247.6839170 (au)

| | | | |
|---|---|---|---|
| O | 4.64270 | 4.24313 | 1.11092 |
| C | 3.25460 | 4.50243 | 0.95605 |
| C | 2.48056 | 3.20400 | 1.07955 |
| O | 2.74368 | 2.35370 | –0.06807 |
| C | 2.81771 | 2.34210 | 2.31568 |
| O | 2.08598 | 2.69741 | 3.46375 |
| C | 2.49349 | 0.92320 | 1.80667 |
| O | 1.10398 | 0.66101 | 1.97677 |
| C | 2.89442 | 0.98364 | 0.32180 |
| N | 4.24289 | 0.52884 | 0.03683 |
| C | 5.45875 | 1.20061 | 0.16256 |
| N | 6.48235 | 0.46232 | –0.18146 |
| C | 5.93624 | –0.75386 | –0.54993 |
| C | 6.56009 | –1.95826 | –1.03246 |
| O | 7.73032 | –2.23479 | –1.23684 |
| N | 5.55040 | –2.94854 | –1.29859 |
| C | 4.19193 | –2.81019 | –1.13598 |
| N | 3.40305 | –3.85809 | –1.50827 |
| N | 3.63968 | –1.70380 | –0.67983 |
| C | 4.54882 | –0.73196 | –0.42656 |
| N | –0.84403 | –1.56454 | 0.95649 |
| C | –0.95346 | –2.98810 | 1.22320 |
| C | –2.35130 | –3.50686 | 0.79296 |
| C | –3.46685 | –2.87199 | 1.62671 |
| S | –5.16126 | –3.20482 | 0.99099 |
| C | 0.11394 | –3.81657 | 0.49210 |
| O | 0.44942 | –5.01371 | 1.02713 |
| O | 0.61985 | –3.45987 | –0.54930 |
| C | –5.33692 | –1.92290 | –0.31438 |
| C | –5.66221 | –0.52108 | 0.17631 |

| | | | |
|---|---|---|---|
| O | –4.53819 | 0.03027 | 0.89803 |
| C | –5.94501 | 0.49769 | –0.94915 |
| O | –7.26989 | 0.47400 | –1.41857 |
| C | –5.55135 | 1.82958 | –0.27877 |
| O | –6.64755 | 2.34660 | 0.47925 |
| C | –4.38668 | 1.42039 | 0.64422 |
| N | –3.04304 | 1.69223 | 0.13411 |
| C | –1.93017 | 1.72119 | 0.94136 |
| N | –0.80173 | 1.87217 | 0.28819 |
| C | –1.18082 | 1.93393 | –1.04792 |
| C | –0.46133 | 2.11537 | –2.25648 |
| N | 0.87069 | 2.28744 | –2.31887 |
| N | –1.15612 | 2.11336 | –3.41516 |
| C | –2.48266 | 1.96107 | –3.37423 |
| N | –3.27451 | 1.81086 | –2.30470 |
| C | –2.56837 | 1.80521 | –1.16819 |
| H | 3.03573 | 4.95768 | –0.02073 |
| H | 2.88221 | 5.17921 | 1.74014 |
| H | 1.40906 | 3.44538 | 1.09536 |
| H | 3.88255 | 2.43479 | 2.54553 |
| H | 1.27567 | 2.16158 | 3.41608 |
| H | 3.03293 | 0.13926 | 2.34083 |
| H | 0.57310 | 1.16093 | 1.30817 |
| H | 2.23156 | 0.34930 | –0.27417 |
| H | 5.48991 | 2.23380 | 0.47909 |
| H | 2.42627 | –3.79342 | –1.22778 |
| H | –0.03040 | –1.15847 | 1.41755 |
| H | –0.68906 | –1.43981 | –0.04286 |
| H | –0.83581 | –3.14525 | 2.30555 |
| H | –2.48009 | –3.26211 | –0.26800 |
| H | –2.39703 | –4.60001 | 0.86656 |
| H | –3.32299 | –1.79421 | 1.69104 |
| H | –3.46356 | –3.27753 | 2.64422 |
| H | –6.16876 | –2.26274 | –0.93935 |
| H | –4.43798 | –1.90008 | –0.93873 |
| H | –6.53346 | –0.57205 | 0.84777 |
| H | –5.27268 | 0.32050 | –1.79405 |
| H | –7.75665 | 1.11738 | –0.87780 |
| H | –5.28391 | 2.60040 | –0.99918 |
| H | –6.80764 | 1.76596 | 1.23785 |
| H | –4.45598 | 1.96916 | 1.59103 |
| H | –2.01434 | 1.60381 | 2.01281 |
| H | –2.98537 | 1.96457 | –4.33881 |
| H | 5.13283 | 5.05612 | 0.93982 |
| H | 5.92766 | –3.80126 | –1.69339 |
| H | 3.80259 | –4.78390 | –1.50964 |

| | | | |
|---|---|---|---|
| H | –0.01455 | –5.14914 | 1.86651 |
| H | 1.46552 | 2.36947 | –1.49784 |
| H | 1.27258 | 2.40838 | –3.23577 |

**Table S21.** Optimized geometry coordinates of G:2QB (S-edge)
Energy: –1004475.9216297 (au)

| | | | |
|---|---|---|---|
| O | –5.20774 | –1.75128 | –0.56394 |
| C | –4.29757 | –2.84526 | –0.57744 |
| C | –2.94012 | –2.52209 | 0.03770 |
| O | –2.23733 | –1.58634 | –0.80689 |
| C | –2.97301 | –1.86964 | 1.43345 |
| O | –3.07805 | –2.78498 | 2.49470 |
| C | –1.65535 | –1.06903 | 1.43895 |
| O | –0.58930 | –1.90078 | 1.88366 |
| C | –1.51805 | –0.63940 | –0.03780 |
| N | –2.03137 | 0.70136 | –0.30571 |
| C | –3.34844 | 1.09542 | –0.53778 |
| N | –3.48276 | 2.39504 | –0.61512 |
| C | –2.20886 | 2.89476 | –0.42393 |
| C | –1.73364 | 4.25328 | –0.36297 |
| O | –2.31625 | 5.31679 | –0.47382 |
| N | –0.31460 | 4.25916 | –0.10049 |
| C | 0.50269 | 3.16794 | 0.04096 |
| N | 1.81835 | 3.37780 | 0.31674 |
| N | 0.04976 | 1.92855 | –0.04537 |
| C | –1.29538 | 1.86055 | –0.23897 |
| N | 1.00020 | –3.62144 | –0.13900 |
| N | 1.63213 | –3.21477 | –1.00155 |
| N | 2.17631 | –2.79194 | –2.02071 |
| C | 3.59590 | –2.32730 | –1.90941 |
| C | 3.86210 | –1.31772 | –0.82595 |
| C | 4.89229 | –1.48876 | 0.08487 |
| N | 5.21890 | –0.60221 | 1.03656 |
| C | 4.47604 | 0.51096 | 1.06811 |
| C | 4.83911 | 1.55476 | 2.08908 |
| N | 3.44389 | 0.78893 | 0.25785 |
| C | 3.10902 | –0.11848 | –0.68252 |
| N | 2.04743 | 0.17590 | –1.47505 |
| H | –4.15107 | –3.11713 | –1.62726 |
| H | –4.71003 | –3.71948 | –0.05615 |
| H | –2.36519 | –3.46050 | 0.09553 |
| H | –3.82103 | –1.17850 | 1.49036 |
| H | –2.16002 | –3.00183 | 2.73304 |
| H | –1.67342 | –0.21123 | 2.11333 |
| H | –0.28625 | –2.48217 | 1.16736 |
| H | –0.46152 | –0.62930 | –0.32565 |

| | | | |
|---|---|---|---|
| H | –4.13078 | 0.36000 | –0.66909 |
| H | 2.24234 | 4.24716 | 0.03198 |
| H | 3.79657 | –1.91216 | –2.90111 |
| H | 5.49640 | –2.39604 | 0.04908 |
| H | 5.43152 | 1.10696 | 2.88748 |
| H | 5.44135 | 2.34266 | 1.62048 |
| H | 1.67668 | –0.55863 | –2.06105 |
| H | 1.37101 | 0.84755 | –1.09886 |
| H | 3.94277 | 2.02400 | 2.50168 |
| H | –5.68565 | –1.76068 | 0.27458 |
| H | 0.06076 | 5.19083 | 0.02875 |
| H | 2.41657 | 2.54396 | 0.32035 |
| H | 4.25027 | –3.19719 | –1.77968 |

**Table S22.** Optimized geometry coordinates of G:G6P (W-edge)
Energy: –1439336.4093151 (au)

| | | | |
|---|---|---|---|
| O | –2.73018 | 2.52943 | –0.83197 |
| C | –3.94538 | 1.96366 | –0.39489 |
| C | –4.28086 | 0.77051 | –1.29262 |
| O | –3.31037 | –0.29637 | –1.22068 |
| C | –5.59345 | 0.05767 | –0.97183 |
| O | –6.08879 | –0.66612 | –2.10175 |
| C | –5.18513 | –0.95639 | 0.13070 |
| O | –5.99419 | –2.10262 | 0.14063 |
| C | –3.69810 | –1.24829 | –0.22861 |
| N | –2.77195 | –1.20599 | 0.90102 |
| C | –2.77829 | –0.36491 | 2.01490 |
| N | –1.62756 | –0.28781 | 2.62459 |
| C | –0.79663 | –1.10938 | 1.88595 |
| C | 0.61538 | –1.29095 | 1.96539 |
| O | 1.43450 | –0.72564 | 2.70289 |
| N | 1.07038 | –2.19706 | 0.98238 |
| C | 0.29263 | –2.74562 | –0.01402 |
| N | 0.91508 | –3.58917 | –0.88386 |
| N | –0.99938 | –2.51036 | –0.13834 |
| C | –1.48176 | –1.68847 | 0.81799 |
| C | 0.58823 | 2.76316 | –1.05410 |
| C | 0.26961 | 2.41070 | 0.42370 |
| C | 1.57292 | 2.35532 | 1.24045 |
| C | 2.50577 | 1.36819 | 0.52161 |
| C | 2.83440 | 1.88562 | –0.87778 |
| C | 3.80210 | 1.01753 | –1.67347 |
| O | 0.84904 | 4.15904 | –1.16833 |
| O | –0.68208 | 3.34402 | 0.92460 |
| O | 1.45958 | 2.04105 | 2.60791 |
| O | 3.68866 | 1.14808 | 1.29629 |

| | | | |
|---|---|---|---|
| O | 1.60689 | 1.96070 | –1.61211 |
| O | 3.46317 | –0.38439 | –1.57905 |
| P | 4.48360 | –1.40048 | –0.84156 |
| O | 4.00378 | –1.45303 | 0.68372 |
| O | 3.89208 | –2.83129 | –1.33231 |
| O | 5.91465 | –1.17767 | –1.13114 |
| H | –4.77736 | 2.68465 | –0.47691 |
| H | –3.89269 | 1.65332 | 0.65616 |
| H | –4.26877 | 1.12622 | –2.32866 |
| H | –6.40157 | 0.72057 | –0.65377 |
| H | –5.31936 | –0.97413 | –2.60491 |
| H | –5.26499 | –0.49010 | 1.11684 |
| H | –6.36158 | –2.15850 | –0.75923 |
| H | –3.59715 | –2.24582 | –0.66057 |
| H | –3.67588 | 0.13428 | 2.34730 |
| H | 0.36033 | –3.81500 | –1.69639 |
| H | –0.29850 | 2.58991 | –1.66545 |
| H | –0.21778 | 1.43173 | 0.45291 |
| H | 2.04042 | 3.35207 | 1.22744 |
| H | 1.97838 | 0.41765 | 0.38835 |
| H | 3.30851 | 2.88025 | –0.80663 |
| H | 3.73475 | 1.29425 | –2.72826 |
| H | 4.82638 | 1.18108 | –1.32893 |
| H | 1.70532 | 4.35974 | –0.76486 |
| H | –0.41808 | 4.20709 | 0.56266 |
| H | 1.14544 | 1.11786 | 2.72753 |
| H | 3.39814 | 1.28408 | 2.21987 |
| H | –2.23027 | 2.88388 | –0.07294 |
| H | 3.96432 | –0.54168 | 1.09094 |
| H | 2.08299 | –2.31249 | 0.96298 |
| H | 1.91118 | –3.47981 | –1.04120 |
| H | 4.52542 | –3.25278 | –1.93129 |

**Table S23.** Optimized geometry coordinates of G:BFT (S-edge)
Energy: –2000454.3551919 (au)

| | | | |
|---|---|---|---|
| O | –5.07275 | 4.71005 | 2.50374 |
| C | –3.76326 | 4.90244 | 1.98624 |
| C | –3.55104 | 3.97821 | 0.80131 |
| O | –3.47906 | 2.60643 | 1.24447 |
| C | –4.65707 | 4.01399 | –0.27637 |
| O | –4.49237 | 5.03730 | –1.22646 |
| C | –4.54505 | 2.59823 | –0.87361 |
| O | –3.53493 | 2.58535 | –1.88665 |
| C | –4.15235 | 1.73796 | 0.34425 |
| N | –5.28421 | 1.10656 | 1.01247 |
| C | –6.16959 | 1.67301 | 1.92930 |

| | | | |
|---|---|---|---|
| N | –7.14585 | 0.86660 | 2.25673 |
| C | –6.91274 | –0.28465 | 1.52961 |
| C | –7.66953 | –1.50689 | 1.45034 |
| O | –8.69912 | –1.85401 | 2.00110 |
| N | –7.03421 | –2.40856 | 0.52410 |
| C | –5.87040 | –2.19675 | –0.17046 |
| N | –5.44658 | –3.17403 | –1.01433 |
| N | –5.18051 | –1.07193 | –0.06989 |
| C | –5.75971 | –0.16033 | 0.75831 |
| C | –3.10638 | –4.59118 | –3.74214 |
| N | 2.08074 | –2.24917 | –0.56344 |
| C | 2.73505 | –3.35284 | –0.07532 |
| S | 3.67202 | –0.22492 | 1.33266 |
| C | 2.96290 | 0.09362 | –0.35749 |
| C | 2.81975 | –1.17940 | –1.24818 |
| C | 2.18220 | –0.85021 | –2.60881 |
| C | 3.56403 | 1.34060 | –1.03813 |
| C | 4.99120 | 1.23550 | –1.57468 |
| O | 5.30070 | 2.42468 | –2.35309 |
| N | –0.82437 | –4.31611 | –2.95397 |
| C | –2.10029 | –3.94971 | –2.82352 |
| N | –2.57481 | –3.07329 | –1.92134 |
| C | –1.69316 | –2.48496 | –1.09137 |
| N | –2.18397 | –1.55697 | –0.21735 |
| C | –0.30793 | –2.78751 | –1.16090 |
| C | 0.04398 | –3.73271 | –2.10680 |
| C | 0.66748 | –2.10415 | –0.22319 |
| P | 5.75133 | 3.77874 | –1.61634 |
| O | 4.85480 | 4.40365 | –0.62450 |
| O | 6.04780 | 4.70952 | –2.89118 |
| O | 7.19708 | 3.34723 | –1.01428 |
| O | 2.19950 | –4.24331 | 0.56266 |
| C | 5.23993 | –1.06635 | 1.03633 |
| O | 5.66810 | –1.27038 | –0.08929 |
| C | 5.98461 | –1.48266 | 2.26045 |
| C | 7.29531 | –1.95344 | 2.07897 |
| C | 8.05375 | –2.35383 | 3.17426 |
| C | 7.50995 | –2.29437 | 4.45987 |
| C | 6.20464 | –1.83446 | 4.64664 |
| C | 5.44233 | –1.42949 | 3.55359 |
| H | –3.62050 | 5.93340 | 1.62901 |
| H | –2.99533 | 4.68980 | 2.74348 |
| H | –2.59260 | 4.25962 | 0.33594 |
| H | –5.62791 | 4.15083 | 0.20716 |
| H | –3.93878 | 4.65265 | –1.92677 |
| H | –5.45935 | 2.24818 | –1.35427 |

| | | | |
|---|---|---|---|
| H | –2.66540 | 2.56421 | –1.46260 |
| H | –3.48671 | 0.92837 | 0.02604 |
| H | –5.99816 | 2.66619 | 2.32055 |
| H | –4.50298 | –3.07354 | –1.40943 |
| H | –3.81894 | –3.85145 | –4.11681 |
| H | –2.59446 | –5.07275 | –4.57565 |
| H | –3.67828 | –5.35772 | –3.20516 |
| H | 3.80947 | –3.35613 | –0.33214 |
| H | 3.81911 | –1.57930 | –1.42636 |
| H | 2.05037 | –1.76903 | –3.18413 |
| H | 2.81700 | –0.17880 | –3.19147 |
| H | 1.20042 | –0.37729 | –2.50220 |
| H | 2.91406 | 1.61481 | –1.87646 |
| H | 3.52005 | 2.17715 | –0.33541 |
| H | 5.11764 | 0.40506 | –2.27027 |
| H | 5.72969 | 1.11727 | –0.78059 |
| H | –3.19889 | –1.42500 | –0.18075 |
| H | 1.08218 | –4.03776 | –2.20680 |
| H | 0.42529 | –1.03603 | –0.18355 |
| H | 0.54151 | –2.50586 | 0.79059 |
| H | 7.69731 | –1.99457 | 1.07285 |
| H | 9.06742 | –2.71374 | 3.02731 |
| H | 8.10090 | –2.60911 | 5.31487 |
| H | 5.77785 | –1.79600 | 5.64390 |
| H | 4.42459 | –1.08798 | 3.70980 |
| H | –5.17136 | 5.23838 | 3.30473 |
| H | –7.55868 | –3.26277 | 0.37993 |
| H | –5.76152 | –4.11782 | –0.84971 |
| H | –1.68159 | –1.38841 | 0.63990 |
| H | 7.38026 | 3.85387 | –0.20970 |
| H | 6.50939 | 4.23469 | –3.59753 |
| H | 1.95112 | 0.39438 | –0.05975 |

**Table S24.** Optimized geometry coordinates of G:EEM (S-edge)
Energy: –2966926.0045106 (au)

| | | | |
|---|---|---|---|
| O | –9.12237 | –1.49567 | –0.56968 |
| C | –8.34949 | –2.66402 | –0.79301 |
| C | –6.88007 | –2.46977 | –0.43566 |
| O | –6.35429 | –1.34150 | –1.16923 |
| C | –6.59313 | –2.14910 | 1.04392 |
| O | –6.56056 | –3.27209 | 1.88608 |
| C | –5.25502 | –1.39840 | 0.93370 |
| O | –4.16221 | –2.33253 | 0.87904 |
| C | –5.40925 | –0.62141 | –0.39049 |
| N | –5.84898 | 0.75675 | –0.21069 |
| C | –7.14991 | 1.23788 | –0.06499 |

| | | | |
|---|---|---|---|
| N | –7.20443 | 2.54036 | 0.01653 |
| C | –5.89227 | 2.95466 | –0.07363 |
| C | –5.33721 | 4.28388 | –0.05948 |
| O | –5.85967 | 5.37560 | 0.04501 |
| N | –3.90211 | 4.20478 | –0.19708 |
| C | –3.14306 | 3.07051 | –0.28849 |
| N | –1.78738 | 3.22321 | –0.31656 |
| N | –3.67192 | 1.85866 | –0.29996 |
| C | –5.03370 | 1.86680 | –0.20394 |
| C | 1.30664 | 0.65734 | –0.73837 |
| N | 2.17964 | –0.35648 | –0.63793 |
| C | 1.56459 | –1.49077 | –0.23382 |
| C | 3.43420 | –3.23921 | –0.32106 |
| C | 3.99405 | –2.91190 | –1.73542 |
| C | 5.50100 | –2.70153 | –1.45385 |
| C | 5.49537 | –2.08242 | –0.05216 |
| C | 5.33747 | –0.55678 | –0.06903 |
| O | 7.08300 | 4.13008 | –1.56516 |
| C | 6.55200 | 3.29725 | –0.68276 |
| O | 7.02736 | 2.20517 | –0.43338 |
| C | 5.28320 | 3.84873 | 0.02033 |
| N | 4.70702 | 4.89098 | –0.83009 |
| C | 4.32780 | 2.69823 | 0.41799 |
| C | 4.90493 | 1.85797 | 1.56879 |
| Se | 4.30772 | –0.02771 | 1.52000 |
| C | 5.57139 | –0.67437 | 2.87485 |
| O | 4.37177 | –2.69037 | 0.62994 |
| O | 6.18755 | –3.94397 | –1.50477 |
| O | 3.73736 | –3.91585 | –2.67190 |
| N | 2.09849 | –2.76075 | –0.06391 |
| C | 1.03768 | –3.55625 | 0.36136 |
| N | –0.09392 | –2.91561 | 0.46393 |
| C | 0.21695 | –1.62114 | 0.09521 |
| N | 0.00060 | 0.67769 | –0.46817 |
| C | –0.59746 | –0.46461 | –0.02501 |
| N | –1.89378 | –0.45687 | 0.26944 |
| H | –8.75048 | –3.45162 | –0.14658 |
| H | –8.43557 | –3.00595 | –1.83261 |
| H | –6.33608 | –3.38753 | –0.71876 |
| H | –7.37049 | –1.47277 | 1.41433 |
| H | –5.64942 | –3.60447 | 1.87730 |
| H | –5.03899 | –0.74690 | 1.78084 |
| H | –4.22660 | –2.84589 | 0.05939 |
| H | –4.45153 | –0.56838 | –0.92002 |
| H | –7.99184 | 0.56275 | –0.02810 |
| H | –1.41902 | 4.09824 | –0.65566 |

| | | | |
|---|---|---|---|
| H | 1.72102 | 1.60118 | –1.08837 |
| H | 3.39704 | –4.32507 | –0.18753 |
| H | 3.53735 | –1.98447 | –2.09288 |
| H | 5.97887 | –2.06326 | –2.20084 |
| H | 6.39912 | –2.36239 | 0.49755 |
| H | 6.27889 | –0.00469 | –0.07669 |
| H | 4.69527 | –0.22973 | –0.88681 |
| H | 5.64289 | 4.34628 | 0.93103 |
| H | 6.46214 | 4.89605 | –1.61926 |
| H | 4.20051 | 5.59220 | –0.29987 |
| H | 4.14456 | 2.08220 | –0.46830 |
| H | 3.36335 | 3.11973 | 0.71665 |
| H | 5.99108 | 1.80940 | 1.51949 |
| H | 4.57349 | 2.22396 | 2.54352 |
| H | 6.58586 | –0.38362 | 2.60330 |
| H | 5.27753 | –0.23859 | 3.83006 |
| H | 5.45109 | –1.75638 | 2.91456 |
| H | 6.02789 | –4.44175 | –0.68946 |
| H | 4.50943 | –4.50730 | –2.66410 |
| H | 1.18572 | –4.60890 | 0.56435 |
| H | –2.46908 | 0.37774 | 0.09317 |
| H | –8.98272 | –0.91195 | –1.32688 |
| H | –3.45255 | 5.11077 | –0.14242 |
| H | –1.22304 | 2.39201 | –0.50004 |
| H | –2.35516 | –1.29927 | 0.59952 |
| H | 4.08069 | 4.51090 | –1.53571 |

**Table S25.** Optimized geometry coordinates of G:LYS (S-edge)
Energy: –963767.0108542 (au)

| | | | |
|---|---|---|---|
| O | 4.10119 | 3.08934 | –0.49485 |
| C | 2.89195 | 3.83253 | –0.50761 |
| C | 1.66380 | 3.08382 | –0.00018 |
| O | 1.27561 | 2.03414 | –0.91761 |
| C | 1.76642 | 2.41322 | 1.38329 |
| O | 1.52131 | 3.29060 | 2.45511 |
| C | 0.70855 | 1.29625 | 1.25524 |
| O | –0.57625 | 1.83059 | 1.55252 |
| C | 0.87422 | 0.86163 | –0.21293 |
| N | 1.89628 | –0.18270 | –0.38192 |
| C | 3.23282 | –0.00240 | –0.73580 |
| N | 3.90438 | –1.12359 | –0.79354 |
| C | 2.98493 | –2.09863 | –0.47060 |
| C | 3.15505 | –3.52355 | –0.36290 |
| O | 4.13629 | –4.23119 | –0.51656 |
| N | 1.90364 | –4.13064 | –0.00738 |
| C | 0.70980 | –3.48815 | 0.20421 |

| | | | |
|---|---|---|---|
| N | –0.35829 | –4.26595 | 0.56203 |
| N | 0.57413 | –2.18033 | 0.11941 |
| C | 1.73099 | –1.54237 | –0.21345 |
| N | –2.15745 | –0.66430 | 0.49355 |
| C | –3.59255 | –0.88352 | 0.48451 |
| C | –3.93386 | –2.34020 | 0.21469 |
| O | –3.13913 | –3.25890 | 0.12595 |
| C | –4.26128 | 0.06292 | –0.54852 |
| C | –3.95198 | 1.52691 | –0.19126 |
| C | –3.76437 | 2.45472 | –1.39756 |
| C | –3.03581 | 3.75215 | –1.03993 |
| N | –1.63317 | 3.49781 | –0.63375 |
| O | –5.25969 | –2.54427 | 0.08879 |
| H | –3.82941 | –0.17908 | –1.52697 |
| H | –5.33839 | –0.12140 | –0.60723 |
| H | –3.01940 | 1.54693 | 0.37541 |
| H | –4.73393 | 1.92836 | 0.46721 |
| H | –3.18413 | 1.93312 | –2.17182 |
| H | –4.72826 | 2.70597 | –1.85786 |
| H | –3.53724 | 4.24130 | –0.19614 |
| H | –3.09059 | 4.44770 | –1.89099 |
| H | –1.11171 | 3.11170 | –1.42098 |
| H | –1.19316 | 4.39281 | –0.42968 |
| H | –4.04671 | –0.67105 | 1.46697 |
| H | 2.72153 | 4.11642 | –1.55081 |
| H | 2.98180 | 4.75663 | 0.08147 |
| H | 0.85115 | 3.82253 | 0.05415 |
| H | 2.76006 | 1.97159 | 1.51869 |
| H | 0.55884 | 3.24081 | 2.59435 |
| H | 0.87623 | 0.46722 | 1.94582 |
| H | –0.94418 | 2.34632 | 0.78995 |
| H | –0.06785 | 0.47911 | –0.61662 |
| H | 3.62688 | 0.98284 | –0.93700 |
| H | –0.34037 | –5.23261 | 0.27191 |
| H | 4.44883 | 3.07710 | 0.40612 |
| H | –1.26881 | –3.82354 | 0.46952 |
| H | 1.96879 | –5.13304 | 0.12059 |
| H | –1.83630 | –0.17666 | 1.32104 |
| H | –1.59757 | –1.50119 | 0.36696 |
| H | –5.37963 | –3.49660 | –0.06701 |

**Table S26.** Optimized geometry coordinates of 2G:SAM (S-edge)
Energy: –1711141.3262374 (au)

| | | | |
|---|---|---|---|
| O | –9.18430 | –1.02445 | 0.29999 |
| C | –8.52825 | –2.27942 | 0.22250 |
| C | –7.01521 | –2.15282 | 0.36415 |

| | | | |
|---|---|---|---|
| O | –6.52425 | –1.22735 | –0.63030 |
| C | –6.52291 | –1.58838 | 1.71071 |
| O | –6.44059 | –2.53684 | 2.74302 |
| C | –5.17279 | –0.97250 | 1.30599 |
| O | –4.14157 | –1.97518 | 1.32229 |
| C | –5.44515 | –0.45418 | –0.12226 |
| N | –5.77039 | 0.96407 | –0.17680 |
| C | –6.99329 | 1.58381 | 0.08063 |
| N | –6.95409 | 2.87765 | –0.09400 |
| C | –5.65589 | 3.14294 | –0.47746 |
| C | –5.02954 | 4.39126 | –0.83054 |
| O | –5.46238 | 5.52564 | –0.87814 |
| N | –3.65122 | 4.15204 | –1.18557 |
| C | –2.99070 | 2.95349 | –1.17367 |
| N | –1.68535 | 2.95044 | –1.56290 |
| N | –3.58095 | 1.81943 | –0.83204 |
| C | –4.90234 | 1.97381 | –0.52903 |
| N | 7.71895 | 4.30818 | –0.73638 |
| C | 6.68227 | 3.66368 | 0.06718 |
| C | 7.12197 | 3.70342 | 1.55386 |
| O | 7.99096 | 4.66214 | 1.83526 |
| O | 6.69037 | 2.93063 | 2.38797 |
| C | 6.30116 | 2.22125 | –0.35073 |
| C | 4.98777 | 1.76128 | 0.30931 |
| S | 4.85499 | 0.01165 | 0.86355 |
| C | 6.30856 | –0.21919 | 1.94061 |
| C | 5.26434 | –0.95225 | –0.64727 |
| C | 5.38990 | –2.44356 | –0.30183 |
| O | 4.34229 | –2.84563 | 0.59006 |
| C | 5.25278 | –3.35057 | –1.53347 |
| O | 5.92228 | –4.59106 | –1.36240 |
| C | 3.72362 | –3.56579 | –1.62237 |
| O | 3.36504 | –4.75038 | –2.27129 |
| C | 3.29049 | –3.53032 | –0.12713 |
| N | 2.00668 | –2.92526 | 0.12188 |
| C | 0.96188 | –3.53193 | 0.81530 |
| N | –0.13151 | –2.82224 | 0.84835 |
| C | 0.18930 | –1.67451 | 0.14830 |
| C | –0.58841 | –0.53495 | –0.18525 |
| N | –1.85601 | –0.38491 | 0.18893 |
| N | 0.01333 | 0.43719 | –0.92747 |
| C | 1.28597 | 0.27555 | –1.29405 |
| N | 2.11956 | –0.74234 | –1.03093 |
| C | 1.50411 | –1.70602 | –0.31177 |
| H | –8.89539 | –2.89277 | 1.05176 |
| H | –8.77060 | –2.79884 | –0.71402 |

| | | | |
|---|---|---|---|
| H | –6.56858 | –3.14918 | 0.20382 |
| H | –7.21013 | –0.79935 | 2.03313 |
| H | –5.55395 | –2.92736 | 2.69613 |
| H | –4.82442 | –0.18745 | 1.97772 |
| H | –4.31937 | –2.62358 | 0.62425 |
| H | –4.55993 | –0.58507 | –0.75453 |
| H | –7.85628 | 1.00913 | 0.38361 |
| H | –1.17362 | 3.81736 | –1.50973 |
| H | 8.22266 | 5.08125 | 0.97274 |
| H | 7.35699 | 4.72183 | –1.58914 |
| H | 5.79039 | 4.30131 | 0.00298 |
| H | 7.13572 | 1.56344 | –0.08473 |
| H | 6.19887 | 2.18534 | –1.43938 |
| H | 4.80507 | 2.30334 | 1.23862 |
| H | 4.12011 | 1.88372 | –0.34516 |
| H | 7.22908 | –0.26872 | 1.35798 |
| H | 6.14367 | –1.14943 | 2.48545 |
| H | 6.34282 | 0.63489 | 2.61966 |
| H | 6.18407 | –0.55995 | –1.08937 |
| H | 4.39374 | –0.76566 | –1.28523 |
| H | 6.34480 | –2.63203 | 0.20114 |
| H | 5.66446 | –2.91198 | –2.44573 |
| H | 5.79591 | –4.90552 | –0.45470 |
| H | 3.25201 | –2.73117 | –2.14899 |
| H | 4.12623 | –5.35108 | –2.20138 |
| H | 3.22144 | –4.55282 | 0.25711 |
| H | 1.09074 | –4.51047 | 1.25932 |
| H | –2.41342 | 0.40720 | –0.15801 |
| H | 1.70208 | 1.08304 | –1.89483 |
| H | –9.06567 | –0.58926 | –0.55483 |
| H | –3.17545 | 4.98894 | –1.50053 |
| H | –1.14460 | 2.09747 | –1.40473 |
| H | –2.32561 | –1.10951 | 0.72367 |
| H | 8.47252 | 3.67178 | –0.98451 |

**Table S27.** Optimized geometry coordinates of 1G:SAM (S-edge)
Energy: –1711146.3317487 (au)

| | | | |
|---|---|---|---|
| O | 5.96113 | 2.17106 | 1.57895 |
| C | 5.31024 | 3.40157 | 1.32326 |
| C | 4.07383 | 3.24200 | 0.44055 |
| O | 4.47482 | 2.85750 | –0.89859 |
| C | 3.10993 | 2.13688 | 0.91001 |
| O | 2.01191 | 2.60712 | 1.70327 |
| C | 2.59781 | 1.52770 | –0.40522 |
| O | 1.53325 | 2.35643 | –0.85974 |
| C | 3.82566 | 1.66980 | –1.31016 |

| | | | |
|---|---|---|---|
| N | 4.76728 | 0.53720 | –1.18244 |
| C | 6.11564 | 0.64908 | –1.51382 |
| N | 6.80078 | –0.44064 | –1.28893 |
| C | 5.87566 | –1.33089 | –0.79130 |
| C | 6.06667 | –2.68295 | –0.32779 |
| O | 7.05614 | –3.38570 | –0.29260 |
| N | 4.81802 | –3.20086 | 0.17713 |
| C | 3.61632 | –2.55212 | 0.19830 |
| N | 2.52778 | –3.23582 | 0.68607 |
| N | 3.45090 | –1.33349 | –0.27851 |
| C | 4.60710 | –0.75635 | –0.72142 |
| N | –1.08963 | 1.65467 | 1.91292 |
| C | –1.07026 | 0.34670 | 2.56711 |
| C | 0.14386 | –0.54273 | 2.21009 |
| O | 1.33271 | 0.04532 | 2.18148 |
| O | 0.03202 | –1.73535 | 1.98681 |
| C | –2.37864 | –0.43186 | 2.29644 |
| C | –3.56692 | 0.41999 | 2.72886 |
| S | –5.25630 | –0.31806 | 2.59853 |
| C | –5.09942 | –1.94884 | 3.39801 |
| C | –5.53491 | –0.79151 | 0.83570 |
| C | –5.82378 | 0.41191 | –0.05494 |
| O | –4.67239 | 1.27299 | –0.09897 |
| C | –6.14000 | –0.03285 | –1.51585 |
| O | –7.51271 | –0.04716 | –1.78903 |
| C | –5.34587 | 0.99844 | –2.35905 |
| O | –6.18128 | 2.09070 | –2.72277 |
| C | –4.20180 | 1.46209 | –1.43618 |
| N | –2.90564 | 0.82776 | –1.62337 |
| C | –1.70867 | 1.51443 | –1.62646 |
| N | –0.65419 | 0.73531 | –1.63547 |
| C | –1.16831 | –0.55680 | –1.63550 |
| C | –0.57282 | –1.84892 | –1.62815 |
| N | 0.74433 | –2.08560 | –1.63034 |
| N | –1.39957 | –2.92119 | –1.63239 |
| C | –2.71471 | –2.73404 | –1.60828 |
| N | –3.39419 | –1.57305 | –1.57688 |
| C | –2.56568 | –0.52158 | –1.61590 |
| H | 5.98470 | 4.13919 | 0.86776 |
| H | 4.99157 | 3.79777 | 2.29397 |
| H | 3.54486 | 4.20219 | 0.37202 |
| H | 3.65229 | 1.39833 | 1.50207 |
| H | 1.42595 | 3.01015 | 1.03401 |
| H | 2.27946 | 0.49572 | –0.27699 |
| H | 0.87962 | 1.77494 | –1.30287 |
| H | 3.54994 | 1.78578 | –2.36229 |

| | | | |
|---|---|---|---|
| H | 6.50820 | 1.57052 | –1.91961 |
| H | 1.77815 | –2.65312 | 1.05548 |
| H | –0.99783 | 0.52725 | 3.64868 |
| H | –2.44030 | –0.66563 | 1.22996 |
| H | –2.32631 | –1.38106 | 2.83433 |
| H | –3.63128 | 1.33506 | 2.13754 |
| H | –3.51837 | 0.69711 | 3.78797 |
| H | –4.37416 | –2.57211 | 2.87454 |
| H | –4.79245 | –1.77821 | 4.43138 |
| H | –6.08861 | –2.40944 | 3.38516 |
| H | –4.67724 | –1.35749 | 0.46294 |
| H | –6.41819 | –1.43745 | 0.87737 |
| H | –6.67557 | 0.98145 | 0.33634 |
| H | –5.72461 | –1.02675 | –1.69431 |
| H | –7.70939 | 0.76019 | –2.29441 |
| H | –4.98018 | 0.57457 | –3.29476 |
| H | –6.36252 | 2.64071 | –1.94578 |
| H | –4.02446 | 2.53032 | –1.58952 |
| H | –1.68213 | 2.59640 | –1.63059 |
| H | 1.44161 | –1.38360 | –1.41250 |
| H | –3.32077 | –3.63736 | –1.60626 |
| H | 6.44619 | 1.90884 | 0.78402 |
| H | 4.89917 | –4.14356 | 0.53997 |
| H | 2.72160 | –4.04378 | 1.26123 |
| H | 1.02985 | –3.04445 | –1.48168 |
| H | –0.53751 | 2.34087 | 2.41630 |
| H | –0.74351 | 1.60847 | 0.95791 |
| H | 1.34149 | 1.03890 | 2.18941 |

**Table S28.** Optimized geometry coordinates of G:SFG (S-edge)
Energy: –1495781.5260297 (au)

| | | | |
|---|---|---|---|
| O | 6.42728 | 1.67503 | –1.96450 |
| C | 5.55418 | 2.20939 | –2.95143 |
| C | 4.12464 | 2.12817 | –2.44733 |
| O | 3.71251 | 0.75716 | –2.37276 |
| C | 3.88758 | 2.69830 | –1.04315 |
| O | 3.62126 | 4.09656 | –1.13623 |
| C | 2.65509 | 1.89886 | –0.57709 |
| O | 1.48047 | 2.47265 | –1.12844 |
| C | 2.88729 | 0.53153 | –1.24242 |
| N | 3.55817 | –0.41664 | –0.34660 |
| C | 4.86690 | –0.88620 | –0.38629 |
| N | 5.11787 | –1.77747 | 0.54045 |
| C | 3.92735 | –1.91029 | 1.23163 |
| C | 3.57778 | –2.73647 | 2.35784 |
| O | 4.22792 | –3.55407 | 2.98892 |

| | | | |
|---|---|---|---|
| N | 2.20828 | –2.49774 | 2.72587 |
| C | 1.32482 | –1.62827 | 2.12985 |
| N | 0.05072 | –1.58878 | 2.60358 |
| N | 1.66597 | –0.88000 | 1.09318 |
| C | 2.94897 | –1.07223 | 0.69617 |
| N | –0.94604 | 0.76494 | 1.10776 |
| C | –0.76219 | 2.12162 | 1.62956 |
| C | 0.13855 | 2.05655 | 2.86010 |
| O | 0.01075 | 1.30173 | 3.79830 |
| O | 1.11717 | 3.00044 | 2.82301 |
| C | –2.12544 | 2.70773 | 2.05580 |
| C | –2.97755 | 3.18515 | 0.85661 |
| C | –4.49496 | 2.95668 | 1.00999 |
| N | –4.95770 | 3.60197 | 2.24779 |
| C | –4.91488 | 1.46993 | 0.88360 |
| C | –4.83652 | 0.86880 | –0.51215 |
| O | –3.46815 | 0.65426 | –0.91899 |
| C | –5.45604 | –0.53458 | –0.65497 |
| O | –6.85947 | –0.55083 | –0.69538 |
| C | –4.79789 | –1.05272 | –1.95374 |
| O | –5.62917 | –0.65219 | –3.05074 |
| C | –3.45469 | –0.27524 | –2.00596 |
| N | –2.25757 | –1.07975 | –1.92117 |
| C | –1.03367 | –0.74139 | –2.48108 |
| N | –0.05472 | –1.54013 | –2.14464 |
| C | –0.64937 | –2.46159 | –1.29615 |
| C | –0.14565 | –3.57913 | –0.59397 |
| N | 1.14575 | –3.97302 | –0.67973 |
| N | –0.99274 | –4.29685 | 0.16859 |
| C | –2.27500 | –3.91686 | 0.23165 |
| N | –2.87485 | –2.88522 | –0.37620 |
| C | –2.00936 | –2.19202 | –1.12724 |
| H | 5.62207 | 1.65214 | –3.89597 |
| H | 5.77902 | 3.26695 | –3.15918 |
| H | 3.47926 | 2.66938 | –3.15441 |
| H | 4.75427 | 2.49447 | –0.40711 |
| H | 3.82419 | 4.50139 | –0.28291 |
| H | 2.57736 | 1.81349 | 0.51315 |
| H | 1.93891 | 0.08323 | –1.55368 |
| H | 5.56412 | –0.50301 | –1.11561 |
| H | –0.49836 | –0.78347 | 2.27027 |
| H | –0.11548 | 0.47103 | 0.59325 |
| H | –0.28491 | 2.79401 | 0.90390 |
| H | –2.62576 | 1.91813 | 2.62922 |
| H | –1.97226 | 3.54327 | 2.74653 |
| H | –2.65030 | 2.69020 | –0.06174 |

| | | | |
|---|---|---|---|
| H | –2.81252 | 4.25846 | 0.71622 |
| H | –4.98198 | 3.49664 | 0.18559 |
| H | –5.97188 | 3.53970 | 2.30995 |
| H | –4.59986 | 3.09547 | 3.05571 |
| H | –5.96867 | 1.38517 | 1.18032 |
| H | –4.34173 | 0.83957 | 1.57621 |
| H | –5.32850 | 1.54221 | –1.23442 |
| H | –5.12285 | –1.15975 | 0.17944 |
| H | –7.08754 | –0.37981 | –1.62452 |
| H | –4.65007 | –2.13253 | –1.93397 |
| H | –5.72774 | –1.39808 | –3.65308 |
| H | –3.41007 | 0.26498 | –2.95977 |
| H | 7.31213 | 1.60222 | –2.34204 |
| H | –2.91391 | –4.53072 | 0.86285 |
| H | –0.94372 | 0.11876 | –3.13174 |
| H | 1.82634 | –3.34342 | –1.07538 |
| H | 1.46408 | –4.67483 | –0.02911 |
| H | –0.08824 | –1.82406 | 3.57546 |
| H | 1.89419 | –3.08104 | 3.49179 |
| H | 1.60072 | 2.90907 | 3.66251 |
| H | –1.73709 | 0.73742 | 0.46718 |
| H | 1.67049 | 3.42366 | –1.20140 |

**Table S29.** Optimized geometry coordinates of G:TPP (S-edge)
Energy: –2092112.5739857 (au)

| | | | |
|---|---|---|---|
| O | 8.13696 | 3.97087 | –1.60121 |
| C | 7.22776 | 4.58170 | –0.69543 |
| C | 6.63441 | 3.60795 | 0.31759 |
| O | 5.74881 | 2.67553 | –0.34825 |
| C | 7.64561 | 2.73372 | 1.08723 |
| O | 8.21341 | 3.36446 | 2.20646 |
| C | 6.78882 | 1.50094 | 1.42637 |
| O | 6.02685 | 1.74666 | 2.60912 |
| C | 5.89126 | 1.36692 | 0.17927 |
| N | 6.43072 | 0.47194 | –0.84015 |
| C | 7.40788 | 0.75928 | –1.79584 |
| N | 7.80804 | –0.30240 | –2.44488 |
| C | 7.08292 | –1.34371 | –1.90372 |
| C | 7.13677 | –2.75581 | –2.18535 |
| O | 7.80459 | –3.39500 | –2.97352 |
| N | 6.21737 | –3.45474 | –1.31805 |
| C | 5.36542 | –2.90026 | –0.40351 |
| N | 4.56543 | –3.73840 | 0.31008 |
| N | 5.31398 | –1.59609 | –0.17495 |
| C | 6.21614 | –0.88640 | –0.91587 |
| N | –0.04645 | –2.69044 | 2.61594 |

| | | | |
|---|---|---|---|
| C | 1.22991 | –3.04470 | 2.39090 |
| C | 1.73401 | –4.26613 | 3.10526 |
| N | 2.07792 | –2.41388 | 1.57002 |
| C | 1.64662 | –1.31706 | 0.91119 |
| N | 2.54479 | –0.68380 | 0.12933 |
| C | 0.29510 | –0.88652 | 1.05596 |
| C | –0.48202 | –1.62580 | 1.93772 |
| C | –0.20352 | 0.32463 | 0.30333 |
| N | –1.68353 | 0.40714 | 0.24783 |
| C | –2.37795 | 1.17576 | 1.07936 |
| S | –4.04939 | 1.06504 | 0.84469 |
| C | –3.81074 | –0.10199 | –0.44418 |
| C | –2.47420 | –0.34328 | –0.63297 |
| C | –1.82470 | –1.25731 | –1.62566 |
| C | –4.98148 | –0.73492 | –1.14797 |
| C | –6.04213 | 0.25937 | –1.63094 |
| O | –6.51649 | 1.03225 | –0.50904 |
| P | –7.85799 | 0.67904 | 0.30758 |
| O | –7.84211 | –0.91818 | 0.34215 |
| O | –7.97172 | 1.43644 | 1.56050 |
| O | –9.02249 | 1.03107 | –0.81181 |
| P | –10.52368 | 0.41761 | –0.68801 |
| O | –11.48241 | 1.57471 | –0.15163 |
| O | –10.51593 | –0.83103 | 0.12108 |
| O | –10.87666 | 0.35608 | –2.23607 |
| H | 6.41334 | 5.00478 | –1.29256 |
| H | 7.69861 | 5.40238 | –0.13945 |
| H | 6.05083 | 4.20283 | 1.04012 |
| H | 8.45622 | 2.44287 | 0.41109 |
| H | 7.64511 | 3.14042 | 2.96152 |
| H | 7.36741 | 0.59836 | 1.62591 |
| H | 5.37662 | 2.43855 | 2.42007 |
| H | 4.91504 | 0.96490 | 0.47209 |
| H | 7.73643 | 1.77633 | –1.96296 |
| H | 4.37891 | –4.65876 | –0.05671 |
| H | 2.82152 | –4.25073 | 3.19156 |
| H | 1.44246 | –5.16688 | 2.55177 |
| H | 1.27885 | –4.33456 | 4.09486 |
| H | 2.31877 | 0.16787 | –0.35584 |
| H | –1.52355 | –1.35669 | 2.11244 |
| H | 0.14021 | 1.26643 | 0.74659 |
| H | –1.90732 | 1.81452 | 1.81432 |
| H | –1.13906 | –1.95074 | –1.13079 |
| H | –1.26118 | –0.69388 | –2.37763 |
| H | –2.57538 | –1.84707 | –2.15012 |
| H | –4.61638 | –1.29708 | –2.01211 |

| | | | |
|---|---|---|---|
| H | –5.46589 | –1.45344 | –0.47693 |
| H | –6.86340 | –0.27697 | –2.11098 |
| H | –5.62937 | 0.98184 | –2.33877 |
| H | 0.14750 | 0.31754 | –0.73371 |
| H | 3.52710 | –1.00779 | 0.09382 |
| H | 6.26543 | –4.46154 | –1.42043 |
| H | 3.82350 | –3.30987 | 0.86467 |
| H | 9.03543 | 4.08404 | –1.26754 |
| H | –8.77043 | –1.25834 | 0.42167 |
| H | –11.60464 | 1.54921 | 0.81069 |
| H | –11.82310 | 0.25653 | –2.41970 |

**Table S30.** Optimized geometry coordinates of G:TPS (S–edge)

Energy: –1735853.8235357 (au)

| | | | |
|---|---|---|---|
| O | 5.34380 | 4.94130 | –1.05438 |
| C | 4.02785 | 5.27866 | –0.64591 |
| C | 3.45503 | 4.14398 | 0.18382 |
| O | 3.22958 | 2.97234 | –0.63767 |
| C | 4.33660 | 3.66367 | 1.35842 |
| O | 4.19407 | 4.43352 | 2.52548 |
| C | 3.86369 | 2.20708 | 1.51199 |
| O | 2.67650 | 2.15911 | 2.30885 |
| C | 3.58671 | 1.78757 | 0.05494 |
| N | 4.72789 | 1.14951 | –0.59817 |
| C | 5.77740 | 1.76712 | –1.28085 |
| N | 6.72304 | 0.92998 | –1.61851 |
| C | 6.30293 | –0.29261 | –1.13806 |
| C | 6.96299 | –1.57309 | –1.16142 |
| O | 8.02827 | –1.92199 | –1.62928 |
| N | 6.15245 | –2.54493 | –0.46444 |
| C | 4.92108 | –2.34111 | 0.09260 |
| N | 4.32198 | –3.39396 | 0.71270 |
| N | 4.31353 | –1.16368 | 0.07335 |
| C | 5.06644 | –0.18220 | –0.50905 |
| C | 1.21920 | –4.83077 | 2.84994 |
| N | –2.83331 | –1.80191 | –1.15437 |
| C | –3.57591 | –2.59146 | –1.92200 |
| S | –5.22676 | –2.46829 | –1.57176 |
| C | –4.90698 | –1.28892 | –0.31989 |
| C | –3.56113 | –1.04225 | –0.22950 |
| C | –2.86171 | –0.09203 | 0.69076 |
| C | –6.03478 | –0.66943 | 0.45250 |
| C | –6.67241 | 0.52700 | –0.27181 |
| O | –7.66309 | 1.03893 | 0.61784 |
| N | –0.74826 | –4.32207 | 1.52107 |
| C | 0.54031 | –4.03382 | 1.77347 |

| | | | |
|---|---|---|---|
| N | 1.27075 | –3.11718 | 1.12689 |
| C | 0.68564 | –2.39055 | 0.15272 |
| N | 1.44488 | –1.44910 | –0.45395 |
| C | –0.68422 | –2.61059 | –0.17188 |
| C | –1.32431 | –3.60925 | 0.55119 |
| C | –1.35061 | –1.78716 | –1.24747 |
| P | –8.64749 | 2.24622 | 0.18942 |
| O | –7.55925 | 3.31098 | –0.33648 |
| O | –9.58328 | 2.64803 | 1.24883 |
| O | –9.33281 | 1.73905 | –1.18639 |
| H | 3.37011 | 5.45873 | –1.50875 |
| H | 4.01841 | 6.17745 | –0.01191 |
| H | 2.48799 | 4.49365 | 0.58130 |
| H | 5.38602 | 3.69583 | 1.05310 |
| H | 3.49361 | 4.00887 | 3.04684 |
| H | 4.58221 | 1.55427 | 2.00828 |
| H | 1.96532 | 2.60220 | 1.82407 |
| H | 2.75752 | 1.07146 | 0.02891 |
| H | 5.75168 | 2.82713 | –1.49029 |
| H | 4.61043 | –4.33031 | 0.47602 |
| H | 0.52268 | –5.02722 | 3.66719 |
| H | 1.52830 | –5.80400 | 2.45028 |
| H | 2.10297 | –4.31223 | 3.22395 |
| H | –3.15625 | –3.24369 | –2.67551 |
| H | –2.07607 | –0.59763 | 1.25928 |
| H | –2.40771 | 0.73564 | 0.13482 |
| H | –3.57122 | 0.33734 | 1.39786 |
| H | –6.81025 | –1.41370 | 0.65865 |
| H | –5.66533 | –0.33213 | 1.42488 |
| H | –7.13405 | 0.21501 | –1.21660 |
| H | –5.92450 | 1.29800 | –0.48679 |
| H | 1.18792 | –1.08459 | –1.35707 |
| H | –2.37062 | –3.84671 | 0.35820 |
| H | –1.10644 | –2.12873 | –2.26010 |
| H | –1.04084 | –0.73921 | –1.18236 |
| H | 2.44120 | –1.34994 | –0.18623 |
| H | –7.94576 | 4.18483 | –0.49534 |
| H | –10.27377 | 1.55674 | –1.04247 |
| H | 5.73405 | 5.69361 | –1.51581 |
| H | 3.35380 | –3.26999 | 1.00686 |
| H | 6.60493 | –3.44730 | –0.37769 |

**Table S31.** Optimized geometry coordinates of U:29G (W–edge)
Energy: –612236.6908763 (au)

| | | | |
|---|---|---|---|
| N | –4.63688 | 0.83507 | –0.01131 |
| C | –3.26128 | 1.06500 | –0.00506 |

| | | | |
|---|---|---|---|
| O | –2.79497 | 2.22211 | –0.00668 |
| N | –2.49671 | –0.08230 | 0.00291 |
| C | –2.97203 | –1.39932 | 0.00541 |
| O | –2.18188 | –2.35914 | 0.01263 |
| C | –4.42378 | –1.53103 | –0.00105 |
| C | –5.19365 | –0.42118 | –0.00919 |
| N | 0.68978 | –2.00324 | 0.00202 |
| C | 1.23276 | –0.77254 | 0.00315 |
| N | 0.41416 | 0.28202 | 0.00749 |
| C | 2.66718 | –0.58403 | –0.00068 |
| C | 3.63868 | –1.59345 | –0.00454 |
| N | 4.94188 | –1.34731 | –0.00831 |
| C | 5.27144 | –0.03072 | –0.00823 |
| N | 4.47297 | 1.01731 | –0.00459 |
| C | 3.13268 | 0.76587 | –0.00044 |
| N | 2.29659 | 1.81732 | 0.00380 |
| C | 0.99283 | 1.52952 | 0.00795 |
| N | 0.14182 | 2.57466 | 0.01367 |
| H | –4.84807 | –2.52487 | 0.00062 |
| H | –6.27735 | –0.45322 | –0.01449 |
| H | 1.27421 | –2.82087 | –0.00015 |
| H | 3.35808 | –2.64803 | –0.00452 |
| H | 0.54968 | 3.49502 | 0.01051 |
| H | –1.45839 | 0.05415 | 0.00611 |
| H | –5.20903 | 1.66708 | –0.01691 |
| H | –0.32399 | –2.12888 | 0.00701 |
| H | –0.86518 | 2.45466 | 0.01222 |
| H | 6.34061 | 0.17767 | –0.01158 |

**Table S32.** Optimized geometry coordinates of U:2BP (W–edge)
Energy: –553581.1112816 (au)

| | | | |
|---|---|---|---|
| N | –3.95586 | 0.74644 | 0.02831 |
| C | –2.59939 | 1.05916 | 0.02105 |
| O | –2.21048 | 2.22237 | 0.03110 |
| N | –1.76934 | –0.03588 | 0.00538 |
| C | –2.15213 | –1.37821 | –0.00681 |
| O | –1.30990 | –2.27794 | –0.01382 |
| C | –3.59162 | –1.60243 | –0.00855 |
| C | –4.42956 | –0.54307 | 0.01103 |
| N | 1.57279 | –1.81221 | –0.03144 |
| C | 2.80335 | –2.44997 | 0.00679 |
| N | 3.82730 | –1.63528 | 0.03602 |
| C | 3.25187 | –0.37107 | 0.01849 |
| C | 3.75221 | 0.93216 | 0.04461 |
| N | 2.93493 | 1.98490 | 0.02217 |
| C | 1.59493 | 1.75824 | –0.02632 |

| | | | |
|---|---|---|---|
| N | 0.79277 | 2.84816 | –0.06346 |
| N | 0.99120 | 0.54809 | –0.04693 |
| C | 1.84753 | –0.47443 | –0.02397 |
| H | –3.95497 | –2.62013 | –0.02195 |
| H | –5.50884 | –0.64564 | 0.01381 |
| H | 0.63056 | –2.20806 | –0.04646 |
| H | 2.87026 | –3.52970 | 0.01007 |
| H | 4.82080 | 1.13471 | 0.08528 |
| H | 1.22497 | 3.75382 | –0.00393 |
| H | –0.21681 | 2.74707 | –0.07174 |
| H | –0.74138 | 0.18166 | –0.01443 |
| H | –4.58134 | 1.53899 | 0.04288 |

**Table S33.** Optimized geometry coordinates of 1U:3AY (W–edge)
Energy: –530438.8046776 (au)

| | | | |
|---|---|---|---|
| N | –3.79810 | –1.17966 | –0.12333 |
| C | –2.40580 | –1.21077 | –0.14143 |
| O | –1.80412 | –2.27190 | –0.26992 |
| N | –1.80613 | 0.01817 | –0.00630 |
| C | –2.46192 | 1.24523 | 0.14103 |
| O | –1.82656 | 2.29356 | 0.24502 |
| C | –3.91800 | 1.17216 | 0.15737 |
| C | –4.52651 | –0.02495 | 0.02414 |
| N | 1.06422 | 2.33644 | –0.29439 |
| C | 1.78274 | 1.19237 | –0.12813 |
| C | 3.18534 | 1.20472 | –0.10984 |
| N | 1.07947 | 0.03992 | –0.01462 |
| C | 1.80685 | –1.09371 | 0.11180 |
| N | 1.12040 | –2.25638 | 0.26279 |
| N | 3.14615 | –1.19032 | 0.12468 |
| C | 3.82039 | –0.03343 | 0.02059 |
| N | 5.19159 | –0.13729 | 0.08906 |
| H | –4.47464 | 2.09135 | 0.27211 |
| H | –5.60398 | –0.14621 | 0.02442 |
| H | 0.06781 | 2.33224 | –0.08161 |
| H | 3.74263 | 2.13133 | –0.18910 |
| H | 1.66840 | –3.09913 | 0.21091 |
| H | 5.54454 | –1.07227 | –0.05581 |
| H | –4.24491 | –2.07993 | –0.21889 |
| H | –0.75087 | 0.02703 | –0.01391 |
| H | 0.12996 | –2.29972 | 0.04364 |
| H | 5.73711 | 0.60299 | –0.32475 |
| H | 1.54772 | 3.21306 | –0.18614 |

**Table S34.** Optimized geometry coordinates of 2U:3AY (W–edge)
Energy: –530438.6992134 (au)

| | | | |
|---|---|---|---|
| N | 3.80496 | –1.18561 | 0.05799 |
| C | 2.41316 | –1.21547 | 0.06830 |
| O | 1.81044 | –2.28158 | 0.13595 |
| N | 1.81337 | 0.01941 | –0.00324 |
| C | 2.47180 | 1.25101 | –0.09029 |
| O | 1.84095 | 2.30533 | –0.15110 |
| C | 3.92797 | 1.17663 | –0.10126 |
| C | 4.53538 | –0.02594 | –0.02630 |
| N | –5.20214 | –0.13787 | –0.01023 |
| C | –3.82731 | –0.03072 | –0.01953 |
| C | –3.19056 | 1.20987 | 0.06500 |
| N | –3.15667 | –1.19209 | –0.07454 |
| C | –1.81647 | –1.10022 | –0.03545 |
| N | –1.13746 | –2.27063 | –0.09807 |
| N | –1.08516 | 0.03643 | 0.04124 |
| C | –1.78743 | 1.19438 | 0.09723 |
| N | –1.07023 | 2.34198 | 0.21148 |
| H | 4.48524 | 2.10022 | –0.16703 |
| H | 5.61271 | –0.14797 | –0.02732 |
| H | –5.54627 | –1.03589 | –0.31965 |
| H | –3.74612 | 2.13868 | 0.12893 |
| H | –0.12926 | –2.30972 | 0.00987 |
| H | –0.06123 | 2.32657 | 0.07884 |
| H | 4.24989 | –2.09043 | 0.10948 |
| H | 0.75882 | 0.02493 | 0.01281 |
| H | –1.68399 | –3.11474 | –0.08887 |
| H | –5.73023 | 0.65933 | –0.33126 |
| H | –1.54661 | 3.22046 | 0.09670 |

**Table S35.** Optimized geometry coordinates of U:6AP (W–edge)
Energy: –588326.5898595 (au)

| | | | |
|---|---|---|---|
| N | –4.57532 | 0.74204 | –0.00155 |
| C | –3.22443 | 1.09995 | 0.00154 |
| O | –2.86869 | 2.26916 | 0.00417 |
| N | –2.36591 | 0.02334 | 0.00094 |
| C | –2.70941 | –1.32727 | –0.00159 |
| O | –1.84024 | –2.20048 | –0.00096 |
| C | –4.14274 | –1.59497 | –0.00494 |
| C | –5.01099 | –0.56009 | –0.00486 |
| N | 1.92882 | 2.37651 | 0.00278 |
| C | 0.57789 | 2.10871 | 0.00433 |
| N | 0.32874 | 0.82111 | 0.00271 |
| C | 1.57187 | 0.20662 | –0.00213 |
| C | 2.58940 | 1.16340 | –0.00196 |
| N | 3.90673 | 0.94705 | –0.00314 |
| C | 4.17761 | –0.37253 | –0.01012 |

| | | | |
|---|---|---|---|
| N | 5.50565 | –0.71872 | –0.06578 |
| N | 3.30951 | –1.40729 | 0.00023 |
| C | 1.99052 | –1.14818 | 0.00300 |
| N | 1.12935 | –2.18223 | 0.01625 |
| H | –4.47537 | –2.62326 | –0.00749 |
| H | –6.08720 | –0.69374 | –0.00732 |
| H | 2.36881 | 3.28378 | –0.00072 |
| H | –0.18130 | 2.87852 | 0.00649 |
| H | 5.72348 | –1.66532 | 0.20307 |
| H | 6.15997 | 0.00090 | 0.19910 |
| H | 1.52182 | –3.10993 | 0.00523 |
| H | 0.11780 | –2.06664 | 0.00856 |
| H | –5.22301 | 1.51629 | –0.00181 |
| H | –1.34469 | 0.26344 | 0.00229 |

**Table S36.** Optimized geometry coordinates of U:6GO (W–edge)
Energy: –625444.0533693 (au)

| | | | |
|---|---|---|---|
| N | –4.42352 | –1.20454 | –0.35936 |
| C | –3.03262 | –1.18325 | –0.34954 |
| O | –2.39253 | –2.17862 | –0.68709 |
| N | –2.48408 | 0.00400 | 0.05825 |
| C | –3.18397 | 1.14915 | 0.50336 |
| O | –2.58720 | 2.13848 | 0.89012 |
| C | –4.64156 | 1.01552 | 0.45132 |
| C | –5.19972 | –0.13592 | 0.02907 |
| C | 5.13599 | –1.72262 | 0.44408 |
| O | 4.58118 | –0.42066 | 0.20646 |
| C | 3.24120 | –0.34251 | 0.09287 |
| N | 2.48173 | –1.41693 | 0.18850 |
| C | 2.65738 | 0.90701 | –0.13693 |
| N | 3.09737 | 2.20783 | –0.31106 |
| C | 1.96947 | 2.95575 | –0.51711 |
| N | 0.85297 | 2.26066 | –0.49252 |
| C | 1.25895 | 0.96559 | –0.25107 |
| N | 0.46658 | –0.11683 | –0.13932 |
| C | 1.12750 | –1.26743 | 0.06883 |
| N | 0.39563 | –2.40371 | 0.20057 |
| H | –5.23886 | 1.86087 | 0.76302 |
| H | –6.27135 | –0.29198 | –0.02766 |
| H | 4.75399 | –2.14415 | 1.37714 |
| H | 4.89361 | –2.40445 | –0.37489 |
| H | 6.21352 | –1.56875 | 0.50691 |
| H | 2.02609 | 4.02358 | –0.68327 |
| H | 0.90858 | –3.26963 | 0.17845 |
| H | –0.57051 | –2.39423 | –0.12406 |
| H | –1.44684 | 0.05940 | 0.02381 |

| | | | |
|---|---|---|---|
| H | –4.83329 | –2.07777 | –0.65539 |
| H | 4.05088 | 2.53233 | –0.28473 |

**Table S37.** Optimized geometry coordinates of U:6GU (W–edge)
Energy: –841982.4906724 (au)

| | | | |
|---|---|---|---|
| N | –4.49058 | –1.03383 | –0.00016 |
| C | –3.11140 | –1.22144 | –0.00025 |
| O | –2.61527 | –2.34281 | –0.00057 |
| N | –2.38597 | –0.05396 | 0.00004 |
| C | –2.88896 | 1.24786 | 0.00018 |
| O | –2.13184 | 2.22106 | 0.00024 |
| C | –4.34249 | 1.33937 | 0.00016 |
| C | –5.07983 | 0.20720 | –0.00005 |
| Cl | 4.93466 | –0.63414 | –0.00023 |
| N | 2.49047 | –1.60909 | 0.00016 |
| C | 1.13309 | –1.52311 | 0.00031 |
| N | 0.44762 | –2.68748 | 0.00086 |
| N | 0.41734 | –0.37825 | 0.00010 |
| C | 1.17369 | 0.72095 | 0.00006 |
| C | 2.58339 | 0.76648 | 0.00007 |
| C | 3.18997 | –0.49305 | 0.00004 |
| N | 3.02578 | 2.07822 | –0.00004 |
| C | 1.92281 | 2.78357 | –0.00015 |
| N | 0.76494 | 2.02319 | –0.00017 |
| H | –4.79742 | 2.31967 | 0.00030 |
| H | –6.16402 | 0.20994 | –0.00009 |
| H | 0.96945 | –3.54685 | –0.00035 |
| H | 1.88033 | 3.86434 | –0.00027 |
| H | –0.21649 | 2.31616 | 0.00018 |
| H | –0.56727 | –2.68730 | –0.00012 |
| H | –1.34441 | –0.17962 | 0.00011 |
| H | –5.04108 | –1.88047 | –0.00076 |

**Table S38.** Optimized geometry coordinates of 1U:7DG (W–edge)
Energy: –590711.2942665 (au)

| | | | |
|---|---|---|---|
| N | –4.72347 | –0.85735 | –0.01401 |
| C | –3.42801 | –1.39155 | 0.01520 |
| O | –3.23161 | –2.58948 | 0.04043 |
| N | –2.43184 | –0.42573 | 0.01199 |
| C | –2.61745 | 0.95128 | –0.01759 |
| O | –1.65184 | 1.72938 | –0.01685 |
| C | –3.99892 | 1.39960 | –0.04794 |
| C | –4.99322 | 0.48347 | –0.04446 |
| N | 4.86336 | –0.33083 | –0.04823 |
| C | 4.75588 | –1.61624 | –0.04963 |
| C | 3.34134 | –2.14954 | –0.01659 |

| | | | |
|---|---|---|---|
| C | 2.59594 | –0.85688 | 0.00709 |
| C | 1.21190 | –0.54198 | 0.03265 |
| O | 0.23957 | –1.31700 | 0.04129 |
| N | 0.99604 | 0.84995 | 0.04666 |
| C | 1.98927 | 1.78405 | 0.02821 |
| N | 1.59947 | 3.08752 | 0.08096 |
| N | 3.27926 | 1.49264 | –0.00632 |
| C | 3.53018 | 0.16750 | –0.01278 |
| H | –4.19426 | 2.46199 | –0.07080 |
| H | –6.04327 | 0.75390 | –0.06492 |
| H | 5.63946 | –2.25059 | –0.07307 |
| H | 3.12667 | –2.77710 | –0.89286 |
| H | 0.64032 | 3.33170 | –0.11319 |
| H | 2.31708 | 3.76997 | –0.10508 |
| H | 3.17029 | –2.78425 | 0.86409 |
| H | 0.00866 | 1.15400 | 0.05676 |
| H | –1.45321 | –0.78825 | 0.02936 |
| H | –5.46188 | –1.54619 | –0.01131 |

**Table S39.** Optimized geometry coordinates of 2U:7DG (W–edge)
Energy: –590705.7215570 (au)

| | | | |
|---|---|---|---|
| N | 4.18522 | –0.96308 | 0.33502 |
| C | 2.80070 | –1.05106 | 0.29128 |
| O | 2.23667 | –2.11707 | 0.54835 |
| N | 2.16466 | 0.11073 | –0.05271 |
| C | 2.77772 | 1.33695 | –0.40976 |
| O | 2.10814 | 2.29458 | –0.75070 |
| C | 4.23929 | 1.31633 | –0.32742 |
| C | 4.88013 | 0.18821 | 0.03560 |
| N | –1.29801 | 1.99211 | 0.46050 |
| C | –2.38514 | 2.68624 | 0.48411 |
| C | –3.65465 | 1.90282 | 0.23893 |
| C | –3.07028 | 0.54014 | 0.06063 |
| C | –3.66022 | –0.73296 | –0.19607 |
| O | –4.83257 | –1.05073 | –0.34978 |
| N | –2.64441 | –1.74459 | –0.26364 |
| C | –1.29853 | –1.54027 | –0.12284 |
| N | –0.46961 | –2.60097 | –0.24621 |
| N | –0.78547 | –0.33593 | 0.10118 |
| C | –1.70135 | 0.65748 | 0.19325 |
| H | 4.77250 | 2.22493 | –0.56955 |
| H | 5.95925 | 0.11436 | 0.11052 |
| H | –2.35993 | 3.75727 | 0.67143 |
| H | –4.35013 | 1.98753 | 1.08557 |
| H | –0.84541 | –3.53455 | –0.20445 |
| H | –3.00436 | –2.67163 | –0.45529 |

| | | | |
|---|---|---|---|
| H | –4.19539 | 2.27737 | –0.64145 |
| H | 0.50393 | –2.46446 | 0.04600 |
| H | 1.12937 | 0.08185 | –0.03635 |
| H | 4.65947 | –1.81798 | 0.58526 |

**Table S40.** Optimized geometry coordinates of U:AMZ (W–edge)
Energy: –854276.0121636 (au)

| | | | |
|---|---|---|---|
| N | –5.33677 | –0.73941 | 0.27121 |
| C | –4.24560 | 0.11715 | 0.13949 |
| O | –4.37304 | 1.33401 | 0.19965 |
| N | –3.04699 | –0.52558 | –0.05789 |
| C | –2.83463 | –1.90368 | –0.11531 |
| O | –1.70041 | –2.35359 | –0.28356 |
| C | –4.03535 | –2.71410 | 0.03589 |
| C | –5.22851 | –2.10711 | 0.22007 |
| O | –0.16649 | 4.57769 | 0.35828 |
| C | –0.81862 | 3.53028 | 0.20023 |
| N | –2.16667 | 3.48837 | 0.33041 |
| C | –0.13195 | 2.27597 | –0.13092 |
| C | 1.24499 | 2.15928 | –0.26076 |
| N | 2.20533 | 3.14910 | –0.26772 |
| N | –0.72159 | 1.04138 | –0.35992 |
| C | 0.25395 | 0.20589 | –0.61125 |
| N | 1.48461 | 0.82592 | –0.54008 |
| C | 2.76575 | 0.19083 | –0.89102 |
| C | 3.79799 | 0.17225 | 0.24607 |
| C | 3.40063 | –1.08379 | 1.05418 |
| O | 4.47815 | –1.63042 | 1.77389 |
| O | 5.11170 | 0.00967 | –0.28443 |
| O | 2.55921 | –1.16468 | –1.19977 |
| C | 2.80406 | –2.01232 | –0.03534 |
| C | 1.54222 | –2.72959 | 0.43953 |
| O | 0.93574 | –3.43829 | –0.61679 |
| H | –3.94059 | –3.78996 | –0.00469 |
| H | –6.15672 | –2.65506 | 0.33762 |
| H | –2.73021 | 2.65541 | 0.19967 |
| H | 1.75525 | 4.02572 | 0.00465 |
| H | 0.14450 | –0.83537 | –0.86267 |
| H | 3.16441 | 0.70294 | –1.77456 |
| H | 3.82512 | 1.07384 | 0.86087 |
| H | 2.62382 | –0.82273 | 1.78233 |
| H | 5.27425 | –1.40434 | 1.26446 |
| H | 5.05404 | –0.59998 | –1.03615 |
| H | 3.54401 | –2.75632 | –0.34991 |
| H | 0.85022 | –2.00086 | 0.88068 |
| H | 1.85743 | –3.40424 | 1.25428 |

| | | | |
|---|---|---|---|
| H | –2.62809 | 4.35714 | 0.54963 |
| H | 3.05454 | 2.96104 | 0.25028 |
| H | –0.02150 | –3.29131 | –0.55881 |
| H | –6.22651 | –0.28403 | 0.41502 |
| H | –2.21790 | 0.10826 | –0.16799 |

**Table S41.** Optimized geometry coordinates of U:FFO (W–edge)
Energy: –1007601.9050733 (au)

| | | | |
|---|---|---|---|
| N | 7.20413 | –1.45958 | –0.88259 |
| C | 6.19783 | –0.50777 | –0.78929 |
| O | 6.35038 | 0.62630 | –1.23753 |
| N | 5.05569 | –0.94830 | –0.16830 |
| C | 4.83030 | –2.22800 | 0.35458 |
| O | 3.75698 | –2.50511 | 0.88629 |
| C | 5.94538 | –3.15332 | 0.20938 |
| C | 7.07850 | –2.73968 | –0.39680 |
| N | 2.90130 | 1.00665 | 0.05693 |
| C | 3.05309 | 2.22528 | –0.42880 |
| N | 4.26063 | 2.62784 | –0.89324 |
| N | 2.02844 | 3.12257 | –0.45378 |
| C | 0.71319 | 2.86023 | 0.02596 |
| O | –0.15124 | 3.72507 | –0.11378 |
| C | 0.60388 | 1.58123 | 0.64878 |
| N | –0.63939 | 1.12939 | 1.17026 |
| C | –0.92070 | –0.30041 | 1.01940 |
| C | 0.27787 | –1.07837 | 1.59057 |
| N | 1.52604 | –0.59267 | 1.01715 |
| C | 1.67308 | 0.68387 | 0.59110 |
| C | –1.21726 | –0.63707 | –0.45904 |
| N | –2.33726 | 0.10003 | –1.01201 |
| C | –6.42342 | –0.85785 | –0.53095 |
| C | –5.43540 | –1.81951 | –0.28493 |
| C | –4.08506 | –1.52512 | –0.41366 |
| C | –3.66708 | –0.23383 | –0.79587 |
| C | –4.66119 | 0.74274 | –1.01916 |
| C | –6.00603 | 0.43489 | –0.88872 |
| C | –7.84941 | –1.26833 | –0.37980 |
| O | –8.17957 | –2.31777 | 0.16469 |
| N | –8.79955 | –0.39122 | –0.86462 |
| C | –1.44037 | 1.91163 | 1.97606 |
| O | –2.46370 | 1.51711 | 2.51005 |
| H | 5.83322 | –4.15680 | 0.59455 |
| H | 7.94256 | –3.37932 | –0.53589 |
| H | 4.32118 | 3.46058 | –1.45636 |
| H | –1.80752 | –0.50030 | 1.62436 |
| H | 0.18963 | –2.14720 | 1.36772 |

| | | | |
|---|---|---|---|
| H | 0.27697 | –0.96888 | 2.68331 |
| H | –0.33569 | –0.41619 | –1.06961 |
| H | –1.39858 | –1.71036 | –0.55878 |
| H | –2.16850 | 1.09680 | –1.06535 |
| H | –5.75630 | –2.81183 | 0.01384 |
| H | –3.35707 | –2.30429 | –0.21747 |
| H | –4.36053 | 1.75325 | –1.28370 |
| H | –6.73352 | 1.22894 | –1.03141 |
| H | –8.55988 | 0.24538 | –1.60872 |
| H | –9.73909 | –0.76130 | –0.87840 |
| H | –1.06132 | 2.93979 | 2.06842 |
| H | 2.14858 | 4.04921 | –0.84244 |
| H | 2.32853 | –1.22058 | 0.98148 |
| H | 4.99878 | 1.93185 | –1.01003 |
| H | 8.05095 | –1.14705 | –1.33563 |
| H | 4.28125 | –0.25290 | –0.07921 |

**Table S42.** Optimized geometry coordinates of U:GLY (W–edge)
Energy: –438803.9211738 (au)

| | | | |
|---|---|---|---|
| N | 2.57510 | –1.39191 | –0.39669 |
| C | 1.23148 | –1.05043 | –0.27178 |
| O | 0.34563 | –1.87953 | –0.44274 |
| N | 1.02015 | 0.26896 | 0.05499 |
| C | 1.99882 | 1.26708 | 0.26430 |
| O | 1.67349 | 2.40792 | 0.54604 |
| C | 3.37229 | 0.78935 | 0.11063 |
| C | 3.60590 | –0.49954 | –0.21013 |
| N | –2.32001 | –1.53789 | 1.13220 |
| C | –3.31205 | –0.83687 | 0.35206 |
| C | –2.89819 | 0.51876 | –0.20996 |
| O | –1.80253 | 1.03702 | –0.10537 |
| O | –3.91225 | 1.12778 | –0.85781 |
| H | –3.62411 | –1.45403 | –0.49966 |
| H | –4.22362 | –0.67191 | 0.94060 |
| H | 4.17598 | 1.49686 | 0.25828 |
| H | 4.60298 | –0.90601 | –0.33840 |
| H | 0.04015 | 0.56109 | 0.11718 |
| H | –3.56602 | 1.97726 | –1.18183 |
| H | 2.74795 | –2.35874 | –0.62923 |
| H | –1.49680 | –1.74092 | 0.56510 |
| H | –2.00741 | –0.95332 | 1.90287 |

**Table S43.** Optimized geometry coordinates of 1U:H4B (W–edge)
Energy: –794623.5812435 (au)

| | | | |
|---|---|---|---|
| N | –4.65927 | –2.12282 | –0.09029 |
| C | –3.40205 | –1.55861 | –0.28934 |

| | | | |
|---|---|---|---|
| O | –2.47423 | –2.22067 | –0.73869 |
| N | –3.31569 | –0.23073 | 0.05982 |
| C | –4.34551 | 0.56007 | 0.57741 |
| O | –4.15385 | 1.74505 | 0.85139 |
| C | –5.61942 | –0.12328 | 0.75480 |
| C | –5.72864 | –1.42567 | 0.41569 |
| N | –0.73007 | 1.04772 | –0.32226 |
| C | –0.60519 | 2.34723 | –0.14184 |
| N | –1.70256 | 3.12623 | 0.09485 |
| N | 0.60465 | 2.96968 | –0.23314 |
| C | 1.82454 | 2.30705 | –0.52547 |
| O | 2.88698 | 2.94289 | –0.51783 |
| C | 1.65954 | 0.92152 | –0.78953 |
| C | 0.40722 | 0.33784 | –0.64924 |
| N | 2.77339 | 0.15529 | –1.17914 |
| N | 0.27955 | –1.01573 | –0.78037 |
| C | 2.73409 | –1.25622 | –0.83965 |
| C | 1.37898 | –1.80950 | –1.31901 |
| C | 2.98164 | –1.59094 | 0.65184 |
| O | 3.07368 | –3.02032 | 0.69052 |
| C | 4.25538 | –0.94158 | 1.24603 |
| C | 4.02827 | 0.42008 | 1.90885 |
| O | 5.23661 | –0.85883 | 0.20894 |
| H | –6.45190 | 0.43919 | 1.15244 |
| H | –6.64896 | –1.98929 | 0.51962 |
| H | –1.55150 | 4.00860 | 0.56016 |
| H | 3.65881 | 0.61456 | –1.00734 |
| H | 3.52498 | –1.76402 | –1.39838 |
| H | 1.37581 | –1.80248 | –2.41771 |
| H | 1.26412 | –2.84072 | –0.98046 |
| H | 2.11919 | –1.25463 | 1.24682 |
| H | 3.06254 | –3.29642 | 1.61569 |
| H | 3.25854 | 0.35341 | 2.68551 |
| H | 4.95172 | 0.75634 | 2.39717 |
| H | 3.73347 | 1.18696 | 1.18882 |
| H | –2.56118 | 2.64054 | 0.36320 |
| H | –0.65565 | –1.41180 | –0.80320 |
| H | –2.38144 | 0.22340 | –0.08263 |
| H | –4.73440 | –3.09736 | –0.34447 |
| H | 6.02170 | –0.43488 | 0.57872 |
| H | 4.60736 | –1.64135 | 2.02515 |
| H | 0.69004 | 3.96941 | –0.10340 |

**Table S44.** Optimized geometry coordinates of U:LYA (W–edge)
Energy: –890902.9370460 (au)

| | | | |
|---|---|---|---|
| N | –6.23052 | –2.69261 | 0.11588 |

| | | | |
|---|---|---|---|
| C | –5.09689 | –1.93808 | –0.15562 |
| O | –4.10474 | –2.42086 | –0.68853 |
| N | –5.21063 | –0.61948 | 0.22437 |
| C | –6.34611 | 0.02993 | 0.73777 |
| O | –6.32855 | 1.24097 | 0.93180 |
| C | –7.47385 | –0.85623 | 0.98804 |
| C | –7.37417 | –2.16564 | 0.67093 |
| C | 4.21696 | –0.89256 | –0.31189 |
| C | 5.04286 | –0.21420 | –1.21899 |
| C | 6.39059 | 0.00049 | –0.94181 |
| C | 6.95141 | –0.46440 | 0.25598 |
| C | 6.13721 | –1.16124 | 1.15676 |
| C | 4.78926 | –1.36406 | 0.87796 |
| C | 2.74559 | –1.08457 | –0.60217 |
| C | 1.90251 | 0.15164 | –0.23459 |
| C | 0.43545 | 0.00102 | –0.52496 |
| C | –0.23785 | –1.10437 | –0.99058 |
| N | –1.59212 | –0.79739 | –1.13993 |
| C | –1.76095 | 0.49272 | –0.74906 |
| C | –0.55593 | 1.03288 | –0.36025 |
| C | –0.51027 | 2.41142 | 0.12477 |
| O | 0.52172 | 3.00260 | 0.41128 |
| N | –1.77226 | 3.04120 | 0.29086 |
| C | –2.86482 | 2.48541 | –0.11026 |
| N | –2.98014 | 1.17950 | –0.67277 |
| N | –4.07210 | 3.13531 | –0.06290 |
| C | 8.39385 | –0.27568 | 0.62249 |
| O | 8.94966 | –0.96930 | 1.46593 |
| N | 9.07139 | 0.70883 | –0.06158 |
| H | –8.37464 | –0.43699 | 1.41316 |
| H | –8.18239 | –2.87110 | 0.82639 |
| H | 4.62562 | 0.14154 | –2.15773 |
| H | 7.01119 | 0.49846 | –1.68121 |
| H | 6.58594 | –1.53664 | 2.07017 |
| H | 4.16848 | –1.90108 | 1.59082 |
| H | 2.37493 | –1.95484 | –0.04726 |
| H | 2.60704 | –1.30649 | –1.66779 |
| H | 2.04082 | 0.38512 | 0.82855 |
| H | 2.28527 | 1.03271 | –0.76243 |
| H | 0.11943 | –2.08903 | –1.24940 |
| H | –2.34664 | –1.47109 | –1.23654 |
| H | –4.00188 | 4.03275 | 0.39797 |
| H | 8.56345 | 1.48107 | –0.46454 |
| H | 9.99607 | 0.91096 | 0.29091 |
| H | –6.17340 | –3.66640 | –0.14645 |
| H | –4.38437 | –0.02952 | 0.07220 |

| | | | |
|---|---|---|---|
| H | –4.88059 | 2.57968 | 0.21498 |
| H | –3.54344 | 1.17909 | –1.52022 |

**Table S45.** Optimized geometry coordinates of U:PRF (W–edge)
Energy: –650134.4750884 (au)

| | | | |
|---|---|---|---|
| N | 4.25488 | –1.77228 | 0.03158 |
| C | 2.90305 | –1.44826 | –0.03492 |
| O | 2.04556 | –2.32151 | –0.13314 |
| N | 2.64427 | –0.10148 | 0.02016 |
| C | 3.58293 | 0.92737 | 0.13630 |
| O | 3.22107 | 2.10396 | 0.17911 |
| C | 4.96996 | 0.48801 | 0.19934 |
| C | 5.25052 | –0.83212 | 0.14439 |
| N | –1.82825 | 2.23659 | –0.10683 |
| C | –0.49243 | 1.92342 | –0.12237 |
| N | –0.06889 | 0.67529 | –0.13506 |
| C | –1.06019 | –0.26455 | –0.15574 |
| C | –2.44495 | –0.04523 | –0.14681 |
| C | –2.91413 | 1.30566 | –0.10198 |
| O | –4.06128 | 1.74547 | –0.05830 |
| C | –3.08792 | –1.33497 | –0.16434 |
| C | –4.57082 | –1.60950 | –0.17978 |
| N | –5.36355 | –0.96289 | 0.87406 |
| C | –2.06897 | –2.25635 | –0.17472 |
| N | –0.83967 | –1.60345 | –0.17476 |
| N | 0.40575 | 2.94676 | –0.17335 |
| H | 5.74468 | 1.23625 | 0.28801 |
| H | 6.26067 | –1.22362 | 0.18567 |
| H | –4.99345 | –1.28509 | –1.13913 |
| H | –4.72909 | –2.69410 | –0.12946 |
| H | –5.27994 | 0.04663 | 0.74757 |
| H | –2.11382 | –3.33536 | –0.19374 |
| H | 0.08922 | –2.02201 | –0.16861 |
| H | 0.12494 | 3.84741 | 0.18338 |
| H | 4.46213 | –2.75976 | –0.00942 |
| H | 1.63480 | 0.18743 | –0.03593 |
| H | –4.93917 | –1.16758 | 1.77741 |
| H | 1.38463 | 2.69088 | –0.02219 |
| H | –2.12230 | 3.20483 | –0.13180 |

**Table S46.** Optimized geometry coordinates of U:SAH (W–edge)
Energy: –1294701.6828546 (au)

| | | | |
|---|---|---|---|
| N | 5.81003 | –1.50904 | –1.97943 |
| C | 4.48667 | –1.53677 | –1.53487 |
| O | 3.61389 | –2.14604 | –2.13536 |
| N | 4.27873 | –0.82040 | –0.37699 |

| | | | |
|---|---|---|---|
| C | 5.22385 | –0.10387 | 0.35660 |
| O | 4.90077 | 0.49772 | 1.38136 |
| C | 6.57413 | –0.13927 | –0.19113 |
| C | 6.81166 | –0.83335 | –1.32572 |
| N | 0.15553 | 1.90413 | –1.63573 |
| C | –0.66078 | 3.06891 | –1.32173 |
| C | –2.08392 | 2.64051 | –0.88343 |
| C | –2.82267 | 1.87964 | –1.98558 |
| S | –4.56020 | 1.43082 | –1.57354 |
| C | –0.02776 | 3.95629 | –0.23106 |
| O | –0.58092 | 5.18633 | –0.05775 |
| O | 0.89456 | 3.60339 | 0.46144 |
| C | –4.38318 | 0.23839 | –0.18902 |
| C | –3.88131 | –1.15002 | –0.55339 |
| O | –2.52109 | –1.06928 | –1.03917 |
| C | –3.83480 | –2.11913 | 0.65328 |
| O | –5.03322 | –2.82728 | 0.86322 |
| C | –2.64097 | –3.02742 | 0.29633 |
| O | –3.08262 | –4.13675 | –0.49353 |
| C | –1.71112 | –2.10818 | –0.52316 |
| N | –0.56425 | –1.55359 | 0.18735 |
| C | 0.72805 | –1.57882 | –0.29379 |
| N | 1.57194 | –0.90753 | 0.45057 |
| C | 0.80630 | –0.38696 | 1.48108 |
| C | 1.12389 | 0.42862 | 2.59732 |
| N | 2.36437 | 0.85180 | 2.87729 |
| N | 0.11381 | 0.77657 | 3.42731 |
| C | –1.12061 | 0.34523 | 3.15705 |
| N | –1.54305 | –0.42285 | 2.14151 |
| C | –0.52979 | –0.76455 | 1.33341 |
| H | 7.35626 | 0.39571 | 0.32852 |
| H | 7.79340 | –0.89709 | –1.78185 |
| H | 1.01687 | 2.21099 | –2.08258 |
| H | –0.75118 | 3.67630 | –2.23510 |
| H | –1.98699 | 2.01152 | 0.00882 |
| H | –2.66901 | 3.51813 | –0.58344 |
| H | –2.26578 | 0.98808 | –2.26785 |
| H | –2.93016 | 2.50978 | –2.87610 |
| H | –5.39590 | 0.13659 | 0.21471 |
| H | –3.75191 | 0.65744 | 0.59931 |
| H | –4.51655 | –1.57493 | –1.34516 |
| H | –3.59428 | –1.55564 | 1.55742 |
| H | –4.94246 | –3.65813 | 0.36840 |
| H | –2.14825 | –3.45987 | 1.16780 |
| H | –3.34447 | –3.80958 | –1.36691 |
| H | –1.27440 | –2.68725 | –1.34608 |

| | | | |
|---|---|---|---|
| H | 1.00083 | –2.10590 | –1.19871 |
| H | 3.16269 | 0.68656 | 2.26536 |
| H | –1.88995 | 0.66182 | 3.85835 |
| H | 5.98897 | –2.02853 | –2.82634 |
| H | 3.29817 | –0.82281 | –0.02164 |
| H | 2.46382 | 1.49400 | 3.64826 |
| H | 0.45870 | 1.48584 | –0.75645 |
| H | –1.30205 | 5.31985 | –0.69002 |

**Table S47.** Optimized geometry coordinates of U:SFG (W–edge)
Energy: –1104243.3060471 (au)

| | | | |
|---|---|---|---|
| N | 4.78808 | 0.63612 | –2.28074 |
| C | 3.62268 | –0.11504 | –2.13587 |
| O | 2.77723 | –0.18657 | –3.01458 |
| N | 3.53791 | –0.76975 | –0.92442 |
| C | 4.38736 | –0.62692 | 0.16354 |
| O | 4.11293 | –1.16103 | 1.24863 |
| C | 5.56661 | 0.18473 | –0.07916 |
| C | 5.71238 | 0.79098 | –1.27964 |
| N | 1.32927 | 2.21344 | –0.84964 |
| C | 1.00992 | 2.69872 | 0.50194 |
| C | 1.99293 | 2.01866 | 1.45219 |
| O | 2.82868 | 2.57570 | 2.12699 |
| O | 1.83159 | 0.67368 | 1.43003 |
| C | –0.45197 | 2.50107 | 0.97296 |
| C | –1.47155 | 3.18094 | 0.04786 |
| C | –2.94227 | 3.11239 | 0.51243 |
| N | –3.09697 | 3.91482 | 1.73716 |
| C | –3.51562 | 1.68249 | 0.66229 |
| C | –3.66484 | 0.83813 | –0.59840 |
| O | –2.37170 | 0.49938 | –1.15597 |
| C | –4.36418 | –0.51762 | –0.34581 |
| O | –5.76944 | –0.45355 | –0.41709 |
| C | –3.73963 | –1.43278 | –1.41844 |
| O | –4.50103 | –1.37506 | –2.62945 |
| C | –2.33116 | –0.84278 | –1.62438 |
| N | –1.23203 | –1.54703 | –0.97683 |
| C | 0.06984 | –1.46492 | –1.43155 |
| N | 0.95510 | –1.87906 | –0.55976 |
| C | 0.21604 | –2.23951 | 0.55365 |
| C | 0.58050 | –2.70267 | 1.84005 |
| N | 1.86035 | –2.87458 | 2.22977 |
| N | –0.40918 | –2.99148 | 2.71187 |
| C | –1.67695 | –2.78672 | 2.33728 |
| N | –2.14185 | –2.30083 | 1.17848 |
| C | –1.14639 | –2.04132 | 0.32202 |

| | | | |
|---|---|---|---|
| H | 6.28192 | 0.31442 | 0.72016 |
| H | 6.55582 | 1.42998 | –1.51583 |
| H | 0.82248 | 2.75903 | –1.54115 |
| H | 1.02027 | 1.24911 | –0.94846 |
| H | 1.25892 | 3.76387 | 0.53773 |
| H | –0.65248 | 1.42626 | 1.03319 |
| H | –0.53377 | 2.90508 | 1.99096 |
| H | –1.40164 | 2.73120 | –0.94684 |
| H | –1.22064 | 4.24481 | –0.04607 |
| H | –3.53571 | 3.62360 | –0.25927 |
| H | –4.08346 | 3.98400 | 1.97926 |
| H | –2.65555 | 3.43599 | 2.52033 |
| H | –4.52812 | 1.77124 | 1.07929 |
| H | –2.92812 | 1.11176 | 1.39276 |
| H | –4.23820 | 1.39994 | –1.35506 |
| H | –4.07903 | –0.89478 | 0.63959 |
| H | –6.00108 | –0.70986 | –1.32495 |
| H | –3.72131 | –2.48141 | –1.12306 |
| H | –4.39058 | –0.49795 | –3.02533 |
| H | –2.08695 | –0.85288 | –2.69373 |
| H | 4.86275 | 1.14707 | –3.14844 |
| H | 2.64966 | –1.28890 | –0.76812 |
| H | –2.43075 | –3.03942 | 3.07986 |
| H | 0.31665 | –1.08362 | –2.41436 |
| H | 2.62612 | –2.42127 | 1.73447 |
| H | 1.99836 | –3.10578 | 3.20242 |
| H | 2.57140 | 0.25951 | 1.91335 |

**Table S48.** Optimized geometry coordinates of U:THF (W–edge)
Energy: –650888.7116509 (au)

| | | | |
|---|---|---|---|
| N | –4.11136 | –1.72374 | –0.08635 |
| C | –2.75956 | –1.39354 | –0.05663 |
| O | –1.89485 | –2.26125 | –0.09860 |
| N | –2.50724 | –0.04366 | 0.02082 |
| C | –3.46163 | 0.97758 | 0.06294 |
| O | –3.11953 | 2.15815 | 0.12874 |
| C | –4.84705 | 0.53005 | 0.02297 |
| C | –5.11807 | –0.79055 | –0.04855 |
| N | 0.29387 | 0.76222 | 0.05198 |
| C | 0.61434 | 2.03831 | 0.12589 |
| N | –0.33671 | 2.98204 | 0.39337 |
| N | 1.90301 | 2.46411 | –0.00667 |
| C | 3.01299 | 1.60469 | –0.22083 |
| O | 4.15239 | 2.07594 | –0.28294 |
| C | 2.64363 | 0.23475 | –0.32699 |
| N | 3.64607 | –0.72377 | –0.59908 |

| | | | |
|---|---|---|---|
| C | 3.37778 | –2.06043 | –0.06688 |
| C | 1.96995 | –2.45866 | –0.55590 |
| N | 0.99009 | –1.46484 | –0.14445 |
| C | 1.32016 | –0.13893 | –0.15048 |
| C | 3.51713 | –2.17398 | 1.46007 |
| H | –5.62763 | 1.27673 | 0.05387 |
| H | –6.12662 | –1.18715 | –0.07980 |
| H | –0.15195 | 3.92333 | 0.08023 |
| H | 4.09174 | –2.74774 | –0.53800 |
| H | 1.99756 | –2.57005 | –1.64930 |
| H | 1.68280 | –3.42412 | –0.12812 |
| H | 3.34873 | –3.20370 | 1.79580 |
| H | 4.52340 | –1.87841 | 1.77503 |
| H | 2.79752 | –1.52697 | 1.96894 |
| H | 4.56074 | –0.34973 | –0.36546 |
| H | 0.00886 | –1.72556 | –0.11801 |
| H | –1.31186 | 2.68674 | 0.31165 |
| H | 2.14614 | 3.43652 | 0.13136 |
| H | –1.49789 | 0.23770 | 0.04650 |
| H | –4.30771 | –2.71262 | –0.14328 |

**Table S49.** Optimized geometry coordinates of 1U:N6M (W–edge)
Energy: –578244.6719088 (au)

| | | | |
|---|---|---|---|
| N | 4.41533 | –1.11718 | 0.00384 |
| C | 3.03275 | –1.32778 | –0.00157 |
| O | 2.55395 | –2.45022 | –0.00406 |
| N | 2.29599 | –0.16263 | –0.00361 |
| C | 2.78737 | 1.14129 | 0.00057 |
| O | 2.01734 | 2.10486 | –0.00047 |
| C | 4.24031 | 1.25237 | 0.00626 |
| C | 4.99050 | 0.12863 | 0.00768 |
| C | –1.43016 | 0.55326 | –0.01454 |
| C | –2.80655 | 0.19341 | –0.00372 |
| C | –3.08085 | –1.18219 | 0.00646 |
| C | –0.95309 | –1.74616 | 0.00023 |
| C | –4.93455 | 0.02524 | 0.00116 |
| C | –1.69059 | 3.03633 | 0.02351 |
| N | –0.92999 | 1.79941 | –0.03873 |
| N | –0.53309 | –0.47071 | –0.01010 |
| N | –2.20624 | –2.19812 | 0.00912 |
| N | –4.45399 | –1.26652 | 0.01006 |
| N | –3.98592 | 0.92750 | –0.00719 |
| H | 4.68223 | 2.23858 | 0.00928 |
| H | 6.07490 | 0.14427 | 0.01188 |
| H | –0.15718 | –2.48805 | 0.00200 |
| H | –5.99528 | 0.23630 | 0.00192 |

| | | | |
|---|---|---|---|
| H | –2.74157 | 2.84436 | –0.18634 |
| H | –1.61270 | 3.49938 | 1.01572 |
| H | –1.29678 | 3.74303 | –0.71357 |
| H | –4.98640 | –2.12326 | 0.01766 |
| H | 4.97422 | –1.95787 | 0.00554 |
| H | 1.25246 | –0.27663 | –0.00788 |
| H | 0.09030 | 1.87403 | –0.01984 |

**Table S50.** Optimized geometry coordinates of 2U:N6M (W–edge)
Energy: –578244.6719088 (au)

| | | | |
|---|---|---|---|
| N | 4.41533 | –1.11718 | 0.00384 |
| C | 3.03275 | –1.32778 | –0.00157 |
| O | 2.55395 | –2.45022 | –0.00406 |
| N | 2.29599 | –0.16263 | –0.00361 |
| C | 2.78737 | 1.14129 | 0.00057 |
| O | 2.01734 | 2.10486 | –0.00047 |
| C | 4.24031 | 1.25237 | 0.00626 |
| C | 4.99050 | 0.12863 | 0.00768 |
| C | –1.43016 | 0.55326 | –0.01454 |
| C | –2.80655 | 0.19341 | –0.00372 |
| C | –3.08085 | –1.18219 | 0.00646 |
| C | –0.95309 | –1.74616 | 0.00023 |
| C | –4.93455 | 0.02524 | 0.00116 |
| C | –1.69059 | 3.03633 | 0.02351 |
| N | –0.92999 | 1.79941 | –0.03873 |
| N | –0.53309 | –0.47071 | –0.01010 |
| N | –2.20624 | –2.19812 | 0.00912 |
| N | –4.45399 | –1.26652 | 0.01006 |
| N | –3.98592 | 0.92750 | –0.00719 |
| H | 4.68223 | 2.23858 | 0.00928 |
| H | 6.07490 | 0.14427 | 0.01188 |
| H | –0.15718 | –2.48805 | 0.00200 |
| H | –5.99528 | 0.23630 | 0.00192 |
| H | –2.74157 | 2.84436 | –0.18634 |
| H | –1.61270 | 3.49938 | 1.01572 |
| H | –1.29678 | 3.74303 | –0.71357 |
| H | –4.98640 | –2.12326 | 0.01766 |
| H | 4.97422 | –1.95787 | 0.00554 |
| H | 1.25246 | –0.27663 | –0.00788 |
| H | 0.09030 | 1.87403 | –0.01984 |

**Table S51.** Optimized geometry coordinates of U:XAN (W–edge)
Energy: –613260.4982540 (au)

| | | | |
|---|---|---|---|
| N | –4.93047 | –0.20485 | 0.01302 |
| C | –3.79449 | –1.02385 | 0.00664 |
| O | –3.87412 | –2.23483 | 0.01013 |

| | | | |
|---|---|---|---|
| N | –2.60263 | –0.31109 | –0.00384 |
| C | –2.46300 | 1.07187 | –0.00865 |
| O | –1.34363 | 1.60513 | –0.01777 |
| C | –3.70390 | 1.82546 | –0.00239 |
| C | –4.88250 | 1.16243 | 0.00802 |
| N | 2.58885 | 2.12333 | 0.00209 |
| C | 2.28288 | 0.79500 | –0.00114 |
| N | 1.04589 | 0.21434 | –0.00806 |
| C | 0.95099 | –1.16183 | –0.00994 |
| O | –0.13447 | –1.74841 | –0.01565 |
| N | 2.15551 | –1.83748 | –0.00507 |
| C | 3.49499 | –1.33142 | 0.00269 |
| O | 4.43500 | –2.10182 | 0.00690 |
| C | 3.48982 | 0.11873 | 0.00425 |
| N | 4.54110 | 1.01720 | 0.01102 |
| C | 3.97983 | 2.19453 | 0.00958 |
| H | –3.65049 | 2.90454 | –0.00609 |
| H | –5.84170 | 1.66827 | 0.01327 |
| H | 1.93435 | 2.89188 | 0.00049 |
| H | 4.49232 | 3.14605 | 0.01329 |
| H | –1.73956 | –0.88476 | –0.00872 |
| H | 0.15834 | 0.74860 | –0.01305 |
| H | 2.07761 | –2.84723 | –0.00614 |
| H | –5.80869 | –0.70364 | 0.01968 |

**Table S52.** Optimized geometry coordinates of U:PQ0 (H–edge)
Energy: –648600.1797040 (au)

| | | | |
|---|---|---|---|
| N | –4.97667 | –0.87795 | 0.44265 |
| C | –5.30185 | 0.44485 | 0.14770 |
| O | –6.43789 | 0.87316 | 0.17978 |
| N | –4.18907 | 1.20743 | –0.18216 |
| C | –2.85211 | 0.79819 | –0.23773 |
| O | –1.98436 | 1.61915 | –0.53859 |
| C | –2.63527 | –0.60335 | 0.07941 |
| C | –3.69657 | –1.37450 | 0.40687 |
| N | 2.97744 | –2.15763 | 0.01992 |
| C | 1.62807 | –2.27219 | –0.18782 |
| N | 0.82361 | –1.24454 | –0.30183 |
| C | 1.45756 | –0.03948 | –0.19092 |
| C | 2.82199 | 0.19334 | 0.00997 |
| C | 3.70856 | –0.92965 | 0.14222 |
| O | 4.91196 | –0.96781 | 0.33530 |
| C | 3.00203 | 1.61953 | 0.04694 |
| C | 4.20499 | 2.35122 | 0.22867 |
| N | 5.17422 | 2.97910 | 0.37530 |
| C | 1.74734 | 2.17425 | –0.13322 |

| | | | |
|---|---|---|---|
| N | 0.82466 | 1.16792 | –0.27639 |
| N | 1.11334 | –3.55228 | –0.22729 |
| H | –1.62453 | –0.99394 | 0.04013 |
| H | –3.59984 | –2.42453 | 0.65949 |
| H | 1.45534 | 3.21229 | –0.16962 |
| H | –0.18384 | 1.29822 | –0.41594 |
| H | 0.17204 | –3.57896 | –0.59414 |
| H | –5.76179 | –1.46243 | 0.69201 |
| H | –4.37448 | 2.17988 | –0.39872 |
| H | 1.71236 | –4.27043 | –0.61149 |
| H | 3.54576 | –2.98359 | 0.16232 |

**Table S53.** Optimized geometry coordinates of C:29G (W–edge)
Energy: –599759.3753018 (au)

| | | | |
|---|---|---|---|
| N | 5.11240 | –0.24336 | –0.01340 |
| C | 3.90306 | –0.99080 | –0.00962 |
| O | 3.96581 | –2.21327 | –0.01785 |
| N | 2.74254 | –0.26725 | 0.00318 |
| C | 2.75620 | 1.06741 | 0.01109 |
| N | 1.57730 | 1.70442 | 0.02407 |
| C | 3.98749 | 1.82442 | 0.00654 |
| C | 5.14359 | 1.11502 | –0.00591 |
| N | –4.23639 | –2.32592 | –0.01935 |
| C | –3.13725 | –1.52407 | 0.00519 |
| N | –1.95910 | –2.11326 | 0.00940 |
| C | –3.26396 | –0.08370 | –0.00594 |
| C | –4.44597 | 0.66412 | –0.04314 |
| N | –4.46539 | 1.99237 | –0.03806 |
| C | –3.24548 | 2.58059 | 0.00097 |
| N | –2.05626 | 2.00852 | 0.02194 |
| C | –2.03731 | 0.64490 | 0.01291 |
| N | –0.84075 | 0.03739 | 0.01776 |
| C | –0.84840 | –1.31027 | 0.00363 |
| N | 0.33031 | –1.93680 | –0.00662 |
| H | 1.53695 | 2.70978 | 0.02720 |
| H | 3.98890 | 2.90664 | 0.01266 |
| H | 6.12324 | 1.58126 | –0.01042 |
| H | –5.13952 | –1.96813 | 0.24235 |
| H | –5.42021 | 0.17537 | –0.08899 |
| H | 1.21294 | –1.40340 | –0.00370 |
| H | 0.33590 | –2.94351 | –0.01490 |
| H | 0.69692 | 1.16634 | 0.02457 |
| H | –4.06836 | –3.31144 | 0.11815 |
| H | –3.25633 | 3.66969 | 0.01229 |
| H | 5.95856 | –0.79483 | –0.02325 |

**Table S54.** Optimized geometry coordinates of 1C:29H (W–edge)
Energy: –612230.4387673 (au)

| | | | |
|---|---|---|---|
| N | –5.06545 | –0.23991 | 0.00151 |
| C | –3.84765 | –0.96748 | 0.00009 |
| O | –3.88294 | –2.19152 | –0.00054 |
| N | –2.69854 | –0.22542 | –0.00028 |
| C | –2.72752 | 1.10970 | –0.00045 |
| N | –1.55635 | 1.75964 | –0.00108 |
| C | –3.96987 | 1.84607 | –0.00016 |
| C | –5.11506 | 1.11847 | 0.00065 |
| N | 1.93316 | –2.07517 | –0.00074 |
| C | 3.24794 | –1.57495 | 0.00001 |
| O | 4.20174 | –2.33639 | 0.00043 |
| C | 3.28422 | –0.11434 | 0.00003 |
| C | 4.47780 | 0.60908 | 0.00061 |
| N | 4.50838 | 1.94152 | 0.00063 |
| C | 3.29898 | 2.53851 | 0.00006 |
| N | 2.09795 | 1.97479 | –0.00046 |
| C | 2.07063 | 0.61938 | –0.00017 |
| N | 0.84497 | 0.02987 | –0.00008 |
| C | 0.79274 | –1.29491 | –0.00022 |
| N | –0.39302 | –1.90665 | 0.00028 |
| H | –1.52542 | 2.76542 | –0.00038 |
| H | –3.98856 | 2.92802 | –0.00055 |
| H | –6.10128 | 1.57021 | 0.00101 |
| H | 5.42702 | 0.07747 | 0.00091 |
| H | –1.25708 | –1.32553 | –0.00024 |
| H | –0.67331 | 1.23027 | –0.00059 |
| H | –5.90473 | –0.80221 | 0.00053 |
| H | –0.47650 | –2.91000 | –0.00069 |
| H | 3.31406 | 3.62749 | 0.00005 |
| H | 1.86444 | –3.08576 | –0.00035 |

**Table S55.** Optimized geometry coordinates of 2C:29H (W–edge)
Energy: –612238.7622669 (au)

| | | | |
|---|---|---|---|
| N | –4.58971 | 0.84817 | –0.00258 |
| C | –3.19254 | 1.03112 | –0.00051 |
| O | –2.75252 | 2.18423 | –0.00061 |
| N | –2.42066 | –0.08915 | 0.00147 |
| C | –2.96957 | –1.30988 | 0.00112 |
| N | –2.14450 | –2.36283 | 0.00306 |
| C | –4.39985 | –1.50017 | –0.00120 |
| C | –5.16896 | –0.38288 | –0.00300 |
| N | 0.45045 | 0.32048 | 0.00192 |
| C | 1.20607 | –0.83625 | 0.00085 |
| O | 0.66862 | –1.95422 | 0.00084 |

| | | | |
|---|---|---|---|
| C | 2.64011 | –0.59394 | –0.00018 |
| C | 3.58211 | –1.62617 | –0.00099 |
| N | 4.89300 | –1.40008 | –0.00193 |
| C | 5.24146 | –0.09263 | –0.00204 |
| N | 4.45918 | 0.97461 | –0.00131 |
| C | 3.12085 | 0.74569 | –0.00031 |
| N | 2.30005 | 1.82797 | 0.00050 |
| C | 1.00280 | 1.58711 | 0.00159 |
| N | 0.13063 | 2.60891 | 0.00280 |
| H | –1.12140 | –2.22702 | 0.00453 |
| H | –4.84125 | –2.48810 | –0.00148 |
| H | –6.25292 | –0.40993 | –0.00478 |
| H | 3.24357 | –2.66042 | –0.00086 |
| H | 0.52949 | 3.53369 | 0.00157 |
| H | 6.31322 | 0.10250 | –0.00284 |
| H | –0.58090 | 0.20440 | 0.00231 |
| H | –0.88422 | 2.47918 | 0.00170 |
| H | –2.51744 | –3.29793 | 0.00260 |
| H | –5.14093 | 1.69519 | –0.00409 |

**Table S56.** Optimized geometry coordinates of C:2BP (W–edge)
Energy: –541098.3482327 (au)

| | | | |
|---|---|---|---|
| N | 4.87960 | –0.50210 | –0.00040 |
| C | 3.59018 | –1.10450 | 0.00057 |
| O | 3.51141 | –2.32449 | 0.00120 |
| N | 2.52280 | –0.24685 | 0.00069 |
| C | 2.69704 | 1.07279 | 0.00004 |
| N | 1.59676 | 1.84527 | 0.00030 |
| C | 4.00603 | 1.68309 | –0.00082 |
| C | 5.07118 | 0.84200 | –0.00098 |
| N | –4.82686 | –0.93726 | –0.00083 |
| C | –5.51626 | 0.26901 | –0.00014 |
| N | –4.74115 | 1.31893 | 0.00053 |
| C | –3.45508 | 0.78893 | 0.00037 |
| C | –2.18079 | 1.34593 | 0.00084 |
| N | –1.09487 | 0.57120 | 0.00056 |
| C | –1.26168 | –0.79188 | –0.00017 |
| N | –0.14426 | –1.53890 | –0.00048 |
| N | –2.44969 | –1.44445 | –0.00069 |
| C | –3.48804 | –0.62347 | –0.00041 |
| H | 1.68329 | 2.84756 | –0.00063 |
| H | 4.13480 | 2.75766 | –0.00132 |
| H | 6.09895 | 1.18956 | –0.00162 |
| H | –5.21683 | –1.86737 | –0.00087 |
| H | –6.59806 | 0.29246 | –0.00028 |
| H | –2.02661 | 2.42358 | 0.00144 |

| | | | |
|---|---|---|---|
| H | 0.79189 | –1.11977 | 0.00054 |
| H | 5.65505 | –1.14957 | –0.00028 |
| H | 0.66211 | 1.40926 | 0.00054 |
| H | –0.24764 | –2.53980 | –0.00061 |

**Table S57.** Optimized geometry coordinates of 1C:5AZ (W–edge)
Energy: –505747.7955300 (au)

| | | | |
|---|---|---|---|
| N | –4.20039 | –0.73254 | –0.00213 |
| C | –2.85842 | –1.19366 | 0.00001 |
| O | –2.63864 | –2.39696 | 0.00041 |
| N | –1.88864 | –0.22689 | 0.00168 |
| C | –2.19516 | 1.07013 | 0.00134 |
| N | –1.17927 | 1.94812 | 0.00389 |
| C | –3.56088 | 1.53532 | –0.00117 |
| C | –4.53052 | 0.58518 | –0.00274 |
| C | 4.21936 | –0.19654 | –0.00104 |
| N | 3.04680 | –1.01512 | –0.00009 |
| C | 1.78008 | –0.51518 | 0.00084 |
| N | 1.59266 | 0.81205 | –0.00006 |
| C | 2.73007 | 1.54934 | –0.00082 |
| N | 3.98142 | 1.16321 | –0.00115 |
| N | 0.73485 | –1.34028 | 0.00276 |
| O | 5.30625 | –0.73911 | –0.00176 |
| H | –0.20789 | 1.60633 | 0.00284 |
| H | –3.80389 | 2.58975 | –0.00142 |
| H | –5.58892 | 0.82272 | –0.00461 |
| H | 2.56316 | 2.62732 | –0.00139 |
| H | 0.85577 | –2.34048 | 0.00216 |
| H | –1.36311 | 2.93738 | 0.00074 |
| H | 3.22070 | –2.01271 | 0.00050 |
| H | –4.90548 | –1.45627 | –0.00321 |
| H | –0.23037 | –0.95285 | 0.00242 |

**Table S58.** Optimized geometry coordinates of 2C:5AZ (W–edge)
Energy: –505760.2264373 (au)

| | | | |
|---|---|---|---|
| N | 3.40582 | –1.18700 | –0.00005 |
| C | 1.99664 | –1.19077 | –0.00040 |
| O | 1.41197 | –2.27892 | –0.00072 |
| N | 1.37606 | 0.01773 | –0.00036 |
| C | 2.07154 | 1.16253 | –0.00008 |
| N | 1.38220 | 2.30680 | 0.00004 |
| C | 3.51513 | 1.16825 | 0.00004 |
| C | 4.13564 | –0.03782 | 0.00000 |
| C | –2.12085 | 1.25514 | 0.00018 |
| N | –1.48925 | 0.00207 | 0.00059 |
| C | –2.19534 | –1.15834 | 0.00049 |

| | | | |
|---|---|---|---|
| N | –3.53012 | –1.15836 | –0.00021 |
| C | –4.07719 | 0.07373 | –0.00086 |
| N | –3.49063 | 1.25616 | –0.00075 |
| N | –1.51604 | –2.31316 | 0.00125 |
| O | –1.42444 | 2.27784 | 0.00063 |
| H | 1.86876 | 3.18841 | 0.00061 |
| H | 4.07953 | 2.09153 | 0.00012 |
| H | 5.21433 | –0.14821 | 0.00009 |
| H | –5.16758 | 0.08523 | –0.00168 |
| H | –0.49253 | –2.34803 | 0.00108 |
| H | –2.06257 | –3.15900 | 0.00035 |
| H | –0.45072 | –0.01117 | 0.00050 |
| H | 0.34447 | 2.30005 | 0.00016 |
| H | 3.84631 | –2.09639 | –0.00030 |

**Table S59.** Optimized geometry coordinates of C:6GO (W–edge)
Energy: –612964.5480213 (au)

| | | | |
|---|---|---|---|
| N | 5.11204 | –0.52394 | 0.23892 |
| C | 3.85590 | –1.03706 | 0.67687 |
| O | 3.84678 | –2.03161 | 1.38380 |
| N | 2.74459 | –0.35488 | 0.25142 |
| C | 2.85643 | 0.70423 | –0.54120 |
| N | 1.71758 | 1.33367 | –0.90094 |
| C | 4.12533 | 1.20711 | –1.00948 |
| C | 5.23312 | 0.54868 | –0.58225 |
| C | –0.93713 | 2.83643 | 0.93807 |
| O | –2.17394 | 2.20068 | 0.59431 |
| C | –2.13917 | 0.89651 | 0.24567 |
| N | –0.99861 | 0.24174 | 0.08900 |
| C | –3.35152 | 0.23475 | 0.05600 |
| N | –4.69450 | 0.56564 | 0.12058 |
| C | –5.37183 | –0.58768 | –0.17280 |
| N | –4.59757 | –1.62274 | –0.41240 |
| C | –3.31060 | –1.13348 | –0.27231 |
| N | –2.17220 | –1.82847 | –0.39430 |
| C | –1.06266 | –1.10451 | –0.20175 |
| N | 0.13204 | –1.73577 | –0.33197 |
| H | 1.73924 | 2.05589 | –1.60147 |
| H | 4.19565 | 2.06899 | –1.65986 |
| H | 6.23984 | 0.83892 | –0.86482 |
| H | –0.31249 | 2.99180 | 0.05375 |
| H | –1.21654 | 3.80076 | 1.36456 |
| H | –0.37728 | 2.24267 | 1.66545 |
| H | –5.08657 | 1.46856 | 0.33528 |
| H | –6.45371 | –0.60932 | –0.19779 |
| H | 0.98197 | –1.32082 | 0.05418 |

| | | | |
|---|---|---|---|
| H | 0.08242 | –2.74040 | –0.39756 |
| H | 0.81477 | 0.97149 | –0.58304 |
| H | 5.91901 | –1.03776 | 0.56331 |

**Table S60.** Optimized geometry coordinates of C:6GU (W–edge)
Energy: –829500.2917551 (au)

| | | | |
|---|---|---|---|
| N | –5.11973 | –0.55143 | 0.01086 |
| C | –3.85653 | –1.20475 | –0.00098 |
| O | –3.82419 | –2.42711 | –0.00476 |
| N | –2.75548 | –0.39083 | –0.00728 |
| C | –2.87566 | 0.93524 | –0.00175 |
| N | –1.74511 | 1.66105 | –0.00844 |
| C | –4.15844 | 1.59747 | 0.01057 |
| C | –5.25640 | 0.79963 | 0.01646 |
| Cl | 1.90033 | 2.60873 | –0.00671 |
| N | 0.89749 | 0.17034 | –0.01012 |
| C | 0.96516 | –1.20354 | –0.00856 |
| N | –0.20381 | –1.86040 | –0.01469 |
| N | 2.10633 | –1.92994 | –0.00172 |
| C | 3.19590 | –1.17753 | 0.00346 |
| C | 3.26846 | 0.23597 | 0.00263 |
| C | 2.02184 | 0.85572 | –0.00463 |
| N | 4.58327 | 0.67226 | 0.00932 |
| C | 5.28267 | –0.42984 | 0.01402 |
| N | 4.50904 | –1.58247 | 0.01093 |
| H | –1.77651 | 2.66645 | –0.00251 |
| H | –4.24345 | 2.67626 | 0.01483 |
| H | –6.26921 | 1.18836 | 0.02568 |
| H | –1.10596 | –1.36781 | –0.01688 |
| H | 6.36319 | –0.48115 | 0.01990 |
| H | 4.83000 | –2.53880 | 0.01326 |
| H | –0.17902 | –2.86656 | –0.01173 |
| H | –0.83606 | 1.18793 | –0.01491 |
| H | –5.92107 | –1.16660 | 0.01520 |

**Table S61.** Optimized geometry coordinates of C:7DG (W–edge)
Energy: –889630.4085984 (au)

| | | | |
|---|---|---|---|
| O | –5.17477 | –1.49992 | 0.73985 |
| C | –4.76763 | –2.52377 | –0.16733 |
| C | –3.31773 | –2.30651 | –0.54837 |
| O | –3.18666 | –1.05608 | –1.25678 |
| C | –2.32227 | –2.22207 | 0.62474 |
| O | –1.88243 | –3.47526 | 1.07740 |
| C | –1.18404 | –1.38302 | –0.00197 |
| O | –0.36769 | –2.26933 | –0.73103 |
| C | –1.95319 | –0.44245 | –0.96098 |

| | | | |
|---|---|---|---|
| N | –2.21765 | 0.90224 | –0.36695 |
| C | –1.17871 | 1.85547 | –0.43092 |
| O | –0.09846 | 1.50730 | –0.94883 |
| N | –1.39171 | 3.09692 | 0.06199 |
| C | –2.56638 | 3.39736 | 0.60870 |
| N | –2.71853 | 4.66315 | 1.06915 |
| C | –3.63891 | 2.45961 | 0.70288 |
| C | –3.41749 | 1.21647 | 0.18951 |
| N | 6.37249 | 0.16275 | 0.42573 |
| C | 6.52457 | –1.08216 | 0.72544 |
| C | 5.27125 | –1.92623 | 0.65902 |
| C | 4.29720 | –0.87739 | 0.23628 |
| C | 2.90677 | –0.90807 | –0.02906 |
| O | 2.13526 | –1.88485 | 0.04909 |
| N | 2.41891 | 0.35363 | –0.41040 |
| C | 3.18407 | 1.49025 | –0.49490 |
| N | 2.54837 | 2.61705 | –0.87336 |
| N | 4.48693 | 1.51435 | –0.23784 |
| C | 4.99056 | 0.31972 | 0.11586 |
| H | –5.38894 | –2.51596 | –1.07438 |
| H | –4.83896 | –3.51915 | 0.29434 |
| H | –3.00870 | –3.12902 | –1.20825 |
| H | –2.78784 | –1.69430 | 1.46278 |
| H | –1.13422 | –3.68282 | 0.48610 |
| H | –0.60813 | –0.82218 | 0.74140 |
| H | 0.58308 | –2.09907 | –0.49919 |
| H | –1.37333 | –0.26631 | –1.86806 |
| H | –3.52633 | 4.93297 | 1.60387 |
| H | –4.59319 | 2.71611 | 1.14528 |
| H | –4.16136 | 0.42723 | 0.19259 |
| H | 7.50073 | –1.47400 | 1.00398 |
| H | 5.38255 | –2.75433 | –0.05508 |
| H | 1.53313 | 2.62608 | –0.94642 |
| H | 3.08545 | 3.46785 | –0.85289 |
| H | 1.42177 | 0.44026 | –0.61833 |
| H | 5.04587 | –2.38738 | 1.63091 |
| H | –1.91307 | 5.26895 | 1.05619 |
| H | –6.08061 | –1.68373 | 1.01601 |

**Table S62.** Optimized geometry coordinates of C:DGP (W–edge)

Energy: –1208769.4603988 (au)

| | | | |
|---|---|---|---|
| N | –7.03047 | 0.53853 | –0.98348 |
| C | –5.68741 | 0.88395 | –0.73054 |
| O | –5.27985 | 1.97651 | –1.13774 |
| N | –4.93119 | –0.01812 | –0.05121 |
| C | –5.44184 | –1.19027 | 0.34853 |

| | | | |
|---|---|---|---|
| N | –4.63593 | –2.02792 | 1.00698 |
| C | –6.81608 | –1.54867 | 0.08705 |
| C | –7.57116 | –0.64518 | –0.58519 |
| P | 3.01614 | –2.50384 | –0.60087 |
| O | 4.19082 | –3.60381 | –0.77974 |
| O | 2.46757 | –2.84233 | 0.84734 |
| O | 2.07787 | –2.43784 | –1.74513 |
| O | 3.86143 | –1.14317 | –0.32924 |
| C | 4.34023 | –0.35533 | –1.43303 |
| C | 4.33203 | 1.12521 | –1.06470 |
| O | 2.99724 | 1.64769 | –1.06550 |
| C | 4.96134 | 1.46999 | 0.31158 |
| O | 6.30759 | 1.89898 | 0.23463 |
| C | 3.98169 | 2.51108 | 0.89426 |
| C | 2.65271 | 2.23363 | 0.18531 |
| N | 1.74633 | 1.35563 | 0.92567 |
| C | 2.07689 | 0.18334 | 1.57703 |
| N | 1.06024 | –0.62330 | 1.75301 |
| C | –0.02013 | 0.04506 | 1.19156 |
| C | –1.39558 | –0.32706 | 1.06510 |
| O | –1.95107 | –1.36375 | 1.45523 |
| N | –2.13035 | 0.67102 | 0.39977 |
| C | –1.62277 | 1.85306 | –0.08636 |
| N | –2.49458 | 2.68555 | –0.68852 |
| N | –0.34552 | 2.19994 | 0.02723 |
| C | 0.38695 | 1.26778 | 0.66194 |
| H | –3.64449 | –1.78103 | 1.17880 |
| H | –7.22660 | –2.49585 | 0.41160 |
| H | –8.61326 | –0.81185 | –0.83430 |
| H | 5.36774 | –0.66201 | –1.66230 |
| H | 3.70514 | –0.51629 | –2.30878 |
| H | 4.89860 | 1.64718 | –1.85224 |
| H | 4.99683 | 0.57395 | 0.93445 |
| H | 6.33790 | 2.70342 | –0.30257 |
| H | 3.90732 | 2.48395 | 1.98303 |
| H | 4.32369 | 3.51297 | 0.61023 |
| H | 2.07030 | 3.13249 | –0.02320 |
| H | 3.08822 | –0.04115 | 1.87437 |
| H | –2.10779 | 3.51859 | –1.09983 |
| H | –3.13278 | 0.45924 | 0.24865 |
| H | –3.47266 | 2.43650 | –0.85770 |
| H | 1.80208 | –2.19524 | 1.21017 |
| H | 3.90246 | –4.27493 | –1.41418 |
| H | –4.97874 | –2.92572 | 1.30732 |
| H | –7.56989 | 1.22609 | –1.49058 |

**Table S63.** Optimized geometry coordinates of C:H4B (W–edge)
Energy: –782140.3734039 (au)

| | | | |
|---|---|---|---|
| N | 5.26117 | –1.89821 | 0.03411 |
| C | 4.19074 | –1.03641 | 0.36995 |
| O | 3.15487 | –1.53315 | 0.80000 |
| N | 4.39631 | 0.30476 | 0.18572 |
| C | 5.54766 | 0.74481 | –0.30289 |
| N | 5.68539 | 2.08381 | –0.45450 |
| C | 6.64124 | –0.12208 | –0.65297 |
| C | 6.44176 | –1.45272 | –0.46259 |
| N | –0.40210 | 1.98718 | 0.67227 |
| C | 0.61635 | 1.16557 | 0.77660 |
| N | 1.75411 | 1.56774 | 1.43973 |
| N | 0.59975 | –0.11168 | 0.29724 |
| C | –0.50623 | –0.66615 | –0.37951 |
| O | –0.49736 | –1.84800 | –0.75468 |
| C | –1.59011 | 0.24863 | –0.54370 |
| C | –1.51111 | 1.50215 | 0.03500 |
| N | –2.72457 | –0.14966 | –1.27704 |
| N | –2.62238 | 2.32348 | 0.01823 |
| C | –3.97342 | 0.50580 | –0.93522 |
| C | –3.71105 | 2.02545 | –0.90773 |
| C | –4.63703 | 0.03317 | 0.38159 |
| O | –5.92238 | 0.67001 | 0.38934 |
| C | –4.78478 | –1.50240 | 0.50632 |
| C | –3.61061 | –2.20207 | 1.19711 |
| O | –5.01588 | –2.03659 | –0.80083 |
| H | 6.47097 | 2.47411 | –0.94723 |
| H | 7.57472 | 0.25959 | –1.04473 |
| H | 7.19045 | –2.20294 | –0.69191 |
| H | 2.63702 | 1.17335 | 1.12318 |
| H | –4.69764 | 0.30217 | –1.72942 |
| H | –3.48070 | 2.34969 | –1.93322 |
| H | –4.61305 | 2.54215 | –0.57332 |
| H | –4.03840 | 0.38808 | 1.23291 |
| H | –6.30049 | 0.56572 | 1.27146 |
| H | –3.42786 | –1.77118 | 2.18740 |
| H | –3.84404 | –3.26424 | 1.34474 |
| H | –2.68830 | –2.13966 | 0.61505 |
| H | 5.07707 | –2.88364 | 0.16691 |
| H | 4.87097 | 2.66098 | –0.31104 |
| H | 1.39829 | –0.72796 | 0.46447 |
| H | –2.43307 | 3.29066 | 0.23927 |
| H | –2.78475 | –1.15107 | –1.41077 |
| H | 1.75944 | 2.56164 | 1.61751 |
| H | –5.05417 | –2.99837 | –0.72131 |

| | | | |
|---|---|---|---|
| H | –5.68465 | –1.66172 | 1.12739 |

**Table S64.** Optimized geometry coordinates of C:PQ0 (W–edge)
Energy: –636142.8278525 (au)

| | | | |
|---|---|---|---|
| N | –4.80581 | –0.49579 | –0.00501 |
| C | –3.44367 | –0.85820 | –0.00167 |
| O | –3.15966 | –2.06039 | –0.00225 |
| N | –2.53260 | 0.15039 | 0.00207 |
| C | –2.91742 | 1.43379 | 0.00170 |
| N | –1.96415 | 2.36961 | 0.00506 |
| C | –4.31214 | 1.80766 | –0.00184 |
| C | –5.22005 | 0.80062 | –0.00511 |
| N | 0.28414 | –0.60554 | 0.00516 |
| C | 0.64931 | –1.93216 | 0.00359 |
| N | 1.91109 | –2.33951 | 0.00037 |
| C | 2.79474 | –1.32099 | –0.00084 |
| C | 2.53887 | 0.05614 | 0.00056 |
| C | 1.17311 | 0.47889 | 0.00315 |
| O | 0.74433 | 1.64330 | 0.00379 |
| C | 3.81278 | 0.73008 | –0.00186 |
| C | 4.06831 | 2.12671 | –0.00173 |
| N | 4.30452 | 3.26678 | –0.00179 |
| C | 4.78056 | –0.25410 | –0.00492 |
| N | 4.15789 | –1.48493 | –0.00395 |
| N | –0.35343 | –2.83232 | 0.00500 |
| H | –0.96275 | 2.10446 | 0.00517 |
| H | –4.62117 | 2.84474 | –0.00197 |
| H | –6.29131 | 0.96849 | –0.00790 |
| H | 5.85542 | –0.16307 | –0.00674 |
| H | 4.61437 | –2.38383 | –0.00582 |
| H | –0.09100 | –3.80359 | 0.00229 |
| H | –2.21420 | 3.34499 | 0.00354 |
| H | –5.46208 | –1.26404 | –0.00753 |
| H | –1.34018 | –2.56186 | 0.00656 |
| H | –0.72236 | –0.36104 | 0.00547 |

**Table S65.** Optimized geometry coordinates of 1C:PRF (W–edge)
Energy: –637665.0915774 (au)

| | | | |
|---|---|---|---|
| N | –4.87752 | –0.48951 | –0.00652 |
| C | –3.52080 | –0.87618 | 0.01454 |
| O | –3.25844 | –2.08177 | 0.05004 |
| N | –2.59289 | 0.11697 | –0.00641 |
| C | –2.95543 | 1.40500 | –0.05114 |
| N | –1.98464 | 2.32333 | –0.07499 |
| C | –4.34336 | 1.80382 | –0.07215 |

| | | | |
|---|---|---|---|
| C | –5.26888 | 0.81303 | –0.04839 |
| N | 0.22831 | –0.67335 | 0.02065 |
| C | 0.57648 | –2.00402 | 0.08854 |
| N | 1.82835 | –2.42891 | 0.06799 |
| C | 2.73073 | –1.42689 | –0.03105 |
| C | 2.49201 | –0.04188 | –0.11219 |
| C | 1.13555 | 0.38760 | –0.07265 |
| O | 0.70770 | 1.56452 | –0.10609 |
| C | 3.76764 | 0.62666 | –0.20266 |
| C | 4.01003 | 2.11035 | –0.32260 |
| N | 3.38810 | 2.95276 | 0.70884 |
| C | 4.71225 | –0.36595 | –0.16557 |
| N | 4.08161 | –1.60750 | –0.06761 |
| N | –0.44026 | –2.88809 | 0.20000 |
| H | –0.98733 | 2.03764 | –0.07933 |
| H | –4.63416 | 2.84562 | –0.10647 |
| H | –6.33704 | 0.99956 | –0.06167 |
| H | 5.09161 | 2.29404 | –0.32606 |
| H | 3.63534 | 2.46490 | –1.29101 |
| H | 2.37876 | 2.81724 | 0.63804 |
| H | 3.65263 | 2.59268 | 1.62459 |
| H | 5.78934 | –0.29909 | –0.21165 |
| H | 4.52994 | –2.50793 | –0.01200 |
| H | –0.18820 | –3.86192 | 0.17853 |
| H | –5.54702 | –1.24602 | 0.00990 |
| H | –2.21844 | 3.30165 | –0.11959 |
| H | –0.77238 | –0.41535 | 0.03342 |
| H | –1.42184 | –2.61406 | 0.13010 |

**Table S66.** Optimized geometry coordinates of 2C:PRF (W–edge)
Energy: –637650.3755579 (au)

| | | | |
|---|---|---|---|
| N | –4.19055 | –1.79951 | 0.32415 |
| C | –3.11268 | –0.89405 | 0.45475 |
| O | –2.07079 | –1.29705 | 0.96201 |
| N | –3.31599 | 0.38106 | –0.00060 |
| C | –4.47606 | 0.71888 | –0.54802 |
| N | –4.61306 | 1.99724 | –0.97240 |
| C | –5.57830 | –0.19457 | –0.69055 |
| C | –5.37941 | –1.45815 | –0.23186 |
| N | 0.54530 | –0.03831 | 0.41963 |
| C | 0.50537 | 1.31713 | 0.62890 |
| N | 1.54689 | 2.11206 | 0.49744 |
| C | 2.67771 | 1.45271 | 0.13453 |
| C | 2.83849 | 0.08477 | –0.12871 |
| C | 1.69601 | –0.76940 | 0.02998 |
| O | 1.61721 | –1.99124 | –0.11717 |

| | | | |
|---|---|---|---|
| C | 4.21535 | –0.13992 | –0.49009 |
| C | 4.85180 | –1.45286 | –0.87452 |
| N | 4.65884 | –2.56676 | 0.06319 |
| C | 4.82626 | 1.08742 | –0.42345 |
| N | 3.88976 | 2.05132 | –0.05116 |
| N | –0.69412 | 1.83892 | 1.03602 |
| H | –5.41430 | 2.28765 | –1.50692 |
| H | –6.51823 | 0.10413 | –1.13527 |
| H | –6.13496 | –2.23409 | –0.28588 |
| H | 4.45374 | –1.78134 | –1.84311 |
| H | 5.92755 | –1.29423 | –1.02398 |
| H | 4.93713 | –2.25917 | 0.99394 |
| H | 3.65338 | –2.74077 | 0.11864 |
| H | 5.85193 | 1.36357 | –0.61989 |
| H | 4.06043 | 3.03395 | 0.08950 |
| H | –1.55064 | 1.36897 | 0.75263 |
| H | –0.70893 | 2.84766 | 1.02367 |
| H | –3.79929 | 2.59233 | –0.95778 |
| H | –0.29799 | –0.59000 | 0.59741 |
| H | –4.00804 | –2.73633 | 0.65864 |

**Table S67.** Optimized geometry coordinates of C:THF (W–edge)
Energy: –650888.7116509 (au)

| | | | |
|---|---|---|---|
| N | –4.11136 | –1.72374 | –0.08635 |
| C | –2.75956 | –1.39354 | –0.05663 |
| O | –1.89485 | –2.26125 | –0.09860 |
| N | –2.50724 | –0.04366 | 0.02082 |
| C | –3.46163 | 0.97758 | 0.06294 |
| O | –3.11953 | 2.15815 | 0.12874 |
| C | –4.84705 | 0.53005 | 0.02297 |
| C | –5.11807 | –0.79055 | –0.04855 |
| N | 0.29387 | 0.76222 | 0.05198 |
| C | 0.61434 | 2.03831 | 0.12589 |
| N | –0.33671 | 2.98204 | 0.39337 |
| N | 1.90301 | 2.46411 | –0.00667 |
| C | 3.01299 | 1.60469 | –0.22083 |
| O | 4.15239 | 2.07594 | –0.28294 |
| C | 2.64363 | 0.23475 | –0.32699 |
| N | 3.64607 | –0.72377 | –0.59908 |
| C | 3.37778 | –2.06043 | –0.06688 |
| C | 1.96995 | –2.45866 | –0.55590 |
| N | 0.99009 | –1.46484 | –0.14445 |
| C | 1.32016 | –0.13893 | –0.15048 |
| C | 3.51713 | –2.17398 | 1.46007 |
| H | –5.62763 | 1.27673 | 0.05387 |
| H | –6.12662 | –1.18715 | –0.07980 |

| | | | |
|---|---|---|---|
| H | –0.15195 | 3.92333 | 0.08023 |
| H | 4.09174 | –2.74774 | –0.53800 |
| H | 1.99756 | –2.57005 | –1.64930 |
| H | 1.68280 | –3.42412 | –0.12812 |
| H | 3.34873 | –3.20370 | 1.79580 |
| H | 4.52340 | –1.87841 | 1.77503 |
| H | 2.79752 | –1.52697 | 1.96894 |
| H | 4.56074 | –0.34973 | –0.36546 |
| H | 0.00886 | –1.72556 | –0.11801 |
| H | –1.31186 | 2.68674 | 0.31165 |
| H | 2.14614 | 3.43652 | 0.13136 |
| H | –1.49789 | 0.23770 | 0.04650 |
| H | –4.30771 | –2.71262 | –0.14328 |

**Table S68.** Optimized geometry coordinates of C:XAN (W–edge)
Energy: –600778.5492517 (au)

| | | | |
|---|---|---|---|
| N | –4.40745 | 0.88792 | 0.23250 |
| C | –3.00099 | 1.04781 | 0.40036 |
| O | –2.57648 | 2.13266 | 0.76085 |
| N | –2.23173 | –0.05975 | 0.13917 |
| C | –2.79375 | –1.22068 | –0.19501 |
| N | –1.98051 | –2.27207 | –0.40191 |
| C | –4.22001 | –1.38111 | –0.35241 |
| C | –4.98691 | –0.28425 | –0.12837 |
| N | 4.71916 | 0.37478 | –0.01924 |
| C | 3.36558 | 0.53580 | –0.07953 |
| N | 2.63043 | 1.66273 | –0.31063 |
| C | 1.22291 | 1.59936 | –0.34600 |
| O | 0.56383 | 2.58867 | –0.59140 |
| N | 0.69672 | 0.33937 | –0.08765 |
| C | 1.37614 | –0.87110 | 0.12590 |
| O | 0.76682 | –1.92924 | 0.27690 |
| C | 2.81663 | –0.71340 | 0.13673 |
| N | 3.81650 | –1.65019 | 0.32888 |
| C | 4.93051 | –0.98150 | 0.23386 |
| H | –2.37075 | –3.18291 | –0.57639 |
| H | –4.65752 | –2.32944 | –0.63589 |
| H | –6.06815 | –0.28942 | –0.21905 |
| H | 5.42602 | 1.08612 | –0.12911 |
| H | 5.92945 | –1.38229 | 0.33133 |
| H | –0.34078 | 0.29799 | –0.02601 |
| H | –0.98522 | –2.18888 | –0.16603 |
| H | –4.95370 | 1.71688 | 0.41916 |
| H | 3.02476 | 2.57008 | –0.50983 |

**Table S69.** Optimized geometry coordinates of A:FMN (W–edge)

Energy: -1484281.9377768 (au)

| | | | |
|---|---|---|---|
| N | 8.53035 | 0.25639 | 0.12226 |
| C | 8.96892 | -1.00444 | -0.23031 |
| N | 7.99501 | -1.86598 | -0.39634 |
| C | 6.85144 | -1.12980 | -0.13880 |
| C | 5.47992 | -1.47342 | -0.14815 |
| N | 5.03653 | -2.70777 | -0.43956 |
| N | 4.59209 | -0.49906 | 0.15590 |
| C | 5.03498 | 0.74072 | 0.44982 |
| N | 6.29263 | 1.17547 | 0.49017 |
| C | 7.15655 | 0.19499 | 0.18636 |
| N | -0.37156 | 0.01465 | 0.70323 |
| C | 1.00821 | 0.18602 | 0.60159 |
| O | 1.52495 | 1.27516 | 0.79718 |
| N | 1.79218 | -0.92227 | 0.26425 |
| C | 1.33641 | -2.16962 | -0.07461 |
| O | 2.07921 | -3.09041 | -0.39899 |
| C | -0.15278 | -2.31808 | -0.00829 |
| N | -0.69591 | -3.45613 | -0.33153 |
| C | -2.05602 | -3.56228 | -0.25537 |
| C | -2.65678 | -4.77555 | -0.65532 |
| C | -4.02366 | -4.96834 | -0.60061 |
| C | -4.63548 | -6.27625 | -1.03595 |
| C | -4.84156 | -3.89574 | -0.13662 |
| C | -6.34029 | -4.04864 | -0.09455 |
| C | -4.26751 | -2.69456 | 0.26779 |
| C | -2.87695 | -2.50516 | 0.22672 |
| N | -2.25452 | -1.32933 | 0.64959 |
| C | -0.90324 | -1.16139 | 0.44779 |
| C | -2.97683 | -0.34886 | 1.49954 |
| C | -3.71263 | 0.82679 | 0.79890 |
| O | -4.42613 | 0.36779 | -0.33794 |
| C | -2.87774 | 2.05940 | 0.41133 |
| O | -1.99020 | 1.73143 | -0.66475 |
| C | -2.18204 | 2.76631 | 1.59598 |
| O | -3.18788 | 3.26159 | 2.47646 |
| C | -1.23047 | 3.87363 | 1.15520 |
| O | -1.97402 | 4.83736 | 0.37207 |
| P | -1.46983 | 5.17557 | -1.14157 |
| O | -2.42006 | 6.42011 | -1.51027 |
| O | -2.11405 | 4.03925 | -2.05837 |
| O | -0.01511 | 5.42267 | -1.24804 |
| H | 10.02167 | -1.22272 | -0.34874 |
| H | 5.70755 | -3.41923 | -0.67897 |
| H | 4.25196 | 1.45976 | 0.67850 |
| H | -1.99030 | -5.55471 | -1.01082 |

| | | | |
|---|---|---|---|
| H | -3.86451 | -6.98231 | -1.35212 |
| H | -5.20747 | -6.74347 | -0.22598 |
| H | -5.32809 | -6.13825 | -1.87442 |
| H | -6.63672 | -4.87386 | 0.56362 |
| H | -6.82400 | -3.13816 | 0.26469 |
| H | -6.74479 | -4.27803 | -1.08714 |
| H | -4.91259 | -1.87242 | 0.54570 |
| H | -3.71663 | -0.93086 | 2.05005 |
| H | -2.24764 | 0.03168 | 2.21588 |
| H | -4.45137 | 1.17553 | 1.52845 |
| H | -3.80587 | 0.43661 | -1.08178 |
| H | -3.59090 | 2.77213 | -0.01624 |
| H | -1.24110 | 1.22587 | -0.25870 |
| H | -1.58623 | 2.05461 | 2.17608 |
| H | -3.54801 | 4.05826 | 2.05733 |
| H | -0.85588 | 4.39106 | 2.04272 |
| H | -0.37212 | 3.47750 | 0.60710 |
| H | 4.03836 | -2.91156 | -0.45340 |
| H | 2.83109 | -0.75958 | 0.21898 |
| H | 9.09197 | 1.07578 | 0.29851 |
| H | -1.93445 | 3.13343 | -1.70475 |
| H | -1.88259 | 7.22364 | -1.55017 |

**Table S70.** Optimized geometry coordinates of U:A2F (W–edge)
Energy: -615843.2608912 (au)

| | | | |
|---|---|---|---|
| N | -4.46258 | 0.82617 | 0.07149 |
| C | -3.09382 | 1.11656 | 0.13689 |
| O | -2.68482 | 2.25232 | 0.26884 |
| N | -2.28879 | -0.00844 | 0.03793 |
| C | -2.71187 | -1.32901 | -0.10331 |
| O | -1.89978 | -2.25684 | -0.16901 |
| C | -4.15370 | -1.51458 | -0.16402 |
| C | -4.96673 | -0.43885 | -0.07461 |
| F | 0.39662 | 2.42592 | -0.15238 |
| C | 1.27695 | 1.42592 | -0.08693 |
| N | 2.54040 | 1.76518 | -0.14839 |
| N | 0.70005 | 0.22207 | 0.02468 |
| C | 1.50157 | -0.86031 | 0.09430 |
| N | 0.92634 | -2.07793 | 0.22800 |
| C | 2.89017 | -0.62485 | 0.05897 |
| C | 3.35818 | 0.69383 | -0.07065 |
| N | 4.03236 | -1.41049 | 0.10436 |
| C | 5.08943 | -0.54180 | 0.01155 |
| N | 4.73678 | 0.71750 | -0.10028 |
| H | -4.53700 | -2.51850 | -0.27679 |
| H | -6.04809 | -0.51334 | -0.11112 |

| | | | |
|---|---|---|---|
| H | -0.08840 | -2.15035 | 0.09013 |
| H | 4.09690 | -2.40548 | 0.25034 |
| H | 6.11067 | -0.89966 | 0.03105 |
| H | -5.06663 | 1.63233 | 0.14036 |
| H | -1.26799 | 0.16132 | 0.06027 |
| H | 1.47479 | -2.90322 | 0.05092 |

**Table S71.** Optimized geometry coordinates of 1U:ADE (W–edge)
Energy: -553579.8194580 (au)

| | | | |
|---|---|---|---|
| N | -4.25801 | -0.98464 | 0.00054 |
| C | -2.89581 | -1.30401 | 0.00002 |
| O | -2.50441 | -2.45703 | -0.00005 |
| N | -2.06686 | -0.19700 | -0.00037 |
| C | -2.45770 | 1.13930 | -0.00025 |
| O | -1.62682 | 2.05400 | -0.00066 |
| C | -3.89662 | 1.36173 | 0.00031 |
| C | -4.73375 | 0.30087 | 0.00068 |
| N | 1.19182 | 1.72260 | -0.00018 |
| C | 2.36857 | 2.43706 | 0.00017 |
| N | 3.44890 | 1.68683 | 0.00038 |
| C | 2.95004 | 0.39610 | 0.00007 |
| C | 3.56295 | -0.87480 | 0.00024 |
| N | 4.90539 | -1.02785 | 0.00103 |
| N | 2.78676 | -1.97760 | -0.00016 |
| C | 1.45661 | -1.82353 | -0.00063 |
| N | 0.75732 | -0.68254 | -0.00070 |
| C | 1.55220 | 0.40187 | -0.00032 |
| H | -4.25885 | 2.37983 | 0.00043 |
| H | -5.81345 | 0.40265 | 0.00112 |
| H | 0.21731 | 2.04678 | -0.00079 |
| H | 2.36838 | 3.51871 | 0.00035 |
| H | 5.50580 | -0.22046 | 0.00013 |
| H | 0.86697 | -2.73704 | -0.00102 |
| H | -4.88048 | -1.77959 | 0.00081 |
| H | -1.04345 | -0.41529 | -0.00066 |
| H | 5.29157 | -1.95735 | -0.00023 |

**Table S72.** Optimized geometry coordinates of 2U:ADE (W–edge)
Energy: -553578.6066158 (au)

| | | | |
|---|---|---|---|
| N | 4.27424 | -1.07711 | -0.00225 |
| C | 2.88630 | -1.24861 | -0.00028 |
| O | 2.37514 | -2.35649 | 0.00033 |
| N | 2.18210 | -0.06293 | 0.00069 |
| C | 2.70961 | 1.22671 | 0.00133 |
| O | 1.96988 | 2.21402 | 0.00316 |
| C | 4.16513 | 1.29630 | -0.00050 |

| | | | |
|---|---|---|---|
| C | 4.88380 | 0.15215 | -0.00192 |
| N | -4.62933 | -0.80663 | -0.00029 |
| C | -4.99301 | 0.52542 | -0.00136 |
| N | -3.96883 | 1.34340 | -0.00156 |
| C | -2.86962 | 0.50201 | -0.00057 |
| C | -1.47826 | 0.75865 | -0.00010 |
| N | -0.95908 | 1.99612 | -0.00084 |
| N | -0.64955 | -0.31165 | 0.00098 |
| C | -1.16579 | -1.55641 | 0.00155 |
| N | -2.44816 | -1.91620 | 0.00126 |
| C | -3.25286 | -0.84211 | 0.00021 |
| H | 4.63462 | 2.26970 | -0.00047 |
| H | 5.96821 | 0.13783 | -0.00318 |
| H | -5.23919 | -1.61023 | 0.00010 |
| H | -6.03191 | 0.82654 | -0.00197 |
| H | -1.58662 | 2.78360 | -0.00085 |
| H | -0.42629 | -2.35405 | 0.00229 |
| H | 0.05327 | 2.13580 | 0.00049 |
| H | 1.13673 | -0.15614 | 0.00127 |
| H | 4.80951 | -1.93303 | -0.00177 |

**Table S73.** Optimized geometry coordinates of 2U:H4B (W–edge)
Energy: -794619.4141100 (au)

| | | | |
|---|---|---|---|
| N | 5.52080 | -2.05567 | 0.03004 |
| C | 4.16161 | -2.00642 | -0.31286 |
| O | 3.57458 | -2.99057 | -0.71815 |
| N | 3.59003 | -0.75644 | -0.14050 |
| C | 4.23052 | 0.38314 | 0.33200 |
| O | 3.62477 | 1.45657 | 0.45856 |
| C | 5.63855 | 0.22522 | 0.66111 |
| C | 6.22674 | -0.98038 | 0.49720 |
| N | -1.10374 | 2.94980 | 0.03052 |
| C | 0.19458 | 2.83141 | 0.16939 |
| N | 0.93789 | 3.90545 | 0.60355 |
| N | 0.88421 | 1.69947 | -0.13905 |
| C | 0.25774 | 0.53959 | -0.60782 |
| O | 0.92432 | -0.50902 | -0.79902 |
| C | -1.13227 | 0.67225 | -0.82434 |
| C | -1.75895 | 1.86622 | -0.48331 |
| N | -1.88996 | -0.40271 | -1.33945 |
| N | -3.10805 | 1.98776 | -0.64812 |
| C | -3.23203 | -0.50801 | -0.76731 |
| C | -3.94524 | 0.84209 | -1.00075 |
| C | -3.27736 | -0.97662 | 0.72045 |
| O | -3.45610 | 0.15865 | 1.56456 |
| C | -4.37788 | -2.02443 | 0.98057 |

| | | | |
|---|---|---|---|
| C | -4.05020 | -3.40255 | 0.40145 |
| O | -5.59703 | -1.50937 | 0.44879 |
| H | 6.17957 | 1.08542 | 1.02842 |
| H | 7.27197 | -1.16263 | 0.72230 |
| H | 1.82831 | 3.70724 | 1.03782 |
| H | 0.38885 | 4.63093 | 1.03994 |
| H | -1.34860 | -1.26122 | -1.30326 |
| H | -3.52606 | 2.73342 | -0.11184 |
| H | -3.77776 | -1.24691 | -1.35997 |
| H | -4.86287 | 0.87005 | -0.41156 |
| H | -4.21422 | 0.89640 | -2.06228 |
| H | -2.31172 | -1.45852 | 0.94359 |
| H | -3.37858 | -0.13729 | 2.48087 |
| H | -3.93856 | -3.36290 | -0.68611 |
| H | -3.12588 | -3.80404 | 0.82966 |
| H | -4.85329 | -4.11352 | 0.62795 |
| H | 5.95243 | -2.95981 | -0.09441 |
| H | 2.57293 | -0.67909 | -0.39853 |
| H | 1.89959 | 1.63844 | 0.02421 |
| H | -4.46212 | -2.12631 | 2.07864 |
| H | -6.29148 | -2.16246 | 0.60324 |

**Table S74.** Optimized geometry coordinates of 1U:SAM (W–edge)
Energy: -1319605.0950895 (au)

| | | | |
|---|---|---|---|
| N | 6.86814 | -0.70262 | -1.52328 |
| C | 5.51711 | -0.89694 | -1.24858 |
| O | 4.86624 | -1.81415 | -1.72720 |
| N | 4.99402 | 0.04656 | -0.38930 |
| C | 5.65252 | 1.12075 | 0.20352 |
| O | 5.04218 | 1.88481 | 0.95749 |
| C | 7.05804 | 1.23473 | -0.14703 |
| C | 7.60487 | 0.32623 | -0.98736 |
| N | -0.99647 | 3.14553 | -0.53623 |
| C | -2.39556 | 3.20116 | -0.94351 |
| C | -3.05477 | 4.47237 | -0.33066 |
| O | -4.25307 | 4.58791 | -0.21608 |
| O | -2.17929 | 5.41485 | 0.00756 |
| C | -3.30393 | 1.98642 | -0.63436 |
| C | -3.10643 | 0.84694 | -1.62556 |
| S | -4.28114 | -0.58513 | -1.47210 |
| C | -5.88333 | 0.17804 | -1.05340 |
| C | -3.79729 | -1.41340 | 0.10919 |
| C | -2.89178 | -2.60210 | -0.16914 |
| O | -1.74880 | -2.15351 | -0.91587 |
| C | -2.35434 | -3.24436 | 1.13778 |
| O | -3.17744 | -4.26234 | 1.62896 |

| | | | |
|---|---|---|---|
| C | -0.95509 | -3.72432 | 0.68919 |
| O | -1.03118 | -5.04573 | 0.16862 |
| C | -0.53620 | -2.70107 | -0.38868 |
| N | 0.34077 | -1.63305 | 0.05786 |
| C | 1.65926 | -1.48015 | -0.34018 |
| N | 2.23626 | -0.43988 | 0.20055 |
| C | 1.26989 | 0.14227 | 0.99589 |
| C | 1.28504 | 1.30092 | 1.81814 |
| N | 2.35114 | 2.08559 | 1.97595 |
| N | 0.12980 | 1.62081 | 2.45986 |
| C | -0.93279 | 0.82672 | 2.31824 |
| N | -1.06889 | -0.28064 | 1.56817 |
| C | 0.07633 | -0.57783 | 0.92657 |
| H | 7.63656 | 2.04392 | 0.27526 |
| H | 8.64704 | 0.35679 | -1.28445 |
| H | -0.39614 | 2.65432 | -1.18824 |
| H | -0.86241 | 2.76280 | 0.39712 |
| H | -2.40777 | 3.38405 | -2.02708 |
| H | -3.11031 | 1.64829 | 0.38885 |
| H | -4.32585 | 2.37132 | -0.67560 |
| H | -2.13275 | 0.35527 | -1.54937 |
| H | -3.25085 | 1.17353 | -2.66063 |
| H | -6.60971 | -0.63470 | -0.99952 |
| H | -5.83032 | 0.71719 | -0.10790 |
| H | -3.27719 | -0.68458 | 0.73657 |
| H | -4.72741 | -1.74397 | 0.57949 |
| H | -3.43086 | -3.36317 | -0.75060 |
| H | -2.24446 | -2.46958 | 1.90168 |
| H | -2.81256 | -5.10138 | 1.29917 |
| H | -0.24072 | -3.76826 | 1.51231 |
| H | -1.43963 | -5.03814 | -0.70948 |
| H | 0.00648 | -3.21570 | -1.18858 |
| H | 2.14751 | -2.16275 | -1.02466 |
| H | 3.26742 | 1.89856 | 1.55187 |
| H | -1.81372 | 1.12488 | 2.88420 |
| H | 3.99622 | -0.08055 | -0.15539 |
| H | 2.26207 | 2.86364 | 2.61360 |
| H | -1.28826 | 5.02685 | -0.15266 |
| H | -6.15266 | 0.85230 | -1.86836 |
| H | 7.28663 | -1.37807 | -2.14727 |

**Table S75.** Optimized geometry coordinates of 3U:SAM (W–edge)
Energy: -1352537.2166018 (au)

| | | | |
|---|---|---|---|
| N | -4.24404 | 3.94428 | -1.80477 |
| C | -5.02050 | 4.54390 | -0.83149 |
| N | -5.55597 | 3.68775 | 0.00455 |

| | | | |
|---|---|---|---|
| C | -5.11660 | 2.46033 | -0.44765 |
| C | -5.34174 | 1.14819 | 0.02884 |
| N | -6.08901 | 0.90736 | 1.11849 |
| N | -4.76551 | 0.12333 | -0.64810 |
| C | -4.01780 | 0.40980 | -1.73326 |
| N | -3.70940 | 1.59235 | -2.25695 |
| C | -4.29793 | 2.59149 | -1.57347 |
| N | -4.76434 | -2.79937 | 0.23830 |
| C | -3.91592 | -2.73119 | 1.42285 |
| C | -4.74562 | -2.13062 | 2.58952 |
| O | -4.24771 | -1.71306 | 3.60537 |
| O | -6.06372 | -2.09911 | 2.33157 |
| C | -2.61821 | -1.89202 | 1.31477 |
| C | -1.69707 | -2.40191 | 0.21001 |
| S | -0.09547 | -1.48129 | 0.19624 |
| C | -0.56929 | 0.22572 | -0.23494 |
| C | 0.69243 | -2.14042 | -1.34565 |
| C | 1.74323 | -1.16845 | -1.88708 |
| O | 2.43109 | -0.60515 | -0.77354 |
| C | 2.81168 | -1.84180 | -2.79675 |
| O | 2.98361 | -1.18625 | -4.04072 |
| C | 4.09825 | -1.76000 | -1.93369 |
| O | 5.27704 | -1.76013 | -2.67495 |
| C | 3.84169 | -0.47642 | -1.11802 |
| N | 4.60824 | -0.27907 | 0.05965 |
| C | 5.08251 | -1.19440 | 1.00135 |
| N | 5.70717 | -0.63437 | 1.99990 |
| C | 5.65039 | 0.72554 | 1.72926 |
| C | 6.14175 | 1.86572 | 2.40615 |
| N | 6.81120 | 1.78898 | 3.57287 |
| N | 5.91642 | 3.07921 | 1.86328 |
| C | 5.24138 | 3.15557 | 0.71027 |
| N | 4.72543 | 2.15903 | -0.02593 |
| C | 4.97186 | 0.97532 | 0.53949 |
| H | -5.14999 | 5.61695 | -0.79438 |
| H | -6.25026 | -0.02993 | 1.46186 |
| H | -3.61390 | -0.45861 | -2.25440 |
| H | -6.14945 | -2.52994 | 1.44120 |
| H | -4.54422 | -3.55928 | -0.39366 |
| H | -3.65850 | -3.75492 | 1.72354 |
| H | -2.90677 | -0.85792 | 1.10706 |
| H | -2.12457 | -1.91440 | 2.29026 |
| H | -2.15071 | -2.30982 | -0.78077 |
| H | -1.39840 | -3.44333 | 0.37244 |
| H | -1.18811 | 0.26416 | -1.13302 |
| H | -1.11370 | 0.62287 | 0.62256 |

| | | | |
|---|---|---|---|
| H | 0.36863 | 0.76926 | -0.35531 |
| H | -0.08957 | -2.34570 | -2.08145 |
| H | 1.14579 | -3.08051 | -1.01876 |
| H | 1.23840 | -0.38214 | -2.46819 |
| H | 2.55981 | -2.87326 | -3.05526 |
| H | 3.05530 | -0.22904 | -3.90936 |
| H | 4.11886 | -2.61013 | -1.24070 |
| H | 5.05054 | -1.49562 | -3.58183 |
| H | 3.98320 | 0.42350 | -1.73040 |
| H | 4.94516 | -2.25934 | 0.87848 |
| H | 7.19962 | 2.63274 | 3.96405 |
| H | 5.09311 | 4.16031 | 0.32145 |
| H | -6.50749 | 1.68575 | 1.60455 |
| H | -3.76567 | 4.40014 | -2.56833 |
| H | -4.81898 | -1.89699 | -0.24854 |
| H | 7.06857 | 0.89274 | 3.95318 |

**Table S76.** Optimized geometry coordinates of C:GMP (W–edge)
Energy: -899691.7051697 (au)

| | | | |
|---|---|---|---|
| N | -5.62994 | -1.86314 | 0.18313 |
| C | -5.08819 | -0.57010 | -0.05174 |
| O | -5.85008 | 0.37485 | -0.19904 |
| N | -3.72198 | -0.48647 | -0.08846 |
| C | -2.95471 | -1.55786 | 0.11421 |
| N | -1.62515 | -1.37922 | 0.06173 |
| C | -3.50708 | -2.86599 | 0.36914 |
| C | -4.86111 | -2.96301 | 0.39107 |
| O | 4.11921 | -1.77159 | 2.61310 |
| C | 2.86682 | -1.35868 | 2.09477 |
| C | 2.60410 | -1.96937 | 0.71488 |
| O | 1.36853 | -1.44512 | 0.16021 |
| C | 3.72085 | -1.68667 | -0.32781 |
| O | 4.49432 | -2.81636 | -0.65542 |
| C | 2.92115 | -1.14900 | -1.54657 |
| O | 2.55084 | -2.24105 | -2.38871 |
| C | 1.66850 | -0.53865 | -0.90944 |
| N | 1.82202 | 0.84032 | -0.48203 |
| C | 2.97695 | 1.61159 | -0.29948 |
| N | 2.72358 | 2.85293 | 0.00382 |
| C | 1.34417 | 2.93665 | 0.02836 |
| C | 0.48466 | 4.06811 | 0.26595 |
| O | 0.74305 | 5.22828 | 0.53128 |
| N | -0.88783 | 3.66388 | 0.13830 |
| C | -1.36441 | 2.40631 | -0.14711 |
| N | -2.69545 | 2.23480 | -0.24196 |
| N | -0.54573 | 1.37622 | -0.35173 |

| | | | |
|---|---|---|---|
| C | 0.76469 | 1.70645 | -0.26550 |
| H | -1.25470 | -0.43225 | -0.07473 |
| H | -2.87551 | -3.72972 | 0.53004 |
| H | -5.38346 | -3.89677 | 0.57033 |
| H | 2.91425 | -0.26900 | 2.01190 |
| H | 2.03181 | -1.60066 | 2.76903 |
| H | 2.48138 | -3.05640 | 0.80771 |
| H | 4.40532 | -0.92902 | 0.06579 |
| H | 4.08960 | -3.18499 | -1.45916 |
| H | 3.48414 | -0.45740 | -2.17458 |
| H | 1.82571 | -2.71265 | -1.95064 |
| H | 0.81134 | -0.56690 | -1.58552 |
| H | 3.96524 | 1.19143 | -0.41405 |
| H | -3.32206 | 2.96430 | 0.05808 |
| H | -1.53951 | 4.42690 | 0.27507 |
| H | -3.07913 | 1.27527 | -0.27833 |
| H | -0.97341 | -2.10474 | 0.31095 |
| H | -6.63905 | -1.90868 | 0.20590 |
| H | 4.11597 | -2.73471 | 2.69689 |

Hydrogen only optimization

**Table 77.** Optimized geometry coordinates of A:PRF (S–edge)
Energy: –994406.4016346 (au)

| | | | |
|---|---|---|---|
| O | –5.48211 | –2.28623 | 0.29498 |
| C | –5.02106 | –3.03025 | –0.83108 |
| C | –3.60139 | –2.63028 | –1.16380 |
| O | –3.54557 | –1.17351 | –1.33105 |
| C | –2.57403 | –2.96798 | –0.07303 |
| O | –1.29771 | –3.20720 | –0.67675 |
| C | –2.50505 | –1.65464 | 0.70397 |
| O | –1.34822 | –1.46660 | 1.48240 |
| C | –2.58760 | –0.64315 | –0.43296 |
| N | –2.94801 | 0.69666 | 0.02685 |
| C | –3.96866 | 1.08907 | 0.86705 |
| N | –3.97076 | 2.38163 | 1.11902 |
| C | –2.85857 | 2.87187 | 0.44733 |
| C | –2.27576 | 4.14956 | 0.33001 |
| N | –2.70626 | 5.28255 | 0.90580 |
| N | –1.17758 | 4.24683 | –0.44521 |
| C | –0.67219 | 3.15397 | –1.02572 |
| N | –1.12684 | 1.90989 | –0.97071 |
| C | –2.22597 | 1.83570 | –0.21972 |
| N | 6.26487 | –0.48730 | 0.17408 |
| C | 6.23391 | 0.78380 | –0.42220 |
| N | 5.03011 | 1.39088 | –0.69895 |
| C | 3.79192 | 0.77187 | –0.38815 |

| | | | |
|---|---|---|---|
| C | 3.78874 | –0.51981 | 0.24302 |
| C | 5.07030 | –1.12444 | 0.52609 |
| O | 5.05548 | –2.35696 | 1.11721 |
| C | 2.41697 | –0.92518 | 0.44649 |
| C | 1.91827 | –2.19980 | 1.09869 |
| N | 2.04628 | –3.37626 | 0.24160 |
| C | 1.65575 | 0.13626 | –0.04556 |
| N | 2.47684 | 1.13084 | –0.54513 |
| N | 7.47558 | 1.44416 | –0.75329 |
| H | –5.67271 | –2.87567 | –1.70210 |
| H | –5.06178 | –4.09022 | –0.56336 |
| H | –3.29950 | –3.10065 | –2.10954 |
| H | –2.82030 | –3.84193 | 0.53081 |
| H | –1.23116 | –2.68967 | –1.48946 |
| H | –3.38117 | –1.56710 | 1.35022 |
| H | –0.65897 | –1.97920 | 1.02940 |
| H | –1.61454 | –0.51212 | –0.91997 |
| H | –4.69660 | 0.38942 | 1.25234 |
| H | –2.18358 | 6.12693 | 0.74047 |
| H | 0.21252 | 3.32423 | –1.63897 |
| H | 2.51976 | –2.38782 | 1.99172 |
| H | 0.89509 | –2.07385 | 1.44553 |
| H | 0.58755 | 0.28683 | –0.04882 |
| H | 2.14925 | 1.99539 | –0.94830 |
| H | 7.11753 | –1.00623 | 0.34404 |
| H | –3.51428 | 5.28739 | 1.50554 |
| H | –5.58603 | –1.37954 | –0.02503 |
| H | 3.04474 | –3.54350 | 0.12230 |
| H | 1.66706 | –3.17173 | –0.67953 |
| H | 7.26258 | 2.39605 | –1.03041 |
| H | 8.14647 | 1.41852 | 0.00779 |

**Table S78.** Optimized geometry coordinates of G:C2E (W–edge)
Energy: –2260799.6748452 (au)

| | | | |
|---|---|---|---|
| N | –7.34093 | –1.19860 | 0.01121 |
| C | –7.20819 | –1.82765 | –1.21829 |
| N | –6.15044 | –1.44079 | –1.87927 |
| C | –5.54939 | –0.50999 | –1.04947 |
| C | –4.36785 | 0.26664 | –1.21749 |
| O | –3.57206 | 0.29001 | –2.18201 |
| N | –4.13397 | 1.10670 | –0.12245 |
| C | –4.89802 | 1.15712 | 1.01642 |
| N | –4.48591 | 2.00343 | 1.99674 |
| N | –6.00184 | 0.44673 | 1.19155 |
| C | –6.27578 | –0.34568 | 0.13637 |
| P | 4.25268 | –0.22331 | –1.00199 |

| | | | |
|---|---|---|---|
| O | 5.60428 | –0.70798 | –1.33151 |
| O | 4.01109 | 0.66428 | 0.30331 |
| O | 3.14129 | –1.38152 | –0.77125 |
| C | 3.15751 | –2.58654 | –1.55306 |
| C | 2.16067 | –3.56144 | –0.95314 |
| O | 2.70685 | –4.14956 | 0.26122 |
| C | 0.81344 | –2.97097 | –0.50316 |
| O | –0.12838 | –2.84022 | –1.56824 |
| C | 0.41694 | –3.91982 | 0.64071 |
| O | 0.01738 | –5.20716 | 0.22742 |
| C | 1.79097 | –3.98427 | 1.34005 |
| N | 2.13615 | –2.74711 | 2.08691 |
| C | 3.40159 | –2.19082 | 2.14467 |
| N | 3.42911 | –0.99612 | 2.68939 |
| C | 2.10796 | –0.74405 | 3.03781 |
| C | 1.49066 | 0.40215 | 3.63620 |
| O | 1.99807 | 1.43409 | 4.09817 |
| N | 0.08108 | 0.27628 | 3.62616 |
| C | –0.63013 | –0.77029 | 3.08969 |
| N | –1.97941 | –0.62925 | 2.98624 |
| N | –0.05278 | –1.88446 | 2.69059 |
| C | 1.28869 | –1.80839 | 2.64911 |
| P | –1.07552 | –1.51106 | –1.43074 |
| O | –2.05789 | –1.73471 | –2.64029 |
| O | –1.61246 | –1.27005 | –0.07127 |
| O | –0.06569 | –0.29748 | –1.84086 |
| C | 0.55951 | –0.28527 | –3.13388 |
| C | 1.43298 | 0.94705 | –3.28328 |
| O | 0.60118 | 2.11683 | –3.47076 |
| C | 2.35202 | 1.31192 | –2.10057 |
| O | 3.64661 | 0.69516 | –2.19419 |
| C | 2.47862 | 2.83230 | –2.26781 |
| O | 3.25732 | 3.15265 | –3.40639 |
| C | 1.03036 | 3.16952 | –2.63316 |
| N | 0.13392 | 3.20994 | –1.44506 |
| C | –1.19936 | 2.83764 | –1.49998 |
| N | –1.76482 | 2.77008 | –0.32188 |
| C | –0.76530 | 3.10160 | 0.57188 |
| C | –0.73012 | 2.97525 | 1.98931 |
| O | –1.62597 | 2.58810 | 2.76760 |
| N | 0.54118 | 3.28266 | 2.49935 |
| C | 1.66418 | 3.53665 | 1.74481 |
| N | 2.83304 | 3.68309 | 2.42958 |
| N | 1.63299 | 3.64844 | 0.43359 |
| C | 0.42462 | 3.38254 | –0.10583 |
| H | –7.93005 | –2.55773 | –1.55793 |

| | | | |
|---|---|---|---|
| H | –5.01876 | 1.96149 | 2.85119 |
| H | 4.00177 | 0.15545 | 1.15194 |
| H | 4.16193 | –3.01913 | –1.55293 |
| H | 2.87256 | –2.36341 | –2.58741 |
| H | 2.00154 | –4.36505 | –1.68513 |
| H | 1.00588 | –1.99069 | –0.06461 |
| H | –0.37894 | –3.51276 | 1.25573 |
| H | 0.73517 | –5.62203 | –0.27091 |
| H | 1.89000 | –4.84913 | 1.99923 |
| H | 4.25267 | –2.70598 | 1.72621 |
| H | –2.35489 | –1.20147 | 2.23629 |
| H | 1.17453 | –1.18254 | –3.26112 |
| H | –0.20422 | –0.28274 | –3.91761 |
| H | 2.04744 | 0.82126 | –4.18300 |
| H | 1.87830 | 1.06434 | –1.15012 |
| H | 2.83271 | 3.32954 | –1.36535 |
| H | 4.09441 | 2.67412 | –3.31325 |
| H | 0.94591 | 4.10916 | –3.18337 |
| H | –1.67685 | 2.57792 | –2.43215 |
| H | 3.64157 | 3.68326 | 1.82196 |
| H | –8.06709 | –1.33454 | 0.69823 |
| H | –3.52928 | 2.34594 | 2.05304 |
| H | –3.28983 | 1.70339 | –0.18490 |
| H | 2.92510 | 3.07866 | 3.24402 |
| H | 0.69563 | 3.07396 | 3.48287 |
| H | –2.31810 | 0.32649 | 2.96757 |
| H | –0.44279 | 1.10923 | 3.88480 |
| H | –2.79730 | –1.03948 | –2.59579 |

**Table S79.** Optimized geometry coordinates of G:D2X (W–edge)
Energy: –1793135.2058358 (au)

| | | | |
|---|---|---|---|
| N | –10.48683 | 0.53075 | 0.18844 |
| C | –10.74157 | –0.78569 | 0.48670 |
| N | –9.66106 | –1.46553 | 0.75578 |
| C | –8.62984 | –0.53992 | 0.62503 |
| C | –7.22284 | –0.68451 | 0.80030 |
| O | –6.57279 | –1.69744 | 1.11926 |
| N | –6.56169 | 0.50725 | 0.56183 |
| C | –7.15271 | 1.67955 | 0.19478 |
| N | –6.32872 | 2.70621 | –0.01510 |
| N | –8.44856 | 1.83599 | 0.03889 |
| C | –9.12428 | 0.69207 | 0.26864 |
| O | –3.28043 | 1.77862 | –0.35895 |
| P | –3.03133 | 0.21027 | –0.61253 |
| O | –4.19070 | –0.55546 | 0.19833 |
| O | –3.01046 | –0.14325 | –2.04892 |

| | | | |
|---|---|---|---|
| O | –1.64473 | –0.13163 | 0.12936 |
| P | –0.44205 | –0.82911 | –0.68220 |
| O | 0.18852 | –1.90889 | 0.33212 |
| O | –0.89207 | –1.42586 | –1.95992 |
| O | 0.68588 | 0.30078 | –0.88664 |
| C | 1.18427 | 1.04464 | 0.22777 |
| C | 2.01985 | 0.14868 | 1.14335 |
| C | 3.23466 | –0.42191 | 0.40924 |
| C | 4.31137 | 0.22267 | –0.18846 |
| C | 4.46342 | 1.74555 | –0.19369 |
| S | 3.49327 | –2.12700 | 0.16375 |
| C | 4.99738 | –1.89291 | –0.68615 |
| N | 5.22048 | –0.57802 | –0.74811 |
| C | 6.40607 | –0.03573 | –1.42268 |
| C | 7.53650 | 0.18412 | –0.41480 |
| C | 7.32705 | –0.15430 | 0.91686 |
| N | 8.26175 | 0.02398 | 1.83377 |
| C | 9.45842 | 0.54345 | 1.52765 |
| C | 10.51398 | 0.73849 | 2.61891 |
| N | 9.74777 | 0.90990 | 0.21022 |
| C | 8.76698 | 0.72151 | –0.77011 |
| O | 8.98715 | 1.01099 | –1.94477 |
| H | –11.74757 | –1.18307 | 0.48828 |
| H | –6.79031 | 3.51818 | –0.40182 |
| H | –0.22546 | –2.76981 | 0.15391 |
| H | 0.35798 | 1.48433 | 0.79746 |
| H | 1.76794 | 1.85871 | –0.19953 |
| H | 1.42696 | –0.65684 | 1.56898 |
| H | 2.38811 | 0.75432 | 1.98185 |
| H | 3.84042 | 2.17685 | 0.58872 |
| H | 5.49401 | 2.04378 | 0.00953 |
| H | 4.15816 | 2.17162 | –1.15525 |
| H | 5.63596 | –2.65914 | –1.09880 |
| H | 6.69334 | –0.74661 | –2.20334 |
| H | 6.11766 | 0.88401 | –1.93559 |
| H | 6.39644 | –0.58213 | 1.27547 |
| H | 10.09221 | 1.34099 | 3.42749 |
| H | 11.39629 | 1.22498 | 2.20469 |
| H | 10.78645 | –0.23506 | 3.03633 |
| H | –5.42375 | 2.51212 | –0.42363 |
| H | –5.56554 | 0.46627 | 0.71259 |
| H | –4.62079 | –1.35819 | –0.14822 |
| H | –3.16692 | 2.10059 | 0.55061 |
| H | 9.88936 | 1.36710 | –2.04962 |
| H | –11.14954 | 1.25063 | –0.05831 |

**Table S80.** Optimized geometry coordinates of G:SAM (W–edge)
Energy: –1399444.6204553 (au)

| | | | |
|---|---|---|---|
| N | –8.27926 | 0.35672 | 1.62499 |
| C | –8.76434 | –0.92141 | 1.50541 |
| N | –7.97973 | –1.70760 | 0.82022 |
| C | –6.91211 | –0.90270 | 0.45206 |
| C | –5.73501 | –1.20827 | –0.29406 |
| O | –5.38158 | –2.29415 | –0.78964 |
| N | –4.92538 | –0.08902 | –0.44174 |
| C | –5.19246 | 1.15743 | 0.05672 |
| N | –4.28293 | 2.10811 | –0.21254 |
| N | –6.27187 | 1.45239 | 0.76292 |
| C | –7.08469 | 0.38139 | 0.92181 |
| N | 0.58647 | 0.77279 | –2.31719 |
| C | –0.35082 | –0.33593 | –2.10527 |
| C | –1.57114 | 0.16870 | –1.36438 |
| O | –2.44432 | –0.61139 | –1.01544 |
| O | –1.67643 | 1.34928 | –1.08146 |
| C | 0.29364 | –1.46435 | –1.28873 |
| C | 1.68380 | –1.78886 | –1.81699 |
| S | 2.44038 | –3.25498 | –1.01608 |
| C | 1.01430 | –4.18859 | –0.37247 |
| C | 2.92347 | –2.40700 | 0.48602 |
| C | 4.35852 | –1.95153 | 0.48986 |
| O | 4.50487 | –0.94543 | –0.52664 |
| C | 4.47533 | –1.25740 | 1.78686 |
| O | 5.04711 | –2.13869 | 2.75217 |
| C | 5.43337 | –0.14964 | 1.47517 |
| O | 6.79352 | –0.60025 | 1.64337 |
| C | 5.18263 | 0.22979 | –0.01520 |
| N | 4.26427 | 1.37453 | –0.28438 |
| C | 4.40591 | 2.23471 | –1.34819 |
| N | 3.48373 | 3.15069 | –1.34999 |
| C | 2.67104 | 2.95119 | –0.30571 |
| C | 1.50751 | 3.60580 | 0.19731 |
| N | 0.99007 | 4.72147 | –0.42898 |
| N | 0.91558 | 3.10326 | 1.30145 |
| C | 1.38753 | 1.99208 | 1.91825 |
| N | 2.51480 | 1.37569 | 1.50554 |
| C | 3.16195 | 1.79606 | 0.40645 |
| H | –9.70359 | –1.21405 | 1.95711 |
| H | –3.33206 | 1.84648 | –0.50389 |
| H | –0.68827 | –0.76351 | –3.05924 |
| H | 0.35264 | –1.13828 | –0.24299 |
| H | –0.39292 | –2.31056 | –1.33055 |
| H | 2.39543 | –0.97647 | –1.66696 |

| | | | |
|---|---|---|---|
| H | 1.65701 | –2.02623 | –2.88557 |
| H | 0.41905 | –3.56635 | 0.29272 |
| H | 0.41182 | –4.53373 | –1.21356 |
| H | 1.42462 | –5.05078 | 0.15756 |
| H | 2.25387 | –1.55724 | 0.62233 |
| H | 2.78842 | –3.11052 | 1.31422 |
| H | 5.07977 | –2.76711 | 0.37135 |
| H | 3.53338 | –0.83778 | 2.14685 |
| H | 5.98620 | –1.89362 | 2.80637 |
| H | 5.33096 | 0.70791 | 2.14135 |
| H | 7.03001 | –1.19587 | 0.91691 |
| H | 6.10797 | 0.42345 | –0.56139 |
| H | 5.20812 | 2.12923 | –2.06537 |
| H | 1.30187 | 4.89814 | –1.37203 |
| H | 0.86711 | 1.61368 | 2.78935 |
| H | –4.39898 | 2.95865 | 0.31554 |
| H | –3.96201 | –0.28133 | –0.79627 |
| H | –8.71052 | 1.15021 | 2.07210 |
| H | 0.03191 | 4.94721 | –0.20287 |
| H | 0.64835 | 0.99772 | –3.30592 |
| H | 0.12760 | 1.57328 | –1.87019 |

**Table S81.** Optimized geometry coordinates of G:SFG (W–edge)
Energy: –1184320.9199512 (au)

| | | | |
|---|---|---|---|
| N | –8.31347 | 0.39473 | 1.41170 |
| C | –8.70546 | –0.91206 | 1.29573 |
| N | –7.81714 | –1.67413 | 0.72979 |
| C | –6.77813 | –0.80475 | 0.42727 |
| C | –5.53930 | –1.07111 | –0.19605 |
| O | –5.11596 | –2.15878 | –0.61328 |
| N | –4.76720 | 0.08120 | –0.31631 |
| C | –5.14728 | 1.33216 | 0.11408 |
| N | –4.26745 | 2.32377 | –0.08968 |
| N | –6.30856 | 1.59260 | 0.70248 |
| C | –7.07301 | 0.47998 | 0.82452 |
| N | 0.79512 | 0.79810 | –2.53621 |
| C | 0.19303 | –0.48914 | –2.19626 |
| C | –1.28269 | –0.40094 | –1.80164 |
| O | –1.84004 | –1.38919 | –1.45581 |
| O | –1.79496 | 0.84750 | –1.39620 |
| C | 1.02524 | –1.35158 | –1.22422 |
| C | 1.66782 | –2.67181 | –1.73339 |
| C | 2.08414 | –3.68588 | –0.63485 |
| N | 1.03123 | –4.65269 | –0.32394 |
| C | 2.66020 | –2.97939 | 0.63096 |
| C | 3.99705 | –2.17513 | 0.49533 |

| | | | |
|---|---|---|---|
| O | 4.13911 | –1.21226 | –0.59090 |
| C | 4.48852 | –1.41294 | 1.75764 |
| O | 4.89599 | –2.18935 | 2.89179 |
| C | 5.43571 | –0.29683 | 1.21801 |
| O | 6.81476 | –0.42057 | 1.47061 |
| C | 4.96139 | –0.12137 | –0.22803 |
| N | 4.26002 | 1.17808 | –0.43862 |
| C | 4.57007 | 2.13396 | –1.44864 |
| N | 3.68308 | 3.22879 | –1.32196 |
| C | 2.79941 | 2.95279 | –0.21568 |
| C | 1.70297 | 3.71785 | 0.36143 |
| N | 1.33331 | 5.01975 | –0.19512 |
| N | 1.01350 | 3.21733 | 1.43163 |
| C | 1.35858 | 1.97269 | 1.97069 |
| N | 2.45813 | 1.20411 | 1.39169 |
| C | 3.15440 | 1.70019 | 0.32196 |
| H | –9.66405 | –1.24496 | 1.67270 |
| H | –3.31285 | 2.09850 | –0.32595 |
| H | 1.28358 | 0.77682 | –3.42246 |
| H | 1.44774 | 1.11408 | –1.82722 |
| H | 0.13525 | –1.05567 | –3.13864 |
| H | –1.16379 | 1.49975 | –1.77041 |
| H | 1.81555 | –0.72123 | –0.81352 |
| H | 0.37713 | –1.60506 | –0.37768 |
| H | 2.55253 | –2.42256 | –2.32787 |
| H | 0.96692 | –3.19334 | –2.39586 |
| H | 2.90420 | –4.27778 | –1.06660 |
| H | 1.33247 | –5.26562 | 0.42979 |
| H | 0.18988 | –4.18038 | 0.00420 |
| H | 2.90707 | –3.73871 | 1.38271 |
| H | 1.88928 | –2.34245 | 1.08102 |
| H | 4.75488 | –2.95243 | 0.27607 |
| H | 3.61223 | –0.85636 | 2.11007 |
| H | 5.86198 | –2.26306 | 2.85716 |
| H | 5.18659 | 0.61443 | 1.76573 |
| H | 7.18346 | –1.15190 | 0.95173 |
| H | 5.82291 | –0.07116 | –0.90789 |
| H | 2.09248 | 5.45904 | –0.69965 |
| H | 0.86250 | 5.60726 | 0.48078 |
| H | –3.85735 | –0.06754 | –0.74403 |
| H | –4.45878 | 3.19442 | 0.37865 |
| H | –8.84093 | 1.16789 | 1.78712 |
| H | 5.35801 | 2.02056 | –2.17556 |
| H | 0.82997 | 1.55683 | 2.81477 |

**Table S82.** Optimized geometry coordinates of G:D2X (S–edge)

Energy: –2104605.0032526 (au)

| | | | |
|---|---|---|---|
| O | –0.07538 | 5.50818 | –0.04104 |
| C | 0.80242 | 4.97051 | 0.93313 |
| C | 0.52253 | 3.48810 | 1.02382 |
| O | 0.80015 | 2.83746 | –0.26634 |
| C | 1.35795 | 2.71215 | 2.05924 |
| O | 0.77983 | 2.69075 | 3.33638 |
| C | 1.45147 | 1.31324 | 1.40979 |
| O | 0.24958 | 0.60761 | 1.69787 |
| C | 1.58108 | 1.65268 | –0.08087 |
| N | 2.96934 | 1.92946 | –0.46526 |
| C | 3.48582 | 3.13415 | –0.94608 |
| N | 4.77385 | 3.09204 | –1.15061 |
| C | 5.14282 | 1.81086 | –0.80690 |
| C | 6.45098 | 1.20335 | –0.79503 |
| O | 7.54125 | 1.64591 | –1.08692 |
| N | 6.34633 | –0.15943 | –0.32782 |
| C | 5.20861 | –0.83025 | 0.01304 |
| N | 5.31510 | –2.11772 | 0.41353 |
| N | 4.01004 | –0.24977 | –0.01226 |
| C | 4.04228 | 1.06567 | –0.39601 |
| O | –1.52971 | 2.69629 | –1.55649 |
| P | –2.22673 | 1.35128 | –1.15387 |
| O | –3.05739 | 0.83714 | –2.39524 |
| O | –1.33406 | 0.31168 | –0.55614 |
| O | –3.36774 | 1.80809 | –0.04131 |
| P | –4.97465 | 1.54971 | –0.12638 |
| O | –5.56470 | 2.57563 | 0.93502 |
| O | –5.46478 | 1.60700 | –1.52468 |
| O | –5.21055 | 0.13362 | 0.56337 |
| C | –4.59851 | –0.29629 | 1.80337 |
| C | –4.52193 | –1.82889 | 1.77123 |
| C | –3.71549 | –2.33667 | 0.60637 |
| C | –2.35388 | –2.44317 | 0.49065 |
| C | –1.34647 | –2.23698 | 1.57613 |
| S | –4.49849 | –2.72987 | –0.90176 |
| C | –3.00079 | –3.00842 | –1.63109 |
| N | –1.97772 | –2.81258 | –0.80695 |
| C | –0.58369 | –2.73794 | –1.35069 |
| C | 0.46320 | –3.52908 | –0.61880 |
| C | 0.38843 | –4.88525 | –0.34101 |
| N | 1.35948 | –5.57257 | 0.27540 |
| C | 2.46198 | –4.88454 | 0.59498 |
| C | 3.58182 | –5.63369 | 1.25823 |
| N | 2.65887 | –3.57239 | 0.35421 |
| C | 1.67239 | –2.89111 | –0.23889 |

| | | | |
|---|---|---|---|
| O | 1.80612 | –1.60234 | –0.49090 |
| H | 0.63125 | 5.39467 | 1.93481 |
| H | 1.85933 | 5.13327 | 0.67560 |
| H | –0.54397 | 3.34345 | 1.23283 |
| H | 2.35553 | 3.15507 | 2.15376 |
| H | 0.11514 | 1.98232 | 3.30503 |
| H | 2.29537 | 0.72803 | 1.77957 |
| H | –0.34031 | 0.60457 | 0.91554 |
| H | 1.17519 | 0.86702 | –0.72184 |
| H | 2.83821 | 3.97490 | –1.13704 |
| H | 4.45182 | –2.65819 | 0.51679 |
| H | –5.70179 | 3.47178 | 0.58948 |
| H | –5.22242 | 0.03846 | 2.63608 |
| H | –3.60182 | 0.14391 | 1.89188 |
| H | –5.53619 | –2.23742 | 1.73656 |
| H | –4.07948 | –2.17061 | 2.71218 |
| H | –1.83030 | –1.80486 | 2.45266 |
| H | –0.89903 | –3.18947 | 1.87461 |
| H | –0.54783 | –1.55601 | 1.27875 |
| H | –2.87289 | –3.27787 | –2.66982 |
| H | –0.66546 | –3.05841 | –2.39299 |
| H | –0.32999 | –1.67666 | –1.34173 |
| H | –0.49902 | –5.45750 | –0.61299 |
| H | 3.22905 | –6.60402 | 1.60661 |
| H | 3.98969 | –5.06110 | 2.09563 |
| H | 4.39581 | –5.80081 | 0.54288 |
| H | 0.10577 | 6.45197 | –0.13574 |
| H | 6.16071 | –2.62814 | 0.21301 |
| H | 7.24461 | –0.62368 | –0.25929 |
| H | 2.70650 | –1.18000 | –0.21493 |
| H | –3.99557 | 1.16515 | –2.42016 |
| H | –0.61766 | 2.82721 | –1.16380 |

**Table S83.** Optimized geometry coordinates of G:D2X (S–edge)
Energy: –2104605.0032526 (au)

| | | | |
|---|---|---|---|
| O | –0.07538 | 5.50818 | –0.04104 |
| C | 0.80242 | 4.97051 | 0.93313 |
| C | 0.52253 | 3.48810 | 1.02382 |
| O | 0.80015 | 2.83746 | –0.26634 |
| C | 1.35795 | 2.71215 | 2.05924 |
| O | 0.77983 | 2.69075 | 3.33638 |
| C | 1.45147 | 1.31324 | 1.40979 |
| O | 0.24958 | 0.60761 | 1.69787 |
| C | 1.58108 | 1.65268 | –0.08087 |
| N | 2.96934 | 1.92946 | –0.46526 |
| C | 3.48582 | 3.13415 | –0.94608 |

| | | | |
|---|---|---|---|
| N | 4.77385 | 3.09204 | –1.15061 |
| C | 5.14282 | 1.81086 | –0.80690 |
| C | 6.45098 | 1.20335 | –0.79503 |
| O | 7.54125 | 1.64591 | –1.08692 |
| N | 6.34633 | –0.15943 | –0.32782 |
| C | 5.20861 | –0.83025 | 0.01304 |
| N | 5.31510 | –2.11772 | 0.41353 |
| N | 4.01004 | –0.24977 | –0.01226 |
| C | 4.04228 | 1.06567 | –0.39601 |
| O | –1.52971 | 2.69629 | –1.55649 |
| P | –2.22673 | 1.35128 | –1.15387 |
| O | –3.05739 | 0.83714 | –2.39524 |
| O | –1.33406 | 0.31168 | –0.55614 |
| O | –3.36774 | 1.80809 | –0.04131 |
| P | –4.97465 | 1.54971 | –0.12638 |
| O | –5.56470 | 2.57563 | 0.93502 |
| O | –5.46478 | 1.60700 | –1.52468 |
| O | –5.21055 | 0.13362 | 0.56337 |
| C | –4.59851 | –0.29629 | 1.80337 |
| C | –4.52193 | –1.82889 | 1.77123 |
| C | –3.71549 | –2.33667 | 0.60637 |
| C | –2.35388 | –2.44317 | 0.49065 |
| C | –1.34647 | –2.23698 | 1.57613 |
| S | –4.49849 | –2.72987 | –0.90176 |
| C | –3.00079 | –3.00842 | –1.63109 |
| N | –1.97772 | –2.81258 | –0.80695 |
| C | –0.58369 | –2.73794 | –1.35069 |
| C | 0.46320 | –3.52908 | –0.61880 |
| C | 0.38843 | –4.88525 | –0.34101 |
| N | 1.35948 | –5.57257 | 0.27540 |
| C | 2.46198 | –4.88454 | 0.59498 |
| C | 3.58182 | –5.63369 | 1.25823 |
| N | 2.65887 | –3.57239 | 0.35421 |
| C | 1.67239 | –2.89111 | –0.23889 |
| O | 1.80612 | –1.60234 | –0.49090 |
| H | 0.63125 | 5.39467 | 1.93481 |
| H | 1.85933 | 5.13327 | 0.67560 |
| H | –0.54397 | 3.34345 | 1.23283 |
| H | 2.35553 | 3.15507 | 2.15376 |
| H | 0.11514 | 1.98232 | 3.30503 |
| H | 2.29537 | 0.72803 | 1.77957 |
| H | –0.34031 | 0.60457 | 0.91554 |
| H | 1.17519 | 0.86702 | –0.72184 |
| H | 2.83821 | 3.97490 | –1.13704 |
| H | 4.45182 | –2.65819 | 0.51679 |
| H | –5.70179 | 3.47178 | 0.58948 |

| | | | |
|---|---|---|---|
| H | –5.22242 | 0.03846 | 2.63608 |
| H | –3.60182 | 0.14391 | 1.89188 |
| H | –5.53619 | –2.23742 | 1.73656 |
| H | –4.07948 | –2.17061 | 2.71218 |
| H | –1.83030 | –1.80486 | 2.45266 |
| H | –0.89903 | –3.18947 | 1.87461 |
| H | –0.54783 | –1.55601 | 1.27875 |
| H | –2.87289 | –3.27787 | –2.66982 |
| H | –0.66546 | –3.05841 | –2.39299 |
| H | –0.32999 | –1.67666 | –1.34173 |
| H | –0.49902 | –5.45750 | –0.61299 |
| H | 3.22905 | –6.60402 | 1.60661 |
| H | 3.98969 | –5.06110 | 2.09563 |
| H | 4.39581 | –5.80081 | 0.54288 |
| H | 0.10577 | 6.45197 | –0.13574 |
| H | 6.16071 | –2.62814 | 0.21301 |
| H | 7.24461 | –0.62368 | –0.25929 |
| H | 2.70650 | –1.18000 | –0.21493 |
| H | –3.99557 | 1.16515 | –2.42016 |
| H | –0.61766 | 2.82721 | –1.16380 |

**Table S84.** Optimized geometry coordinates of G:RS3 (S–edge)
Energy: –1545960.0516360 (au)

| | | | |
|---|---|---|---|
| O | 5.30881 | 4.23024 | 1.61122 |
| C | 4.37805 | 4.51788 | 0.58779 |
| C | 3.53437 | 3.29473 | 0.32651 |
| O | 4.32469 | 2.33263 | –0.40233 |
| C | 3.09310 | 2.58394 | 1.59125 |
| O | 1.81339 | 3.05522 | 1.94054 |
| C | 3.03999 | 1.11435 | 1.19448 |
| O | 1.74203 | 0.74694 | 0.75201 |
| C | 4.04468 | 1.02111 | 0.04590 |
| N | 5.31276 | 0.35347 | 0.36127 |
| C | 6.39055 | 0.86408 | 1.04182 |
| N | 7.38507 | 0.02657 | 1.14950 |
| C | 6.94127 | –1.10671 | 0.48630 |
| C | 7.58123 | –2.34777 | 0.26671 |
| O | 8.70991 | –2.69383 | 0.63890 |
| N | 6.78275 | –3.23936 | –0.44434 |
| C | 5.51547 | –2.95625 | –0.89281 |
| N | 4.87959 | –3.91574 | –1.57348 |
| N | 4.90742 | –1.79560 | –0.70305 |
| C | 5.67285 | –0.92216 | –0.00289 |
| O | –8.62790 | –0.78503 | 1.00908 |
| C | –7.49797 | –1.33314 | 1.02338 |
| N | –7.36248 | –2.62891 | 1.40219 |

| | | | |
|---|---|---|---|
| C | –6.14404 | –3.22649 | 1.41717 |
| O | –6.05488 | –4.42927 | 1.76902 |
| C | –6.37125 | –0.61084 | 0.64699 |
| C | –5.11502 | –1.22486 | 0.66340 |
| N | –5.03094 | –2.53086 | 1.04944 |
| N | –6.49479 | 0.68124 | 0.27317 |
| C | –5.40090 | 1.38179 | –0.09001 |
| C | –5.53732 | 2.71125 | –0.47397 |
| C | –4.42829 | 3.46044 | –0.85123 |
| C | –4.64832 | 4.89514 | –1.25695 |
| C | –3.15752 | 2.87132 | –0.84485 |
| N | –2.02284 | 3.54476 | –1.20280 |
| C | –0.74222 | 2.86403 | –1.17044 |
| C | –2.03758 | 4.93523 | –1.62615 |
| C | –3.03603 | 1.53138 | –0.45712 |
| C | –4.14558 | 0.77593 | –0.07922 |
| N | –4.01663 | –0.51899 | 0.29033 |
| C | –2.68210 | –1.14890 | 0.31163 |
| C | –2.38811 | –1.83456 | –1.02113 |
| O | –3.41611 | –2.79613 | –1.26232 |
| C | –1.03564 | –2.55247 | –1.05261 |
| O | –0.93222 | –3.44560 | 0.05026 |
| C | 0.18066 | –1.63665 | –1.00566 |
| O | –0.15678 | –0.34106 | –1.49145 |
| C | 1.30640 | –2.19773 | –1.86361 |
| O | 1.97774 | –3.23632 | –1.14280 |
| H | 3.71315 | 5.34482 | 0.88444 |
| H | 4.87261 | 4.80492 | –0.34869 |
| H | 2.64882 | 3.57595 | –0.26121 |
| H | 3.82338 | 2.74472 | 2.38692 |
| H | 1.74455 | 3.12886 | 2.90082 |
| H | 3.34162 | 0.45108 | 2.01512 |
| H | 1.14614 | 1.36700 | 1.21021 |
| H | 3.60201 | 0.42158 | –0.75730 |
| H | 6.37789 | 1.87990 | 1.41332 |
| H | 3.86839 | –3.78870 | –1.62564 |
| H | –6.53210 | 3.14052 | –0.46905 |
| H | –5.72335 | 5.09329 | –1.17129 |
| H | –4.17736 | 5.62639 | –0.60535 |
| H | –4.40803 | 5.11283 | –2.29412 |
| H | 0.03591 | 3.55335 | –1.49538 |
| H | –0.48558 | 2.52769 | –0.15726 |
| H | –0.72887 | 1.99537 | –1.83718 |
| H | –2.58813 | 5.10416 | –2.54723 |
| H | –2.35752 | 5.62491 | –0.85001 |
| H | –1.00314 | 5.20767 | –1.85084 |

| | | | |
|---|---|---|---|
| H | –2.07441 | 1.05690 | –0.53959 |
| H | –1.94188 | –0.40339 | 0.59360 |
| H | –2.73025 | –1.90990 | 1.09430 |
| H | –2.42565 | –1.11136 | –1.84094 |
| H | –3.70162 | –3.17989 | –0.41302 |
| H | –1.04520 | –3.18064 | –1.94719 |
| H | –0.73521 | –2.93595 | 0.84608 |
| H | 0.55063 | –1.55635 | 0.02568 |
| H | 0.55580 | 0.23805 | –1.17754 |
| H | 2.06798 | –1.43385 | –2.04337 |
| H | 0.92424 | –2.54024 | –2.83533 |
| H | 1.30251 | –3.85460 | –0.82155 |
| H | 6.09875 | 4.76798 | 1.48443 |
| H | 7.21771 | –4.13400 | –0.63121 |
| H | 5.19724 | –4.86870 | –1.48453 |
| H | –8.18422 | –3.15317 | 1.66911 |

**Table S85.** Optimized geometry coordinates of G:SAH (S–edge)
Energy: –1686163.9360389 (au)

| | | | |
|---|---|---|---|
| O | 5.02180 | 4.11873 | –1.10652 |
| C | 3.74558 | 4.70739 | –0.90411 |
| C | 2.75465 | 3.68237 | –0.43012 |
| O | 2.75318 | 2.55024 | –1.33532 |
| C | 3.05913 | 3.03485 | 0.91817 |
| O | 2.70660 | 3.84336 | 2.01375 |
| C | 2.22210 | 1.76985 | 0.84810 |
| O | 0.83853 | 2.08321 | 0.96614 |
| C | 2.50403 | 1.36590 | –0.59898 |
| N | 3.67718 | 0.47710 | –0.69232 |
| C | 4.92346 | 0.72128 | –1.21807 |
| N | 5.71985 | –0.31770 | –1.14140 |
| C | 4.94956 | –1.28795 | –0.52299 |
| C | 5.24500 | –2.61720 | –0.17305 |
| O | 6.29224 | –3.25446 | –0.33589 |
| N | 4.15850 | –3.23236 | 0.42124 |
| C | 2.94441 | –2.64400 | 0.65994 |
| N | 2.00679 | –3.39995 | 1.26261 |
| N | 2.65545 | –1.40278 | 0.32856 |
| C | 3.70172 | –0.80601 | –0.25084 |
| N | 0.18699 | –0.25359 | 1.45698 |
| C | –0.68012 | –0.37214 | 2.63251 |
| C | –2.10758 | 0.15553 | 2.43935 |
| C | –2.36148 | 1.57816 | 1.97230 |
| S | –4.01538 | 2.02153 | 2.55179 |
| C | –0.80373 | –1.79530 | 3.18054 |
| O | –1.44189 | –1.96435 | 4.21997 |

| | | | |
|---|---|---|---|
| O | –0.30483 | –2.77838 | 2.61870 |
| C | –4.63667 | 0.71930 | 1.46111 |
| C | –4.86018 | 1.16032 | 0.01065 |
| O | –3.66407 | 1.61683 | –0.63124 |
| C | –5.40766 | 0.01299 | –0.82595 |
| O | –6.81784 | 0.02240 | –0.85181 |
| C | –4.78032 | 0.20914 | –2.20003 |
| O | –5.60214 | 0.99278 | –3.04154 |
| C | –3.52605 | 1.00780 | –1.90462 |
| N | –2.30240 | 0.17513 | –1.96583 |
| C | –1.04836 | 0.58992 | –2.35759 |
| N | –0.20269 | –0.47014 | –2.30849 |
| C | –0.89790 | –1.54520 | –1.87478 |
| C | –0.53539 | –2.85579 | –1.63527 |
| N | 0.70825 | –3.24731 | –1.83441 |
| N | –1.46087 | –3.77400 | –1.20656 |
| C | –2.75856 | –3.37287 | –1.01256 |
| N | –3.11211 | –2.05812 | –1.23257 |
| C | –2.21210 | –1.16028 | –1.67115 |
| H | 3.37427 | 5.15672 | –1.83824 |
| H | 3.77136 | 5.50042 | –0.14092 |
| H | 1.76623 | 4.17171 | –0.40595 |
| H | 4.12504 | 2.80244 | 0.98374 |
| H | 1.75597 | 3.71649 | 2.15427 |
| H | 2.44601 | 1.00775 | 1.58790 |
| H | 0.57550 | 2.63985 | 0.21852 |
| H | 1.65992 | 0.82285 | –1.04050 |
| H | 5.17290 | 1.68697 | –1.63245 |
| H | 1.11133 | –2.99066 | 1.54139 |
| H | 0.18377 | 0.66723 | 1.07965 |
| H | 1.09369 | –0.69167 | 1.51058 |
| H | –0.26277 | 0.19262 | 3.50682 |
| H | –2.51325 | –0.55137 | 1.71530 |
| H | –2.65890 | 0.01562 | 3.37443 |
| H | –2.28946 | 1.69404 | 0.89662 |
| H | –1.64429 | 2.26240 | 2.44221 |
| H | –5.64405 | 0.45675 | 1.81202 |
| H | –4.06827 | –0.19403 | 1.51289 |
| H | –5.58306 | 1.98743 | 0.00930 |
| H | –5.06290 | –0.94670 | –0.43329 |
| H | –7.07105 | 0.37289 | –1.72142 |
| H | –4.59680 | –0.72635 | –2.72313 |
| H | –5.67661 | 1.88612 | –2.67497 |
| H | –3.38149 | 1.78641 | –2.66089 |
| H | –0.81684 | 1.59013 | –2.68920 |
| H | –3.50499 | –4.07744 | –0.67147 |

| | | | |
|---|---|---|---|
| H | 5.67417 | 4.82529 | –1.17981 |
| H | 4.31816 | –4.19899 | 0.67676 |
| H | 2.26793 | –4.25199 | 1.73203 |
| H | –1.77340 | –1.14062 | 4.60948 |
| H | 1.42057 | –2.53735 | –1.91878 |
| H | 0.97884 | –4.14467 | –1.46151 |

**Table S86.** Optimized geometry coordinates of U:FFO (W–edge)
Energy: –1318178.4319365 (au)

| | | | |
|---|---|---|---|
| N | –6.69664 | 3.96290 | –0.38015 |
| C | –6.06141 | 2.78808 | –0.02966 |
| O | –4.98767 | 2.75277 | 0.54555 |
| N | –6.72183 | 1.64004 | –0.36557 |
| C | –7.92795 | 1.54315 | –1.01021 |
| O | –8.37525 | 0.42241 | –1.22965 |
| C | –8.52872 | 2.79589 | –1.35859 |
| C | –7.91086 | 3.93934 | –1.03230 |
| N | –5.42532 | –0.96701 | 0.02292 |
| C | –5.70165 | –2.00810 | –0.76482 |
| N | –7.00235 | –2.03880 | –1.47842 |
| N | –4.84472 | –3.05529 | –0.93073 |
| C | –3.66082 | –3.07228 | –0.30359 |
| O | –2.81728 | –4.14688 | –0.50281 |
| C | –3.31613 | –1.98358 | 0.52831 |
| N | –2.12631 | –1.85774 | 1.27162 |
| C | –1.58468 | –0.61646 | 1.52973 |
| C | –2.61014 | 0.20342 | 2.23644 |
| N | –3.90842 | 0.24464 | 1.56815 |
| C | –4.24155 | –0.92863 | 0.67877 |
| C | –1.19644 | 0.04295 | 0.20769 |
| N | –0.17416 | –0.65161 | –0.58187 |
| C | 3.77725 | 0.17050 | 0.54548 |
| C | 3.55829 | –0.67441 | –0.54407 |
| C | 2.25205 | –0.95973 | –0.92905 |
| C | 1.19087 | –0.38842 | –0.22343 |
| C | 1.42379 | 0.45660 | 0.86277 |
| C | 2.72004 | 0.74455 | 1.23578 |
| C | 5.18960 | 0.54668 | 1.01673 |
| O | 5.44331 | 0.64249 | 2.25474 |
| N | 6.22161 | 0.82913 | 0.04080 |
| C | 7.50462 | 1.22742 | 0.52526 |
| C | 8.68039 | 0.25482 | 0.18103 |
| C | 8.93142 | –0.08418 | –1.22962 |
| C | 8.28827 | –1.31537 | –1.74117 |
| O | 8.94489 | –2.44120 | –1.76345 |
| O | 7.09534 | –1.25063 | –2.14463 |

| | | | |
|---|---|---|---|
| C | 7.71011 | 2.56351 | –0.06637 |
| O | 7.51372 | 2.73807 | –1.35541 |
| O | 8.03997 | 3.51763 | 0.66551 |
| C | –1.45677 | –3.03682 | 1.86989 |
| O | –0.45451 | –2.96109 | 2.52160 |
| H | 8.52316 | –0.64921 | 0.78169 |
| H | 9.58292 | 0.72056 | 0.59984 |
| H | 8.66043 | 0.76449 | –1.87145 |
| H | 9.99870 | –0.26721 | –1.36894 |
| H | 7.41645 | 1.27006 | 1.61687 |
| H | –9.48025 | 2.79448 | –1.87166 |
| H | –8.32618 | 4.91314 | –1.26715 |
| H | –2.27402 | 1.23981 | 2.36487 |
| H | –2.74254 | –0.20339 | 3.24845 |
| H | –2.09434 | 0.08830 | –0.41742 |
| H | –0.89414 | 1.08200 | 0.33857 |
| H | –0.39377 | –1.60520 | –0.84041 |
| H | 4.39964 | –1.10564 | –1.07703 |
| H | 2.04628 | –1.62140 | –1.76504 |
| H | 0.62356 | 0.88612 | 1.44146 |
| H | 2.93146 | 1.37895 | 2.08949 |
| H | –1.98760 | –3.95772 | 1.60465 |
| H | –5.03516 | –3.82115 | –1.56592 |
| H | –7.47084 | –2.93411 | –1.41392 |
| H | –7.57795 | –1.23270 | –1.22501 |
| H | –6.26624 | 0.73877 | –0.13258 |
| H | –6.22796 | 4.82279 | –0.13556 |
| H | –4.28399 | 1.13420 | 1.24462 |
| H | –0.71352 | –0.76501 | 2.17208 |
| H | 5.99017 | 1.02219 | –0.92366 |
| H | 8.09664 | 3.26854 | 1.60105 |
| H | 6.84839 | –2.14773 | –2.43647 |

**Table S87.** Optimized geometry coordinates of 2U:SAM (W–edge)
Energy: –1319523.9736398 (au)

| | | | |
|---|---|---|---|
| N | –5.78540 | 3.75056 | –0.64023 |
| C | –5.02023 | 2.85072 | 0.06863 |
| O | –5.27791 | 2.54018 | 1.20843 |
| N | –3.94428 | 2.33826 | –0.60283 |
| C | –3.55795 | 2.63745 | –1.88693 |
| O | –2.57795 | 2.06800 | –2.36635 |
| C | –4.38600 | 3.60218 | –2.55590 |
| C | –5.44562 | 4.11406 | –1.92346 |
| N | 6.85893 | 1.61310 | 1.58549 |
| C | 6.22343 | 2.24274 | 0.40084 |
| C | 6.07932 | 3.74858 | 0.61228 |

| | | | |
|---|---|---|---|
| O | 5.61370 | 4.45315 | –0.27725 |
| O | 6.36199 | 4.25646 | 1.68875 |
| C | 4.82073 | 1.64234 | 0.17425 |
| C | 4.85514 | 0.31375 | –0.60503 |
| S | 3.58661 | –0.79875 | 0.09150 |
| C | 4.68632 | –2.08042 | 0.74688 |
| C | 2.96979 | –1.66896 | –1.37694 |
| C | 1.88087 | –2.63311 | –0.91371 |
| O | 0.94078 | –1.89040 | –0.17127 |
| C | 1.12028 | –3.24282 | –2.09608 |
| O | 1.53022 | –4.58905 | –2.36725 |
| C | –0.36833 | –3.19872 | –1.63648 |
| O | –0.97569 | –4.46910 | –1.90238 |
| C | –0.26510 | –2.63817 | –0.17634 |
| N | –1.38674 | –1.76622 | 0.19930 |
| C | –1.79897 | –0.62982 | –0.44923 |
| N | –2.82988 | –0.10447 | 0.16425 |
| C | –3.14396 | –0.85999 | 1.24186 |
| C | –4.12925 | –0.80497 | 2.26256 |
| N | –5.05733 | 0.20450 | 2.28109 |
| N | –4.14401 | –1.75752 | 3.21305 |
| C | –3.24827 | –2.74302 | 3.22603 |
| N | –2.31622 | –2.84921 | 2.27300 |
| C | –2.21942 | –1.93933 | 1.28135 |
| H | –4.13562 | 3.89286 | –3.56660 |
| H | –6.10653 | 4.83983 | –2.38475 |
| H | 6.88492 | 2.56521 | 2.17483 |
| H | 7.81682 | 1.28831 | 1.44874 |
| H | 6.85970 | 2.07482 | –0.47190 |
| H | 4.31681 | 1.54279 | 1.14441 |
| H | 4.26630 | 2.39859 | –0.38752 |
| H | 5.81306 | –0.20828 | –0.54121 |
| H | 4.62266 | 0.46321 | –1.66227 |
| H | 5.34970 | –2.44371 | –0.03932 |
| H | 5.25791 | –1.65065 | 1.57135 |
| H | 4.07783 | –2.89491 | 1.14314 |
| H | 3.79352 | –2.17407 | –1.88702 |
| H | 2.53835 | –0.89568 | –2.01899 |
| H | 2.29819 | –3.46557 | –0.32732 |
| H | 1.25053 | –2.62869 | –2.99832 |
| H | 0.71212 | –5.09463 | –2.51325 |
| H | –0.88748 | –2.43157 | –2.22196 |
| H | –1.93338 | –4.35899 | –1.96805 |
| H | –0.24399 | –3.44660 | 0.56574 |
| H | –1.32764 | –0.20952 | –1.32521 |
| H | –5.66184 | 0.21019 | 3.09067 |

| | | | |
|---|---|---|---|
| H | –3.28154 | –3.48868 | 4.01270 |
| H | –3.46507 | 1.51196 | –0.18787 |
| H | –6.57162 | 4.15776 | –0.15496 |
| H | –4.94743 | 1.08205 | 1.77411 |
| H | 6.31476 | 0.91208 | 2.08610 |

**Table S88.** Optimized geometry coordinates of C:5GP (W–edge)

Energy: –1255944.4350053 (au)

| | | | |
|---|---|---|---|
| N | 6.04557 | –2.55961 | –0.94998 |
| C | 5.75996 | –1.18045 | –0.76377 |
| O | 6.63283 | –0.35435 | –0.98877 |
| N | 4.49191 | –0.88219 | –0.33611 |
| C | 3.58345 | –1.83477 | –0.12673 |
| N | 2.36672 | –1.45420 | 0.30352 |
| C | 3.87577 | –3.23364 | –0.32713 |
| C | 5.13070 | –3.54241 | –0.74413 |
| P | –4.76666 | –1.00038 | –1.86375 |
| O | –4.97819 | 0.56214 | –1.47544 |
| O | –6.01340 | –1.77971 | –1.94431 |
| O | –3.83144 | –0.87088 | –3.17117 |
| O | –3.71391 | –1.58392 | –0.80697 |
| C | –2.42511 | –0.97312 | –0.60432 |
| C | –1.91640 | –1.46224 | 0.74797 |
| O | –0.53350 | –1.05266 | 0.89587 |
| C | –2.68172 | –0.89722 | 1.98069 |
| O | –3.45797 | –1.84043 | 2.67143 |
| C | –1.53201 | –0.32993 | 2.86258 |
| O | –1.06642 | –1.35240 | 3.74202 |
| C | –0.44031 | 0.01599 | 1.84673 |
| N | –0.56536 | 1.33478 | 1.25019 |
| C | –1.62804 | 2.24803 | 1.27229 |
| N | –1.34983 | 3.37576 | 0.67920 |
| C | –0.04839 | 3.23048 | 0.23654 |
| C | 0.81563 | 4.15884 | –0.45325 |
| O | 0.61661 | 5.29428 | –0.84595 |
| N | 2.11382 | 3.56603 | –0.64446 |
| C | 2.51696 | 2.30290 | –0.27471 |
| N | 3.78072 | 1.93142 | –0.52627 |
| N | 1.67455 | 1.46531 | 0.31746 |
| C | 0.45346 | 1.97904 | 0.57231 |
| H | 2.13740 | –0.45447 | 0.32935 |
| H | 3.13315 | –4.00137 | –0.15405 |
| H | 5.45858 | –4.56009 | –0.92739 |
| H | –2.51026 | 0.11834 | –0.61759 |
| H | –1.74392 | –1.28392 | –1.40118 |
| H | –1.95095 | –2.55607 | 0.76915 |

| | | | |
|---|---|---|---|
| H | –3.35482 | –0.09571 | 1.65651 |
| H | –2.89045 | –2.17872 | 3.38467 |
| H | –1.83509 | 0.50347 | 3.49762 |
| H | –0.53628 | –1.97205 | 3.21750 |
| H | 0.55980 | –0.04745 | 2.28064 |
| H | –2.56651 | 2.01714 | 1.75432 |
| H | 4.38602 | 2.51592 | –1.07934 |
| H | 6.98305 | –2.76592 | –1.26561 |
| H | 1.59241 | –2.09809 | 0.31878 |
| H | 4.07331 | 0.95056 | –0.37548 |
| H | 2.76992 | 4.19312 | –1.09384 |
| H | –5.83661 | 0.67636 | –1.04179 |
| H | –4.29607 | –0.46206 | –3.91515 |

**Table S89.** Optimized geometry coordinates of C:5GP (W–edge)
Energy: –1255956.1817503 (au)

| | | | |
|---|---|---|---|
| N | 8.23780 | 0.35791 | 0.94029 |
| C | 6.87015 | 0.62137 | 0.71278 |
| O | 6.42199 | 1.71594 | 1.06835 |
| N | 6.13727 | –0.35597 | 0.11269 |
| C | 6.69566 | –1.52701 | –0.22840 |
| N | 5.91172 | –2.44660 | –0.79728 |
| C | 8.09805 | –1.79639 | –0.00048 |
| C | 8.82810 | –0.81713 | 0.58999 |
| P | –6.99524 | –1.36979 | 0.52141 |
| O | –6.39053 | –2.62710 | –0.31667 |
| O | –8.37407 | –1.00885 | 0.14723 |
| O | –6.68817 | –1.85694 | 2.02987 |
| O | –5.95864 | –0.16066 | 0.35624 |
| C | –4.57434 | –0.29096 | 0.74071 |
| C | –3.88533 | 1.04657 | 0.48585 |
| O | –2.54671 | 0.89899 | 0.98596 |
| C | –3.75602 | 1.47570 | –0.98735 |
| O | –4.84958 | 2.29442 | –1.32656 |
| C | –2.37508 | 2.17698 | –1.02232 |
| O | –2.58445 | 3.54642 | –0.72533 |
| C | –1.58823 | 1.45623 | 0.10133 |
| N | –0.67011 | 0.43113 | –0.37618 |
| C | –0.95572 | –0.79346 | –0.97597 |
| N | 0.10664 | –1.50556 | –1.23859 |
| C | 1.16135 | –0.73162 | –0.78351 |
| C | 2.57228 | –0.97519 | –0.77348 |
| O | 3.18467 | –1.96863 | –1.19543 |
| N | 3.27582 | 0.10028 | –0.19910 |
| C | 2.71379 | 1.25274 | 0.30278 |
| N | 3.56319 | 2.18167 | 0.79219 |

| | | | |
|---|---|---|---|
| N | 1.40736 | 1.48553 | 0.30134 |
| C | 0.70171 | 0.47101 | –0.24082 |
| H | 4.90386 | –2.26002 | –0.95779 |
| H | 8.54646 | –2.73936 | –0.28475 |
| H | 9.88571 | –0.91510 | 0.80816 |
| H | –4.09901 | –1.08406 | 0.15247 |
| H | –4.50299 | –0.54935 | 1.80044 |
| H | –4.40838 | 1.84137 | 1.03138 |
| H | –3.75167 | 0.60770 | –1.65601 |
| H | –4.57048 | 3.20446 | –1.13708 |
| H | –1.87364 | 2.05810 | –1.98971 |
| H | –1.78346 | 4.04061 | –0.93765 |
| H | –0.96080 | 2.17376 | 0.63904 |
| H | –1.97297 | –1.09567 | –1.18086 |
| H | 4.56845 | 2.01731 | 0.88945 |
| H | 4.30101 | –0.02233 | –0.12816 |
| H | 3.14268 | 2.98351 | 1.23143 |
| H | 6.29501 | –3.33812 | –1.06597 |
| H | 8.75629 | 1.10259 | 1.38468 |
| H | –6.84566 | –2.69261 | –1.16902 |
| H | –7.24415 | –2.60632 | 2.28517 |

**Table S90.** Optimized geometry coordinates of C:C2E (E–edge)
Energy: –2167970.4510178 (au)

| | | | |
|---|---|---|---|
| N | 8.36566 | 0.09373 | 1.37392 |
| C | 7.02074 | –0.26753 | 1.14856 |
| O | 6.28119 | –0.54505 | 2.09986 |
| N | 6.55399 | –0.30109 | –0.12554 |
| C | 7.35241 | 0.00237 | –1.14832 |
| N | 6.82827 | –0.05369 | –2.37733 |
| C | 8.72331 | 0.37027 | –0.94775 |
| C | 9.17969 | 0.40424 | 0.31207 |
| P | –3.78426 | –0.40333 | –2.75174 |
| O | –5.00763 | –1.05001 | –3.87299 |
| O | –2.60713 | 0.57696 | –3.64469 |
| O | –3.04390 | –1.60690 | –1.98350 |
| C | –3.82031 | –2.64407 | –1.38792 |
| C | –2.97983 | –3.46410 | –0.42293 |
| O | –1.90483 | –4.12521 | –1.11356 |
| C | –2.32846 | –2.64293 | 0.68043 |
| O | –3.15457 | –2.55212 | 1.84059 |
| C | –1.05923 | –3.41585 | 0.96780 |
| O | –1.34060 | –4.47827 | 1.88105 |
| C | –0.68540 | –4.03829 | –0.35778 |
| N | 0.30384 | –3.19452 | –1.09056 |
| C | 0.14398 | –2.73717 | –2.35363 |

| | | | |
|---|---|---|---|
| N | 1.21914 | –2.00984 | –2.75821 |
| C | 2.09381 | –1.98269 | –1.74303 |
| C | 3.42323 | –1.38284 | –1.51856 |
| O | 4.04160 | –0.65067 | –2.48651 |
| N | 4.01924 | –1.58489 | –0.32337 |
| C | 3.43204 | –2.31219 | 0.66404 |
| N | 4.10023 | –2.45477 | 1.83661 |
| N | 2.20237 | –2.88817 | 0.51692 |
| C | 1.50025 | –2.76356 | –0.64591 |
| P | –2.92854 | –1.39256 | 2.93730 |
| O | –3.98622 | –1.63337 | 4.34467 |
| O | –1.22318 | –1.28972 | 3.43416 |
| O | –3.35564 | –0.04033 | 2.17868 |
| C | –4.67905 | 0.13378 | 1.68023 |
| C | –4.68232 | 1.28305 | 0.68454 |
| O | –4.17342 | 2.45979 | 1.31814 |
| C | –3.77705 | 1.03210 | –0.51144 |
| O | –4.50430 | 0.45023 | –1.58752 |
| C | –3.24438 | 2.40612 | –0.85929 |
| O | –4.13609 | 3.06871 | –1.75838 |
| C | –3.28925 | 3.16726 | 0.44265 |
| N | –1.93030 | 3.29388 | 1.04439 |
| C | –1.49992 | 2.72647 | 2.19576 |
| N | –0.20776 | 3.07208 | 2.45783 |
| C | 0.19978 | 3.87793 | 1.46044 |
| C | 1.44342 | 4.59399 | 1.11205 |
| O | 2.52991 | 4.50969 | 1.91032 |
| N | 1.47292 | 5.33421 | –0.02194 |
| C | 0.38368 | 5.43172 | –0.83921 |
| N | 0.45911 | 6.19097 | –1.96419 |
| N | –0.78334 | 4.78960 | –0.56824 |
| C | –0.93029 | 4.02140 | 0.53902 |
| H | 5.81375 | –0.27587 | –2.47374 |
| H | 9.36934 | 0.61533 | –1.78112 |
| H | 10.20068 | 0.67446 | 0.55958 |
| H | –4.21989 | –3.27179 | –2.18808 |
| H | –4.66732 | –2.22075 | –0.83584 |
| H | –3.64088 | –4.21638 | 0.02827 |
| H | –2.08931 | –1.64297 | 0.30472 |
| H | –0.24869 | –2.78704 | 1.34989 |
| H | –1.42955 | –4.09441 | 2.76265 |
| H | –0.27118 | –5.03948 | –0.22341 |
| H | –0.73368 | –2.94157 | –2.94045 |
| H | 3.55131 | –2.83554 | 2.59092 |
| H | –5.03346 | –0.77637 | 1.18402 |
| H | –5.33627 | 0.34190 | 2.52841 |

| | | | |
|---|---|---|---|
| H | –5.71275 | 1.44800 | 0.34262 |
| H | –2.95399 | 0.37583 | –0.21325 |
| H | –2.23054 | 2.37671 | –1.27204 |
| H | –4.05785 | 2.63971 | –2.62126 |
| H | –3.66797 | 4.17765 | 0.27089 |
| H | –2.12069 | 2.09062 | 2.80131 |
| H | –0.33745 | 6.06877 | –2.57326 |
| H | 7.38039 | 0.18650 | –3.18372 |
| H | 4.94191 | –1.10998 | –0.19676 |
| H | 4.84752 | –1.78990 | 2.05264 |
| H | 2.34009 | 5.83395 | –0.18232 |
| H | 1.35049 | 6.26136 | –2.43438 |
| H | 8.68037 | 0.12314 | 2.33265 |
| H | –1.31833 | –1.11056 | 4.38802 |
| H | –2.78431 | 0.27759 | –4.55583 |

**Table S91.** Optimized geometry coordinates of C:FFO (W–edge)
Energy: –1617084.1062825 (au)

| | | | |
|---|---|---|---|
| O | 10.33507 | 3.79829 | –0.21237 |
| C | 9.23558 | 4.63436 | 0.08335 |
| C | 7.95380 | 3.85938 | 0.27170 |
| O | 7.63219 | 3.02971 | –0.87715 |
| C | 7.88259 | 2.88437 | 1.42114 |
| O | 7.80722 | 3.54711 | 2.66776 |
| C | 6.62887 | 2.08812 | 1.06798 |
| O | 5.46277 | 2.83004 | 1.37983 |
| C | 6.74000 | 2.00870 | –0.46450 |
| N | 7.21349 | 0.67404 | –0.90153 |
| C | 6.20759 | –0.29106 | –1.03763 |
| O | 5.03317 | 0.04404 | –0.82657 |
| N | 6.52330 | –1.55092 | –1.40199 |
| C | 7.80255 | –1.85407 | –1.61313 |
| N | 8.05769 | –3.11250 | –1.96160 |
| C | 8.86244 | –0.90086 | –1.46461 |
| C | 8.52681 | 0.34875 | –1.10490 |
| N | 2.32925 | –4.11635 | –0.29755 |
| C | 3.21917 | –3.16913 | –0.60010 |
| N | 4.45671 | –3.55572 | –1.32188 |
| N | 3.04737 | –1.86017 | –0.26011 |
| C | 1.94730 | –1.46556 | 0.39519 |
| O | 1.80622 | –0.13010 | 0.71538 |
| C | 0.96806 | –2.42996 | 0.72262 |
| N | –0.22833 | –2.18525 | 1.42465 |
| C | –1.35535 | –2.94112 | 1.18116 |
| C | –1.01677 | –4.37425 | 1.41499 |
| N | 0.17503 | –4.83157 | 0.70474 |

| | | | |
|---|---|---|---|
| C | 1.19657 | –3.77486 | 0.36082 |
| C | –1.79519 | –2.72869 | –0.26615 |
| N | –2.18037 | –1.36046 | –0.62730 |
| C | –6.13267 | –0.33872 | 0.31909 |
| C | –5.33960 | 0.60593 | –0.33485 |
| C | –4.02419 | 0.28318 | –0.65325 |
| C | –3.53002 | –0.97976 | –0.31989 |
| C | –4.33441 | –1.91588 | 0.33189 |
| C | –5.64068 | –1.59578 | 0.63768 |
| C | –7.59821 | –0.06225 | 0.68529 |
| O | –8.06857 | –0.47748 | 1.78642 |
| N | –8.45204 | 0.64169 | –0.24887 |
| C | –9.82127 | 0.82829 | 0.11153 |
| C | –10.27109 | 2.31695 | 0.28003 |
| C | –10.07738 | 3.24859 | –0.84356 |
| C | –8.82700 | 4.04086 | –0.84109 |
| O | –8.81339 | 5.24880 | –0.35150 |
| O | –7.78069 | 3.51927 | –1.31367 |
| C | –10.56887 | 0.15043 | –0.96503 |
| O | –10.27464 | 0.41034 | –2.22059 |
| O | –11.45111 | –0.68122 | –0.67359 |
| C | –0.30269 | –1.16871 | 2.50050 |
| O | –1.29955 | –0.94946 | 3.12758 |
| H | –9.78208 | 2.68873 | 1.18861 |
| H | –11.34048 | 2.27812 | 0.52869 |
| H | –10.16657 | 2.70910 | –1.79578 |
| H | –10.86702 | 4.00230 | –0.82342 |
| H | –9.95092 | 0.33012 | 1.07921 |
| H | 9.08076 | 5.37230 | –0.71908 |
| H | 9.37890 | 5.18962 | 1.02288 |
| H | 7.15591 | 4.60408 | 0.40895 |
| H | 8.76152 | 2.23242 | 1.43200 |
| H | 6.87055 | 3.76648 | 2.79718 |
| H | 6.60745 | 1.09737 | 1.53512 |
| H | 4.69714 | 2.25085 | 1.25141 |
| H | 5.76219 | 2.13896 | –0.92700 |
| H | 8.98172 | –3.39965 | –2.23991 |
| H | 9.89472 | –1.17327 | –1.64524 |
| H | 9.23079 | 1.16708 | –0.99304 |
| H | 5.22115 | –2.88493 | –1.21270 |
| H | 4.66603 | –4.52043 | –1.09841 |
| H | –1.84757 | –5.03425 | 1.13576 |
| H | –0.85923 | –4.52222 | 2.49184 |
| H | –0.95025 | –2.99114 | –0.91131 |
| H | –2.59729 | –3.40715 | –0.55879 |
| H | –1.46933 | –0.65486 | –0.48223 |

| | | | |
|---|---|---|---|
| H | –5.74718 | 1.58045 | –0.58408 |
| H | –3.37672 | 0.99802 | –1.15195 |
| H | –3.96739 | –2.88399 | 0.62826 |
| H | –6.28175 | –2.29408 | 1.16440 |
| H | 0.65554 | –0.64610 | 2.59779 |
| H | 11.14179 | 4.31731 | –0.11499 |
| H | 7.28517 | –3.71132 | –2.21006 |
| H | 3.74221 | –1.13204 | –0.50581 |
| H | 0.07087 | –5.50707 | –0.03979 |
| H | –8.19276 | 0.74016 | –1.22045 |
| H | –11.52985 | –0.82337 | 0.28211 |
| H | –7.07055 | 4.18160 | –1.22348 |
| H | –2.12742 | –2.61708 | 1.88350 |
